# Standing genetic variation and chromosome differences drove rapid ecotype formation in a major malaria mosquito

**DOI:** 10.1101/2022.11.21.517335

**Authors:** Scott T. Small, Carlo Costantini, N’Fale Sagnon, Moussa W. Guelbeogo, Scott J. Emrich, Andrew D. Kern, Michael C. Fontaine, Nora J. Besansky

## Abstract

Species distributed across heterogeneous environments often evolve locally adapted ecotypes, but understanding of the genetic mechanisms involved in their formation and maintenance in the face of gene flow is incomplete. In Burkina Faso, the major African malaria mosquito *Anopheles funestus* comprises two strictly sympatric and morphologically indistinguishable yet karyotypically differentiated forms reported to differ in ecology and behavior. However, knowledge of the genetic basis and environmental determinants of *An. funestus* diversification was impeded by lack of modern genomic resources. Here, we applied deep whole-genome sequencing and analysis to test the hypothesis that these two forms are ecotypes differentially adapted to breeding in natural swamps versus irrigated rice fields. We demonstrate genome-wide differentiation despite extensive microsympatry, synchronicity, and ongoing hybridization. Demographic inference supports a split only ~1,300 years ago, closely following the massive expansion of domesticated African rice cultivation ~1,850 years ago. Regions of highest divergence, concentrated in chromosomal inversions, were under selection during lineage splitting, consistent with local adaptation. The origin of nearly all variation implicated in adaptation, including chromosomal inversions, substantially predates the ecotype split, suggesting that rapid adaptation was fueled mainly by standing genetic variation. Sharp inversion frequency differences likely facilitated adaptive divergence between ecotypes, both by suppressing recombination between opposing chromosomal orientations of the two ecotypes, and by maximizing recombination within the structurally monomorphic rice ecotype. Our results align with growing evidence from diverse taxa that rapid ecological diversification can arise from evolutionarily old structural genetic variants that modify genetic recombination.

**Significance Statement:** Local adaptation to heterogeneous environments is pervasive, but its underlying genetic basis is incompletely understood. Within a major African malaria vector, *An. funestus*, are two chromosomally differentiated groups that are co-localized, morphologically indistinguishable, and reported to differ both in ecology and behavior relevant to malaria transmission and control. Progress in understanding the genetic basis and environmental determinants of vector diversification was impeded by the lack of modern genomic resources. Here we perform deep whole-genome sequencing on individuals from these groups, establishing that they are differentiated genome-wide in a manner consistent with recent ecotype formation associated with the exploitation of a new anthropogenic larval habitat. Such rapid malaria vector diversification was facilitated by standing genetic variation, including evolutionarily old chromosomal rearrangements.

## Introduction

Widespread species inhabiting heterogeneous environments are likely to experience spatially varying selection. This often leads to local adaptation and formation of ecotypes—intraspecific groups distinguished by suites of heritable differences that increase fitness to the local conditions of their home environments (1, 2). Well-characterized examples include the annual and perennial ecotypes of yellow monkeyflowers (3), dune and nondune ecotypes of sunflowers (4), marine and freshwater ecotypes of threespine stickleback fish (5), and forest/prairie ecotypes of deer mice (6). Ecotypic differentiation is of central importance in evolution, as it shapes species diversity and plays potential roles in range expansion, response to climate change, and ecological speciation (7). Ecotype formation also has underappreciated public health significance. Malaria, one of humankind’s deadliest infectious diseases, is transmitted by highly evolvable *Anopheles* mosquito vectors (8) whose effective control may be complicated or even compromised by local adaptations in the vector that diversify the ecological and environmental range (9), and alter epidemiologically relevant behaviors (10).

Although ecotypes are widely observed across a diversity of plant and animal species, the genetic mechanisms that promote and maintain this form of intraspecific structuring of variation has been a longstanding puzzle. Because ecotypes are at least partially interfertile, how do they remain differentiated by a composite of variation in many traits and loci (1) when connected by dispersal and gene flow? Population models and simulations suggest that “genomic islands of divergence” in which locally adapted alleles are clustered into genomic regions with low recombination, may provide one explanation (11). Leading candidates for such genomic regions are chromosomal inversions (12), a form of structural variation in which gene order along a chromosome is reversed. Recombination suppression between opposite orientations in heterozygotes can maintain linkage between locally adapted (13) and/or coadapted (epistatically interacting) (14) alleles at multiple loci, protecting them from homogenizing gene flow with maladapted immigrant alleles (15). Historically, segregating chromosomal inversions were known principally from those species (particularly dipterans like Drosophila) whose chromosome banding patterns were readily visualized cytogenetically. Today, genomics-enabled studies are revealing that inversions are pervasive (16). Most plant and animal inversions are large (~8 Mb on average), spanning tens to hundreds or thousands of genes, and many are relatively ancient, predating ecotypic divergence and even species origins (16). Empirical data have associated inversions with complex traits and environmental variables (4, 6, 15, 17), and have implicated balancing or divergent selection in the longterm maintenance of inversion polymorphism (18). Growing evidence from diverse species—including the aforementioned monkeyflowers (3), sunflowers (4), stickleback (19), and deer mice (6)—reinforces the view that chromosomal inversions are major instruments of ecotypic differentiation.

*Anopheles funestus* bears primary responsibility for highly efficient human malaria transmission across its vast tropical African distribution (20, 21), sharing that role only with three members of the better-characterized *Anopheles gambiae* complex. The *An. funestus* genome harbors extensive amounts of nucleotide diversity that is only weakly structured at the continental level, consistent with large effective population size and high connectivity (22, 23). Genomic structural variation is also abundant, with 17 paracentric chromosomal inversions segregating on the two autosomes (no inversions are known from the X) (24). Following historical tradition in an era with few molecular markers, extensive cytogenetic surveys of *An. funestus* population structure in Burkina Faso, West Africa were begun in the 1990s based on the frequencies of five polymorphic chromosomal inversions (25). Fourteen localities were surveyed along a latitudinal transect crossing the country’s three ecoclimatic zones (Sahel, Sudan–Savanna, and Sudan–Guinea), and several localities were repeatedly sampled over a three-year period (25). Variation in inversion frequency followed no temporal pattern nor any obvious geographic or climatic cline, although the degree of polymorphism appeared to be related to local ecological conditions—proximity to natural swamps (highest inversion frequencies) versus large-scale rice crop areas (lowest frequencies) (25). However, large, temporally stable departures from Hardy-Weinberg equilibrium (HWE) marked by significant heterozygote deficits were found for four of the five inversions (2Ra, 2Rs, 3Ra, and 3Rb) *within* samples collected from individual localities, at nearly all sampling locations (25). An intensive three-year follow-up study conducted five years later focused on two nearby (~1km) Sudan-Savanna villages in central Burkina Faso that surround a natural swamp and rice fields (26). The cytogenetic analysis of >4,600 specimens successfully scored from both villages confirmed a marked and temporally stable departure from HWE due to significant deficits of heterokaryotypes, and uncovered significant linkage disequilibrium among inversion systems on independently assorting chromosomes (26). The possibility of a confounding effect due to the pooling of temporally or spatially distinct subpopulations with different allele frequencies was ruled out, on the basis of a set of subsamples collected at the minimum possible temporal (one week) and spatial (20 meters) scale (26). This pattern of chromosomal variation, incompatible with a single panmictic population, sparked the working hypothesis that populations of *An. funestus* in Burkina Faso consist of two temporally stable, strictly sympatric units of uncertain taxonomic status, sharing a common set of inversions while exhibiting contrasting degrees of polymorphism (26). These units, designated as ‘Kiribina’ and ‘Folonzo’ chromosomal forms, show restrictions to gene flow interpreted by the authors as an ongoing incipient speciation process promoted by positive assortative mating (26), although ecologically-dependent postmating isolation was not excluded. On the basis of available evidence (25), alternative explanations such as hybridization between intergrading allopatric populations (which fit the data from *An. funestus* populations in Senegal and Cameroon) (27–32) appear less likely in Burkina Faso (26).

Application of deterministic algorithms to assign chromosomally-scored *An. funestus* specimens to chromosomal form restored HWE and linkage equilibria for all inversion systems within each form (25, 26). Folonzo was hypothesized to belong to the same taxon as *An. funestus* from eastern and southern Africa, based on its high level of inversion polymorphism on chromosomes 2R and 3R, and its association with species-characteristic larval habitats—natural swamps and ditches with abundant aquatic vegetation (25). Kiribina was considered a new taxon that was nearly depauperate of inversion polymorphisms, carrying mainly the standard chromosomal orientations, and was found in association with large-scale rice cultivation. The working hypothesis of differences in larval ecology between the forms was reinforced by intensive longitudinal studies of adults (polytene chromosome analyses are possible only at the adult stage). These revealed seasonal variation in the relative frequency of the two forms, confirmed across six successive years (33). Only Folonzo abundance was correlated with climatic variables related to temperature and rainfall, as expected if the growth of aquatic vegetation is rainfall dependent in its natural larval habitats, but not the irrigated rice field sites of Kiribina (33). Behavioral and bionomic studies demonstrated that although their human biting behavior and malaria parasite infection rates were statistically indistinguishable and comparably high—indicating that both Kiribina and Folonzo are formidable malaria vectors—there were important epidemiologically relevant differences (10). In particular, Folonzo was significantly over-represented in indoor-resting collections and showed stronger post-prandial endophily, while Kiribina predominated outdoors (10). This suggests that the two taxa may not be uniformly exposed to indoor-based vector control interventions. Mitochondrial and microsatellite markers outside of inversions indicated weak but significant differentiation between the groups (34). However, the combined absence of a molecular diagnostic assay and modern genomic tools (a high-quality reference genome and low-cost genomic re-sequencing) challenged further elucidation of this system, not only at the genomic level but also at the level of larval field studies necessary to identify locally adapted traits and ecological forces driving selection.

Here, we leverage the recent chromosome-scale *An. funestus* assembly (35) and apply individual, wholegenome deep sequencing to investigate retrospective collections of Kiribina and Folonzo from Burkina Faso. We sought to establish the extent to which inversion-based differentiation extends to collinear genomic regions, and to explore whether the timing and pattern of genetic differentiation is consistent with local adaptation by Kiribina to breeding in rice fields. Our data conclusively demonstrate that Kiribina and Folonzo are differentiated genome-wide despite extensive sympatry and synchronicity. Because the genetic lineages are not fully congruent with the karyotype-based classifications, we follow precedent (36) in renaming these taxa ‘K’ and ‘F’ forms of *An. funestus*. Remarkably, the recent K-F split follows African rice domestication and the massive expansion of its cultivation ~1,850 years ago (37), suggesting that this newly abundant larval habitat may have been a driver of K diversification. As expected for local adaptation, the genomic landscape of differentiation is heterogenous, and we infer that the regions of highest differentiation were under selection during lineage splitting. The origin of nearly all variation implicated in local adaptation, including chromosomal inversions, substantially predates separation of the two groups, suggesting that rapid exploitation of a novel anthropogenic habitat was fueled mainly by standing genetic variation and not *de novo* point or structural mutations. Structural homozygosity for the standard orientation of chromosomal inversions in K likely promoted adaptive divergence, by maximizing recombination within K and suppressing it between K and F, which carries the opposite chromosomal orientations at high frequencies.

## Results and Discussion

### Genetic structure reveals two differentiated groups of *An. funestus* in Burkina Faso

We analyzed 168 *An. funestus* sampled from 11 villages spanning a ~300-km east–west transect of Burkina Faso, where Kiribina and Folonzo co-occur at the village level (Fig 1A; *SI Appendix*, Text, Fig. S1, Tables S1-S2). Individual whole genome sequencing yielded a mean coverage of 38X after filtering (*SI Appendix*, Text, Tables S3-S4). Following read mapping to the AfunF3 reference and further filtering (*SI Appendix*, Text), we identified 24,283,293 high quality single nucleotide polymorphisms (SNPs) among 114,270,430 accessible sites scored across all individuals. Using a subset of these SNPs, we estimated genetic population structure based on principal components analysis (PCA) and the individual ancestry-based algorithm implemented in ALStructure (38) (*SI Appendix*, Text).

**Figure 1.**
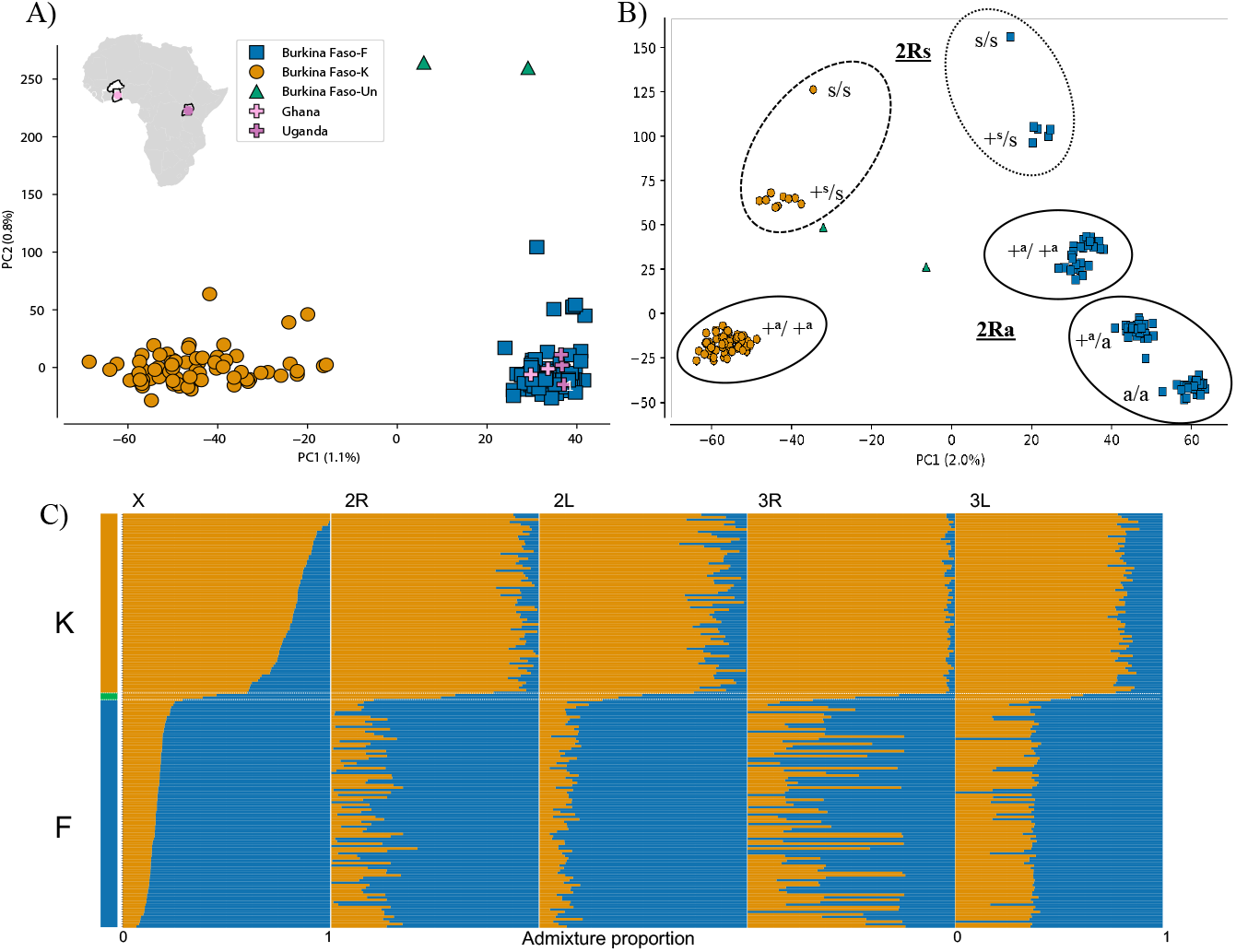
Genetic structure of *An. funestus* mosquitoes sampled from Burkina Faso. A) Plot of the first two principal components from the X chromosome PCA that included *An. funestus* reference samples from Ghana and Uganda. Orange circles and blue squares correspond to Burkina Faso genotypes assigned to ecotypes K and F, respectively. Green triangles represent two unclustered Burkina Faso genotypes that were unassigned (‘Burkina Faso-Un’). B) Plot of first two principal components from arm 2R with inversion 2Ra (solid-line) and 2Rs (dashed-line). C) Estimated individual ancestry proportions for Burkina Faso *An. funestus*. Y-axis color bar at left reflects individual assignments based on the X chromosome PCA (K, orange; F, blue; unassigned, green). Individuals are represented as thin horizontal lines, grouped by assignment and displayed across the plot in descending order of ancestry proportion to K for the X chromosome. Different chromosome arms (X, 2R, 2L, 3R, 3L), designated by columns across the top of the plot, are separated by white lines.

To avoid the potentially confounding effects of autosomal inversion polymorphisms, our initial inference by PCA relied on the inversion-free X chromosome. Given the pan-African distribution of *An. funestus*, previously published genomic variation data from Ghana and Uganda (23) were included to provide broader context (*SI Appendix*, Table S1). The analysis revealed two main clusters (Fig 1A). One contains all 70 of the Burkina Faso *An. funestus* that would have been classified as Folonzo by inversion karyotyping using the deterministic algorithm of Guelbeogo, *et al*. (26), as well as another 22 Burkina Faso *An. funestus* that would have been (mis)classified as Kiribina due to low inversion polymorphism (blue squares, Fig 1A; *SI Appendix*, Table S1) [See (26) for a discussion of deterministic algorithms and misclassification]. This same cluster also includes the geographically distant samples from both Ghana and Uganda (crosses, Fig 1A). The second cluster corresponds to 74 Burkina Faso *An. funestus* that would have been classified as Kiribina (26) (orange circles, Fig 1A). Only two Burkina Faso *An. funestus* mosquitoes were not included in either of the two clusters; these were considered unassigned (green triangles, Fig 1A).

PCA based on SNPs from individual autosome arms showed exactly the same two clusters and unassigned mosquitoes (*SI Appendix*, Fig S2). However, the pattern within clusters for autosome arms carrying polymorphic inversions was more complex, as expected, reflecting the inversion karyotype of the mosquito carriers (*SI Appendix*, Fig S3). Strikingly, mosquito carriers of inversion karyotypes that segregate in both populations (*e.g*., homokaryotypes for 3R+^a^, 3R+^b^, 2R+^a^, 2R+^s^, and 2Rs) cluster by population and not by inversion karyotype. There was no discernable contribution of geographic sampling location to the clustering patterns for any chromosome arm (*SI Appendix*, Fig S4).

Individual-based ancestry analysis of the Burkina Faso sample supported the existence of the same two genetically differentiated populations as those identified by PCA (Fig 1B). Notably, the two unassigned mosquitoes have admixture proportions suggestive of inter-population hybrids (rows enclosed by horizontal white lines in Fig 1B). Compelling evidence supporting this suggestion is presented below.

Taken together, these results are indicative of the existence of two genetically differentiated ecotypes of *An. funestus* in Burkina Faso. They closely correspond to the formerly recognized chromosomal forms Kiribina and Folonzo, but the correspondence is imperfect because the deterministic classification system misclassifies a fraction of karyotypes that are shared between ecotypes (in particular, the standard karyotype characteristic of, but not exclusive to, Kiribina; *SI Appendix*, Table S1). Following precedent (36), we distinguish ‘molecularly defined’ forms from the prior chromosomal forms with the new designations K and F. The fact that the F ecotype from Burkina Faso clusters together with other *An. funestus* sampled from neighboring Ghana and distant Uganda (Fig 1A) is consistent with the hypothesis that F is genetically allied with *An. funestus* populations across tropical Africa, while K has a more recent and local origin (25).

### Demographic inference of a K-F split 1,300 years ago correlates with expanded rice agriculture

To explore the history of K-F divergence, we used an approximate Bayesian computation (ABC) method (39) to compare five alternative demographic scenarios, involving different amounts and timing of gene flow over four epochs (*SI Appendix*, Text, Fig S5). Model selection in this statistical framework (*SI Appendix*, Text, Tables S5-S8) allowed us to reject histories of panmixia, allopatric isolation, and secondary contact. The most strongly supported scenario, chosen as the most likely, involved isolation with initial migration (IIM). However, its posterior probability (0.44) was only slightly higher than a model of isolation with migration (0.40), reflecting the difficulty of distinguishing these two closely related models (*SI Appendix*, Table S7).

We estimated demographic parameters under the chosen IIM model using two independent methodologies, ABC and an alternative approach based on Generative Adversarial Networks (GAN) implemented in the software *pg-gan* (40). The latter uses real data to adaptively learn parameters capable of generating simulated data indistinguishable from real data by machine learning (40). It operates directly on genotype matrices and avoids reducing genotypes to summary statistics, unlike ABC. Importantly, genetic data simulated under the IIM model by *pg-gan* and ABC produced similar summary statistics (*SI Appendix*, Text, Table S8). If the chosen IIM model closely approximates the true demographic history, we expect that the observed data and data simulated under the IIM model should have comparable summary statistics. To verify this expectation, posterior predictive checking was performed by simulating 10,000 new datasets with population parameter values drawn from the posterior distributions of the population parameters estimated from the retained *pg-gan* runs. Summary statistics were calculated using noncoding regions only, under the assumption that noncoding regions are less affected by selection. The median of each observed summary statistic was compared with the distribution of the 10,000 simulated test statistics. We calculated the percentile for each median value within the distribution of the simulated data statistic. All observed medians fell within the 17% – 84% percentiles for K, and the 20% – 59% percentiles for F, providing confidence in the IIM model.

Under the IIM model, we estimated that the ancestral K-F lineage split from an outgroup population of *An. funestus* ~6,000 years before present, consistent with our previous inference of a range expansion of *An. funestus* to its present continent-wide distribution less than 13,000 years ago (23). The relatively steep decline of the ancestral K-F lineage to an N_e_ of ~11,000 about 2,000 years ago (Fig 2A) may reflect widespread unfavorable environmental conditions connected with the drying of the Sahara (41). Parallel climate-driven population reduction of wild African rice observed at the continental scale, beginning more than 10,000 years ago and reaching a minimum at ~3,400 years ago, is the hypothesized trigger for African rice domestication, which may have originated in the inner Niger delta of West Africa (37) (but see ref 42). By ~1,850 years ago, there was a strong expansion of domesticated rice cultivation (37). Strikingly, we estimate that K and F split ~1,300 (credible interval ~800-1,800) years ago, which fits well with the timing of a newly abundant anthropogenic larval habitat (Fig 2A, *SI Appendix*, Table S8). Around the time of K-F splitting, we inferred an ancestral N_e_ of ~64,000. Since K-F divergence, F expanded to an N_e_ of ~3,250,000 while K maintained a steady population size (Fig 2A, *SI Appendix*, Table S8). Postdivergence K-F gene flow, initially mainly from F to K, slowed ~500 years ago and effectively ceased by ~100 years ago (Fig 2B).

**Figure 2.**
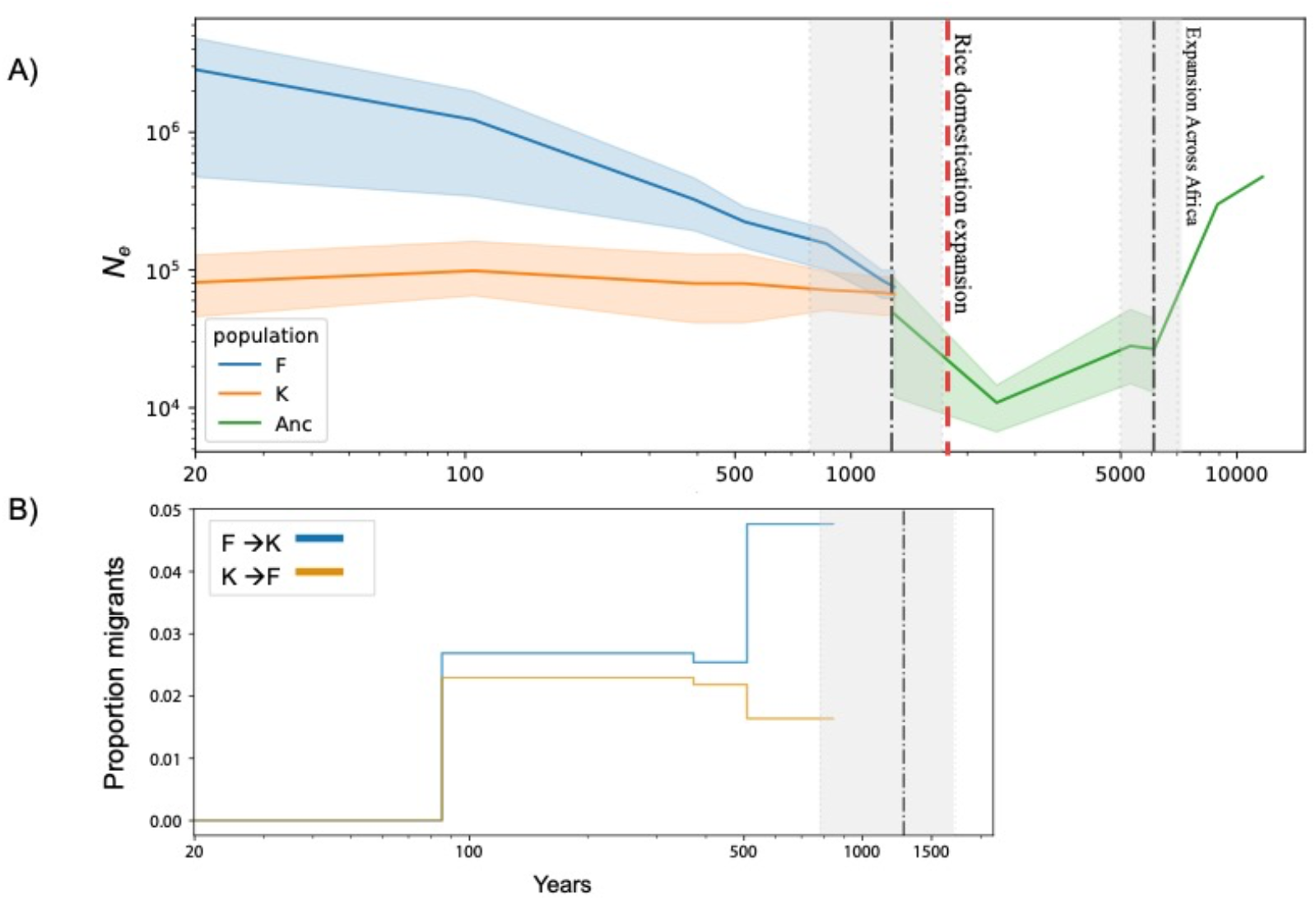
Demographic history inferred with ABC for *An. funestus* ecotypes K and F. A) Effective population size (N_e_) trajectories and population split times in years before present, assuming 11 generations/year. Median N_e_ values and 95% credible intervals are plotted as colored lines and corresponding shading. Median split times and 95% credible intervals are represented as dash-dotted black vertical lines with gray shading. The dashed red vertical line represents the proposed time (~1,850 years) of a strong domesticated rice expansion in the Niger River delta (33). B) K-F migration rates through time in years before present, subsequent to their split (dash-dotted black vertical line). The proportion of migrants declined to 0 in both directions at ~100 years.

A recent advance in genome-wide genealogy estimation, implemented in a new method called *‘Relate’*, allows inference of genealogical trees across all loci (haplotypes) in a large sample of genomes (43). Using these genealogies directly, *Relate* infers recombination, mutational ages, and—based on the tracking of mutation frequencies through time—identifies loci under positive natural selection (see below). Applying *Relate*, we find that the first K-F cross-coalescent events, estimated from autosome arm genealogies, occurred with a mean timing of ~100 years ago (*SI Appendix*, Text, Fig S6), providing further evidence of the recent cessation of realized gene flow between K and F. Note that the most recent cross-coalescence is older on the X chromosome than for the autosomes (*SI Appendix*, Fig S6), but whether this result is due to smaller effective population size of the X, demography, and/or resistance to gene flow on the X—as seen, *e.g.*, in humans (44) and *An. gambiae* (45)—is unclear.

### Genetic diversity and differentiation along the genome are heterogeneous between ecotypes

Overall, nucleotide diversity (π) is comparable between K and F (genome-wide mean values of 0.603E-2 and 0.624E-2, respectively). The same was true in windows along the genome, except around chromosomal rearrangements (Fig 3). Inversions 2Ra, 3Ra and 3Rb segregate exclusively in the F sample at frequencies of 0.44, 0.73, and 0.40, respectively, while K carries only the corresponding standard orientation of those inversions. [Inversions 2Rs and 3La are not considered here, as the former is at low frequency in our samples of K (0.07) and F (0.04), and the latter segregates only in F at a frequency of 0.05 (*SI Appendix*, Table S1)]. Inversions 2Ra, 3Ra, and 3Rb are large [8 Mb, 9.4 Mb, and 12.5 Mb, respectively (46)], collectively spanning 14% of the AfunF3 reference genome. Nucleotide diversity appears slightly elevated at these rearrangements in F relative to K (Fig 3), particularly for inversions 2Ra and 3Rb whose frequencies are more intermediate than 3Ra. In addition, values of Tajima’s *D* are less negative across the genome in K relative to F (genome-wide means, −2.031 and −2.408, respectively), likely owing to the rapid post-divergence population size expansion of F and the lower N_e_ in K (possibly involving a bottleneck at the time of the split) (Fig 2A).

**Figure 3.**
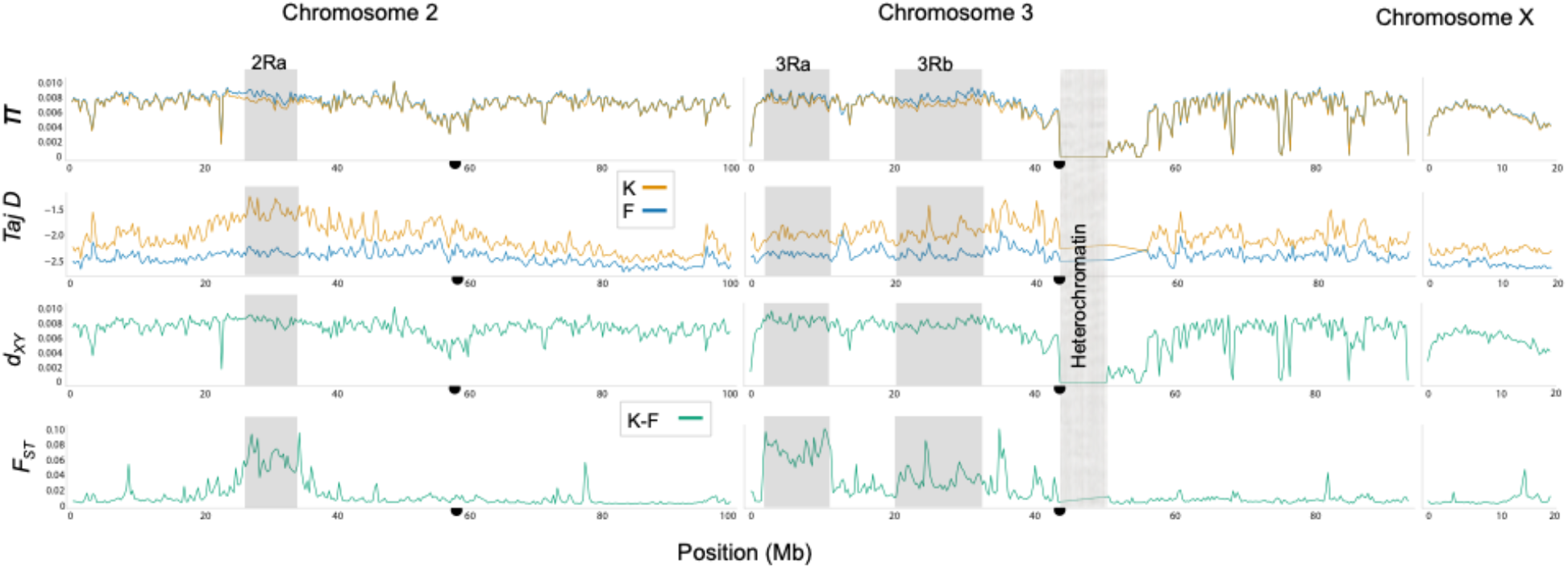
Whole-genome scans of diversity and divergence for K and F. Each statistic was calculated in 10-kb non-overlapping windows and smoothed with a moving average over 10 windows for plotting. From top to bottom, panels display mean values of nucleotide diversity (*π*), Tajima’s *D* (Taj D), absolute divergence (*d_XY_*), and differentiation (*F_ST_*) at genomic positions (in millions of bases, Mb) along chromosomes 2, 3, and X. Rearranged and heterochromatic (no data) regions are shaded and labeled. Centromeres are indicated as black half circles.

We examined the genomic landscape of divergence using both absolute (d_XY_) and relative (F_ST_) measures, recognizing that relative measures of divergence may reflect reduced diversity instead of reduced gene flow (47). We found that the pattern of absolute K-F divergence closely corresponds to the pattern of nucleotide diversity in the ecotypes, probably owing to their recent split, and differs strikingly from the pattern of relative K-F divergence along the chromosomes (Fig 3). The relative measure clearly takes on much larger values in the rearranged versus collinear chromosomal regions (Fig 3). Indeed, the mean F_ST_ value for rearranged regions (0.045) is fourfold higher than for collinear regions (0.011), consistent with relatively high inversion frequencies in F and homozygosity for the standard arrangements in K. Although the corresponding d_XY_ values for these regions follow the same trend, being significantly larger for rearranged versus collinear regions (0.769E-2 and 0.620E-2; K-S test p-value < 9.56E-173), d_XY_ has less power to detect differences among loci due to the very recent K-F split, as new mutations must arise for d_XY_ to increase, while F_ST_ only requires changes in allele frequency (47). Genomic regions of high F_ST_ are most noticeable in rearranged regions, but additional peaks of F_ST_ also were identified in collinear regions, such as position ~13.8 Mb on the X chromosome (Fig 3).

### Ancient standing variation and chromosome differences fueled rapid ecotype formation

To provide insight into the genomic underpinnings of local adaptation, we identified candidate targets of differential selection or linked selection using a two-step process. We began with an empirical outlier approach to identify exceptionally diverged genomic regions, under the expectation that loci responsible for local adaptation should be more diverged than neutral loci. Considering separately the collinear and rearranged genomic partitions due to large differences in mean F_ST_ between them, we computed average normalized F_ST_ in 10-kb windows and found a total of 44 outlier windows, or 26 outlier blocks when consecutive windows were combined (*SI Appendix*, Text, Table S9). Because neutral loci can vary widely in levels of differentiation for reasons other than selection, we also used *Relate* (43) to infer genome-wide genealogies [1,484,174 genealogies with a mean length of 133 bp (SD = 1,337 bp)] and to identify those under positive selection. To arrive at a list of candidate targets of differential selection, we found the intersection between the 44 outlier F_ST_ windows and the SNPs from 40,453 genealogies inferred to be under selection in K and/or F. At this intersection were 37 outlier F_ST_ windows arranged in 22 genomic blocks, spanning 202 genealogies involving at least one SNP inferred to be under differential selection (262 SNPs in total; *SI Appendix*, Text, Figs S7-S8, Tables S9-S10). As candidate SNPs from the same genealogy are linked, we enumerate genealogies as targets of selection rather than component SNPs, and for simplicity, refer to them as ‘loci’, except where noted. We predicted that if these candidate loci contribute to local adaptation, they should show no evidence of migration between K and F. To test this prediction, we used a supervised machine learning approach implemented in the program FILET (Finding Introgressed Loci using Extra Trees Classifiers) (48) to identify 10 kb windows along the genome with a high probability of no-migration (*SI Appendix*, Text). All 37 outlier F_ST_ windows were classified with high confidence as no-migration windows (*SI Appendix*, Tables S9, S11), consistent with our expectation.

Under the hypothesized scenario that K recently adapted to exploit a novel habitat for growth and development of its immature stages, we expect asymmetries between ecotypes in the number of candidate loci and the timing when selection was first inferred by *Relate*. In particular, we predict an excess of candidate loci, and a more recent onset of selection, in K versus F. Consistent with these predictions, we detect more candidate loci in K: 123 compared to only 79 in F (with 5 loci shared between ecotypes) (*SI Appendix*, Table S10). Furthermore, considering the set of individual SNPs inferred to be under differential or linked selection in K and F, the timing of first selection is more recent for K (Kruskal-Wallis p-value=0.00015). Shown in Fig 4 as overlapping histograms along the right-hand Y-axis, the mean time in years before present was 1,867 for K (0.05–0.95 quantile, 243–4,708) and 2,283 for F (1,192–4,708) (see also *SI Appendix*, Fig S9). These histograms suggest that K experienced more selection than F following their split ~1,300 years ago (indicated in Fig 4 as a horizontal black line against the right-hand Y-axis). Indeed, 107 SNPs in K versus 15 in F were inferred to have come under selection since that time (see *Data Availability*).

**Figure 4.**
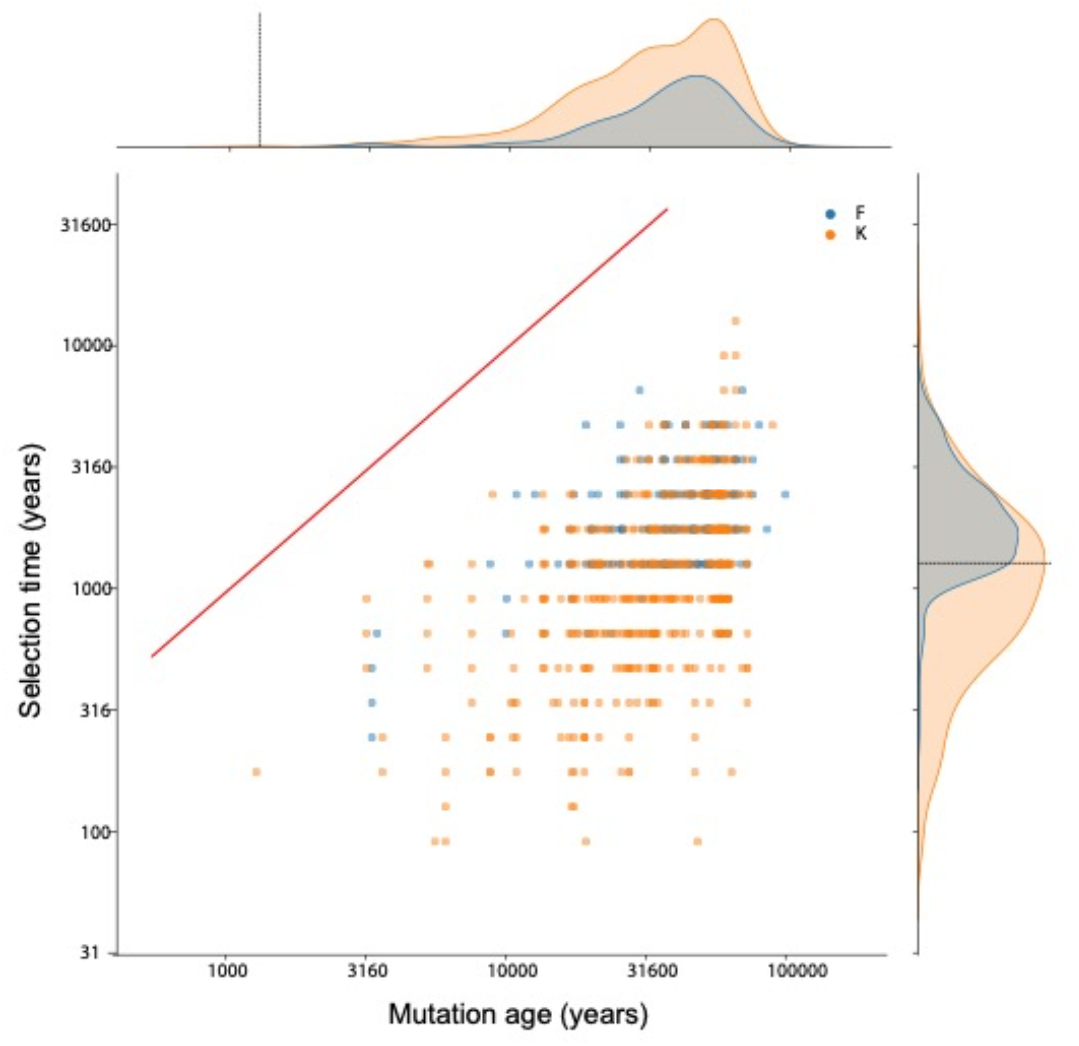
Age of mutation versus time at onset of selection. In the central panel, SNPs under selection in K (orange) or F (blue) are plotted as dots according to their estimated age (X axis, in years) against the inferred time of first selection (Y axis, in years), as inferred by *Relate* (39). The red line, included for reference, indicates a hypothetical 1:1 correspondence between mutation age and onset of selection. Time in years assumes a generation time of 11 per year. Against the top-most X axis and the right-hand Y axis and are the corresponding densities of mutation ages and selection times, respectively. Black lines on the density plots mark the split time of K and F.

The recency and rapidity of local adaptation to a new habitat in K is difficult to reconcile with new mutation alone, because many traits involved in adaptation to local environments are expected to be highly polygenic (49) and it is unlikely that all alleles involved are newly mutated. To gain insight into the origin of selected variation in K and F, we focused on the same set of individual SNPs inferred to be under differential or linked selection, and estimated their frequency at the earliest detected onset of selection using *Relate*. We found that most selected variants (89% in K; 93% in F) were segregating at frequencies >5% at this time, implying that selection must have arisen from standing genetic variation (*SI Appendix*, Fig S10). To investigate further, we estimated the mutation age of each variant (*SI Appendix*, Text). Shown in Fig 4 as overlapping histograms along the upper X-axis, the mean mutation age in years before present was 37,058 for K (0.05–0.95 quantile, 9,240–61,993) and 40,687 for F (12,153–67,594). These mutation ages not only greatly predate the K-F split, they overlap the split of *An. funestus* from its sister species, *An. funestus-like* (~38,000 years ago; 95% CI, 31,000 – 45,000 years ago) (23). Taken together, Fig 4 and Fig S10 (*SI Appendix*) indicate that the candidate loci implicated in local adaptation were present as standing variation for substantial lengths of time and at low to intermediate frequencies prior to the onset of selection, factors that could have facilitated the rapid adaptation by K to a human-modified habitat (50).

Both the *Relate* method and our simulations under the chosen IIM model of K-F divergence suggest that there was migration between these ecotypes until the last ~100 years. We explored the genetic architecture of candidate loci in the K ecotype, to test the hypothesis that chromosomal inversions help maintain divergence through suppressed recombination despite gene flow. Although K and F are not distinguished by fixed differences between alternative chromosomal arrangements, three inversions on chromosome arms 2R and 3R segregate in F at high frequencies but are absent (or undetectable due to very low frequency) in K. These inversions (2Ra, 3Ra, and 3Rb) did not arise *de novo* near the time of the K-F split, as our estimates of inversion age suggest that all three are relatively ancient, well-predating that split and approaching or exceeding the age of the split between *An. funestus* and its sister species *An. funestus-like* >30,000 years ago (*SI Appendix*, Text, Fig S11). Nevertheless, three observations implicate inversion differences in maintaining K-F divergence. First, whether considering the set of candidate loci or the set of individual candidate SNPs in K and F under differential or linked selection (*SI Appendix*, Table S10), the number that map to rearranged versus collinear genomic regions is significantly greater in K (Pearson Chi-square p<0.00047; *SI Appendix*, Fig S12). Second, considering K alone, 5,483 of 28,749 (19%) selected loci across the genome map to genomic regions rearranged between ecotypes (red dot in Fig 5A). Drawing 1000 random samples of equal size (5,483) from putatively neutral SNPs across the genome to create a ‘null’ distribution, the fraction of neutral SNPs in K mapping to rearranged regions did not overlap the fraction of candidate loci that did so from any one the draws (box plot in Fig 5A), suggesting that within K there is a significant enrichment of candidate loci inside regions that are rearranged between ecotypes. The third observation exploits the fact that the reduction in recombination in genomic regions rearranged between ecotypes is expected only between chromosomes with opposite orientations. Although inversions segregate at high frequency in the F ecotype, the standard configuration is present (*SI Appendix*, Table S1, Fig S13), and can recombine with the corresponding standard configuration in K. We considered the set of 168 candidate SNPs under differential selection in K, and compared their frequencies in K to the frequency of the same SNPs in F, after partitioning F into two groups by karyotype: homozygous standard (denoted F_S_) and all remaining F (*i.e*., those carrying at least one chromosomal inversion, denoted F_I_). Fig 5B suggests that recombination suppression between opposite inversion configurations of F and K preserves higher K-F differentiation in rearranged genomic regions, despite the potential for gene flow between the shared standard orientation in these rearranged regions.

**Figure 5.**
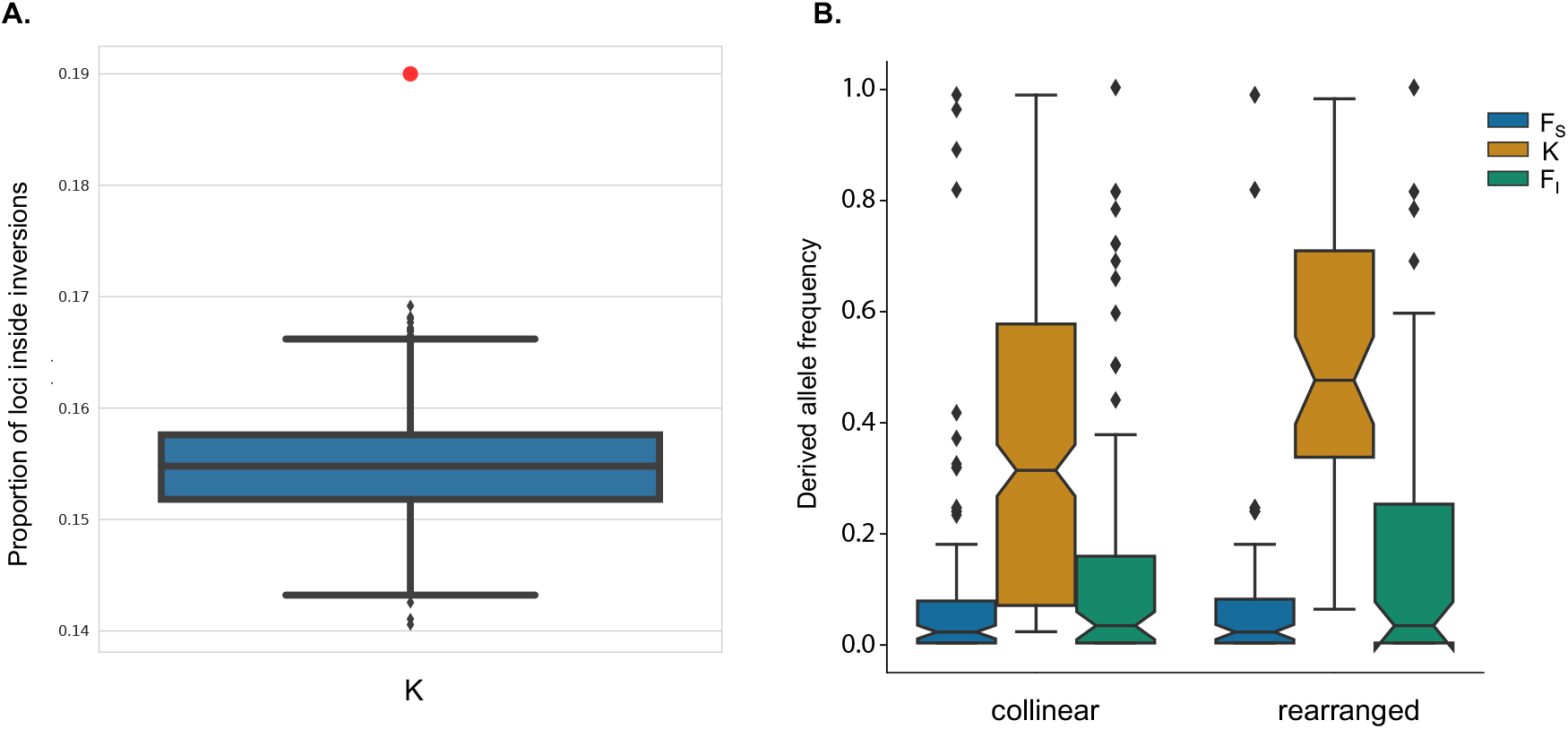
Selected loci in K are enriched within and protected by chromosomal rearrangements. A) Proportion of SNPs inside inversions. The red dot represents the observed fraction of candidate loci in the K genome that map inside rearranged regions. The boxplot beneath represents the percentage in 1,000 random sample sets of the same number of putatively neutral SNPs. B) Boxplot of allele frequencies for individual candidate SNPs in ecotype K, and their corresponding frequencies in F, in collinear versus rearranged genomic compartments. Ecotype F was partitioned by genotype, where F_S_ represents mosquitoes carrying the homozygous standard karyotype with respect to all three focal inversions (2Ra, 3Ra, 3Rb) and F_I_ represents all other karyotypic combinations. Boxplots whose notches do not overlap another boxplot are considered significantly different.

The set of candidate SNPs under differential selection (or linked selection) in the ecotypes map to 25 annotated genes, of which 9 genes are implicated in both, 13 are unique to K, and 3 are unique to F (*SI Appendix*, Table S10). Highly incomplete functional annotation of the *An. funestus* FUMOZ reference renders functional inference particularly fraught. Using orthology to the better-annotated *Drosophila melanogaster* genes in Flybase (flybase.org) where possible, we attribute putative functions to 15 genes (*SI Appendix*, Table S10). In broad categories, these include pattern formation during development (e.g., Wnt and Egfr signaling), metabolism (gluconeogenesis, steroid and neurotransmitter biosynthesis), reproduction (male fertility, sperm storage), and nervous system function (phospholipase C-based and GPCR signaling). Of special note in the last category are two diacylglycerol kinase genes on chromosomes 2 and X. The X-linked gene, an ortholog of *D. melanogaster retinal degeneration A*, has been implicated in pleiotropic functions ranging from visual, auditory, and olfactory signal processing to starvation resistance and ecological adaptation in *Drosophila* and *An. gambiae* (51–54). As the ortholog of this gene in *An. gambiae* was recently identified as the target of a selective sweep in populations from Uganda (55), further investigation may be warranted despite the significant challenges of definitively linking candidate targets of natural selection to adaptive consequences (56).

### Prospects for a molecular diagnostic tool for ecotype identification

To our knowledge, formal morphometric analyses have not been conducted on the K and F ecotypes in Burkina Faso. However, for all practical purposes, K and F are morphologically indistinguishable, and we have shown here that chromosome inversion-based identification tools are imprecise at best. Both future evolutionary ecology studies, and vector surveillance for malaria epidemiology and control, will critically depend upon molecular diagnostics, which are lacking.

Motivated by this goal, we began by assembling a complete ribosomal DNA (rDNA) gene unit by leveraging the long single molecule (PacBio) database developed during construction of the *An. funestus* FUMOZ AfunF3 reference genome (35) (*SI Appendix*, Text). Noncoding spacer regions of rDNA have historically proven fruitful targets for identifying fixed SNP differences between closely related anopheline sibling species (57), owing to the uniquely rapid evolutionary dynamics of the tandemly arrayed rDNA genes. Despite near-complete success including a substantial amount of intergenic spacer upstream and downstream of the transcription unit (*SI Appendix*, Table S12; GenBank accession OP870452), alignment of genomic sequences from K and F to the rDNA assembly yielded no evident fixed differences (*SI Appendix*, Fig S14), consistent with their early stage of divergence.

As an alternative, we screened the genomic sequence data for fixed differences between K and F, using variation (VCF) files compiled separately for K and F samples. This resulted in a total of seven fixed SNP differences, all of which mapped to intronic positions of a single gene (the GPCR AFUN019981) on chromosome 2R, spanning ~15 kb (*SI Appendix*, Fig S15; for alignment, see *Data Availability*). The 5’-end of the predicted mRNA lies only ~2.5 kb outside the proximal breakpoint of inversion 2Ra, a location that is likely protected from gene flux (recombination and gene conversion) between the inverted and standard orientations, potentially explaining—at least in part—the heightened differentiation. As four of the fixed SNP differences between K and F are separated by only 600 bp, it appears feasible that a simple PCR amplicon strategy could be developed. How widely applicable this result might be remains to be investigated, and the inability to detect additional fixed differences across the accessible genome despite deep sequencing is remarkable in itself. It is noteworthy that the two putative K-F hybrids revealed by individual ancestry analysis (Fig 1B) are detectably heterozygous at four or five of the six diagnostic positions that could be unambiguously genotyped, consistent with contemporary K-F hybridization.

### Role of inversions in ecotypic differentiation

The detection of two likely K-F hybrids is indicative of ongoing hybridization. That K and F remain diverged in the face of gene flow suggests that the linkage of allelic combinations adapted to alternative local environments is protected, by selection and reduced recombination countering gene flow in rearranged genomic regions, where the majority of differentially selected candidate loci map in the K ecotype. This suggestion is supported by growing numbers of theoretical and empirical studies in other systems, showing that chromosomal inversions aid divergence with gene flow at different phases along the speciation continuum, from ecotypes to species, across diverse plants, animals and fungi (4, 15, 58). In *An. funestus*, multiple chromosomal inversions are implicated in ecotype differentiation, in particular 2Ra, 3Ra, and 3Rb. Each is large (≥ 8 Mb), with opposite orientations showing elevated F_ST_ even within the F ecotype where they segregate at high frequency (*SI Appendix*, Fig S13). Theory and empirical studies have shown that any one large inversion may span many adaptive loci and control multiple adaptive phenotypes along different environmental axes. That all three inversions seem to play a role implies that they may contribute additively (59) or even epistatically to phenotypes that confer local adaptation. Evidence is still lacking in *An. funestus*, but a systems genetic study of two widespread and adaptive inversion polymorphisms on different arms of chromosome 2 in the congener *An. gambiae* uncovered epistatic interactions between inversions, affecting the expression of hundreds of genes (54).

Circumstantial evidence from *An. funestus* suggests that inversions 2Ra, 3Ra, and 3Rb are selected (17, 32), and in combination, their selective advantage may be strong even if the effect size of individual loci inside inversions is small. Each of the three inversions is found across the entire tropical African range (21), and at a macrogeographic scale, all three inversion frequencies are environmentally structured, supporting a role in ecoclimatic adaptation (17). In Cameroon, the frequency of inversion 3Ra shows a dramatic latitudinal pattern in which it is essentially fixed for opposite orientations between allopatric *An. funestus* populations from the southern rainforest and northern savanna ecological zones (32). Heterozygotes for 3Ra are found mainly in the highlands that separate the two zones, where northern and southern populations intergrade (32). Population genetic modeling of these data supports strong local adaptation, with 3Ra predicted to impact both viability and assortative mating under the assumptions of the models (32).

These lines of evidence, together with the new data presented here, prompt a speculative working model for the role of inversions in *An. funestus* ecotype formation, to be tested by future field work and modeling. We assume that F is part of the widespread population system of *An. funestus* found across tropical Africa (25). This conjecture has little empirical support at present, because available genomic variation data from natural populations of *An. funestus* remains quite limited. The PCA plot in Fig 1A showing complete overlap between F and *An. funestus* population samples from Ghana and Uganda is consistent with our assumption, as are estimates of mean divergence (F_ST_) between the Uganda sample and either K or F, calculated from genomic variation data. These values are always higher in the contrast involving K, regardless of chromosome arm or collinearity [*e.g*., mean F_ST_ for the collinear X is 0.020 (Uganda-F) versus 0.027 (Uganda-K); mean F_ST_ for the 3Ra rearrangement is 0.019 (Uganda-F) versus 0.100 (Uganda-K)]. We propose that inversion polymorphism is maintained in *An. funestus* by spatially divergent selection in the ecoclimatically diverse environments comprising its range, because opposite orientations confer fitness advantages in alternative environments. It should be noted that inversion heterozygotes are common in natural populations of *An. funestus*, and their frequencies are broadly consistent with Hardy-Weinberg expectations within localities. Importantly, the inverted orientations of the three rearrangements (2Ra, 3Ra, 3Rb) segregate at relatively high frequencies in the F ecotype, while K is monomorphic for the standard orientations (2R+^a^, 3R+^a^, 3R+^b^). Structurally monomorphic K populations likely reflect the sieving of ancestral balanced polymorphism during local adaptation to a novel and potentially suboptimal environment (presumed to be large scale rice cultivation), with sieving resulting either from random genetic drift or from selection favoring the fixation of the standard orientations (60). Regardless, in the K-F system, the relatively high frequencies of the inverted orientations in F, versus only the opposing orientations in sympatric K populations, may have facilitated ecotype formation both by suppressing inter-ecotype gene flow in rearranged genomic regions, and by allowing free recombination within K. Free recombination speeds adaptation by allowing natural selection to more efficiently sort beneficial from deleterious mutations (61).

### Role of standing genetic variation in ecotypic differentiation

Our demographic inference suggests a remarkably recent split between K and F ecotypes, approximately 1,300 years ago. Such a rapid time-frame is by no means unprecedented, as the well-studied case of host shifting in *Rhagoletis pomonella* from downy hawthorn to domesticated apple is as recent as 150 years ago (62). However, it begs the question of how ecotypic differentiation, which is expected to involve allelic variants at multiple loci underlying polygenic traits, could arise so quickly on the basis of new mutation alone. The pattern that is emerging from a growing empirical database is that many of the adaptive alleles as well as adaptive genomic architectures (*e.g*., chromosomal inversions) implicated in ecotype formation are considerably older than the ecotypes themselves (4–6). Rather than the lengthy waiting time required for a plethora of new adaptive point and structural mutations, local adaptation and ecological speciation can be jump-started by relatively ancient haplotypes maintained in ancestral populations and/or newly introduced into them through hybridization with other taxa (50, 58). Chromosomal inversions whose origin predates the radiation of the *An. gambiae* species complex were inferred to have been introduced into the primary malaria vector *An. gambiae* through introgressive hybridization with its arid-adapted sibling species *An. arabiensis*, likely fueling the range expansion of *An. gambiae* into arid parts of tropical Africa (9, 45, 63). In the case of stickleback, repeated colonization of freshwater lakes and streams formed tens of thousands of years ago involves standing genetic variation—including multiple large chromosomal inversions—that arose millions of years ago and is maintained polymorphic, perhaps by migration-selection balance, in the ancestral marine population (5). In the case of *An. funestus*, our estimates for inversion and mean candidate mutation ages are older than the inferred K-F split by more than an order of magnitude. While we do not discount a role for new mutations in K, nor do we dismiss the possible contribution of adapted alleles outside of inverted regions, our data emphasize that standing genetic variation in the form of both adapted alleles and recombination modifying inversions has been instrumental for ecotypic divergence of the malaria vector *An. funestus*. Considering its widespread geographic distribution and the high degree of both structural and nucleotide polymorphism segregating in this species, it is plausible that further ecotypes await discovery, and that additional rapid ecotype formation could be predicted in response to new environmental heterogeneities arising from anthropogenic or other changes to the landscape.

### Outlook

Ecotypic differentiation in a heterogeneous environment implies some form of resource partitioning. Despite their genetic divergence, K and F female adult stages share the same specialization on human blood meal hosts (10). Adults of both ecotypes can be found in strict sympatry host seeking and resting inside houses from the same village, although K is less likely than F to rest indoors after a blood meal (10). This behavioral difference is epidemiologically if not ecologically relevant, as an exophilic fraction of the malaria vector population is expected to reduce overall exposure to indoor-based vector control measures. However, the inability to directly karyotype larvae means that evidence for the proposed ecological divergence between the immature stages of K and F is only circumstantial thus far, based on repeated temporal patterns of shifting relative abundance of the adult stages, correlated with seasonal variation in climate and land use (33). In the better-studied Afrotropical *An. gambiae* complex, resource partitioning of the larval habitat has long been appreciated. The most striking ecological difference among *An. gambiae s.l*. taxa is the larval habitat, a characteristic correlated with chromosome structural differences involving the central part of chromosome 2R in this group (64). It was the advent of molecular diagnostics for *An. gambiae s.l*. that empowered field studies aimed at identifying the larval adaptative traits and ecological variables promoting divergent selection in alternative larval habitats (e.g., 65). We anticipate that the development of diagnostics for K and F will similarly advance our understanding of ecotypic differentiation in *An. funestus*, by facilitating direct studies of larval stages, as well as by enabling rapid surveillance of adult females of ecotypes that likely play different roles in malaria transmission in Burkina Faso, if not beyond.

## Materials and Methods

Please see *SI Appendix* Text for detailed materials and methods.

## Supporting information

SampleInfo

## Acknowledgments

We thank Claudia Witzig, and previous members of the Emrich lab (especially Aaron Steele) for their numerous intellectual and analytical contributions to an earlier conception of this project based on population pool sequencing and an unanchored genome assembly. We thank Matt Hahn, Jeff Powell, members of the Besansky lab, and members of the Kern-Ralph Co-lab for discussion, Leo Speidel for help with implementing *Relate*, and developers of *tskit* and *tsdate* for discussion and coding suggestions. We gratefully acknowledge reviewer suggestions that improved the manuscript.

## Author contributions

Conceptualization and design: S.T.S, N.J.B.; Planning, conduct, supervision of field collections and contribution of study materials: C.C., N’F.S., M.W.G.; Analysis and visualization: S.T.S.; Contribution to analysis: M.C.F., S.J.E.; Funding and supervision: N.J.B., A.D.K.; Wrote the paper: S.T.S., N.J.B.; Edited and reviewed the paper: all authors.

## Data and materials availability

Genomic resources used in this study can be downloaded from VectorBase: (i) *An. funestus* reference assembly, www.vectorbase.org/common/downloads/Legacy%20VectorBase%20Files/Anopheles-funestus/Anopheles-funestus-FUMOZ_CHROMOSOMES_AfunF3.fa.gz; (ii) repeat annotations used for masking, www.vectorbase.org/common/downloads/Legacy%20VectorBase%20Files/Anopheles-funestus/Anopheles-funestus-FUMOZ_REPEATFEATURES_AfunF3.gff3.gz; (iii) gene annotations, www.vectorbase.org/common/downloads/Legacy%20VectorBase%20Files/Anopheles-funestus/Anopheles-funestus-FUMOZ_BASEFEATURES_AfunF3.1.gff3.gz. All sequencing data are available as aligned bam-formatted files in NCBI Sequence Read Archives under accessions listed in Table S1. Custom scripts used for analyses are available at GitHub: www.github.com/stsmall/abc_scripts2 and www.github.com/stsmall/Kiribina_Folonzo. The latter link also contains a Jupyter Notebook (www.jupyter.org/) that can be used to recreate figures in the manuscript. Data files are available in the Figshare repository, including: (i) PCA-associated Zarr files for use in scikit-allel (10.6084/m9.figshare.21585543); (ii) admixture proportions (10.6084/m9.figshare.21585555); (iii) nucleotide diversity (10.6084/m9.figshare.21585561); (iv) recombination (10.6084/m9.figshare.21585537); (v) input file for Stairway Plot 2 (10.6084/m9.figshare.21585540); (vi) FILET analyses (10.6084.m9.figshare.21585741); (vii) selection analyses (10.6084/m9.figshare.21585534); (viii) rDNA and GPCR alignments (10.6084/m9.figshare.21585573).

