## Supplementary material for "Standing genetic variation and chromosome differences drove rapid ecotype formation in a major malaria mosquito": SampleInfo

**This PDF file includes:**

Supplementary Text
Figs. S1 to S15
Tables S1 to S13
SI References

SUPPLEMENTARY TEXT

### Mosquito sampling

<u>Burkina Faso</u>

Eleven villages were sampled along a ~200 km east-west transect in Burkina Faso from September to December of 2000-2002 (1, 2) (Fig. S1). Villages consist of family compounds, each containing 2–13 closely spaced huts surrounded by fencing. Adult mosquito indoor resting collections were made from multiple huts and compounds per village by pyrethrum spray catches.

Initial processing of freshly collected mosquitoes was performed in the field. Morphological identification to the Funestus Group (3) was made under a dissecting scope. Ovaries from *An. funestus s.s.* females at the appropriate gonotrophic stage were dissected into individually labeled tubes containing modified Carnoy's solution and held on ice until they could be stored at -20°C for later polytene chromosome analysis. Mosquito carcasses were placed into correspondingly labeled individual tubes with desiccant and stored at room temperature for later DNA analysis.

Following transport to the laboratory, *An. funestus s.s.* was distinguished from potentially co-occurring cryptic species using an intact leg or leg fragment as template in an rDNA-based PCR diagnostic assay (4) as modified by (1). Inversion karyotyping was initially conducted using classical cytogenetic methods (polytene chromosome analysis) (5), and later validated computationally following whole genome sequencing, using PCA (6).

<u>Other African localities</u>

Previously sequenced specimens of *An. funestus s.s.* from three other countries in West, East and southern Africa (7) were added for geographic context, given the pan-African distribution of this species (8). In addition, previously sequenced specimens from other closely related species in the *An. funestus* complex (*An. funestus-like* and *An. longipalpis type C*) were included as outgroups for inferring the ancestral allelic state (7).

Detailed information for the full set of mosquitoes included in this study appears in Table S1.

**Whole genome sequencing and sequence processing**

Genomic DNA extracted from single mosquito carcasses using a CTAB method (9) was quantified via fluorometry (Quantifluor dsDNA System, Promega Corp). Female mosquitoes with >500 ng total genomic DNA were selected for individual library preparation. Libraries were prepared at McGill University and Génome Québec Innovation Centre (Montreal, Canada), and sequenced on the HiSeq X with 150 paired-end cycles.

Adapter sequences and low-quality bases were removed from sequencing reads using trim_galore (github.com/FelixKrueger/TrimGalore). Read pairs with either read shorter than 75 base pairs were removed. Trimmed reads were decontaminated by aligning to a custom file of PhiX and bacterial genomes (*Pantoea sp., Asaia bogorensis*, *Enterobacter asburiae*, *Klebsiella oxytoca*, *K. variicola*, and *Pseudomonas aeruginosa*) using BWA v0.7.17 (10). Only unmapped read pairs were retained. Trimmed and decontaminated reads were then aligned to the nuclear and mitochondrial reference genome of *An. funestus s.s*. (11) using BWA. See Data Availability for *An. funestus* reference version and links.

Variants were called separately for each individual mosquito using GATK v.4.2.1 (12) with HaplotypeCaller using the following options: --emit-ref-confidence GVCF --heterozygosity 0.01 --indel-heterozygosity 0.001 --min-base-quality-score 17. Variant filtering was done in two steps. First, the resulting GVCFs produced by HaplotypeCaller were genotyped using GenotypeGVCFs and each individual VCF file was filtered based on the following metrics: QD < 5, QUAL < 30, DP < 14, MQ < 30, MQRankSum < -12.5, ReadPosRankSum < -8.0, FS > 60.0. Next, filtered GVCFs were merged into a single population GVCF using CombineGVCFs followed again by GenotypeGVCFs. This produced a single VCF with all variant and invariant sites for all sequenced mosquitoes. We refer to this VCF as the *An. funestus* Burkina Faso VCF.

For the final *An. funestus* Burkina Faso VCF, repeats were masked using an available repeat table and regions identified using SNPable (http://lh3lh3.users.sourceforge.net/snpable.shtml) (see Data Availability). After masking, there remained ~160 Mb of accessible sites. Genotypes with a GQ < 30 and DP <20 were marked as missing and individuals with > 20% of callable sites missing were removed. Finally, any sites with > 10% missing genotype calls were removed.

The allelic state was polarized using the program *est-sfs* v2.03 (13) with *An. funestus-like* and *An. longipalpis type C* as outgroup species (7). Read mapping and SNP calling for the outgroup species followed the same methods as detailed above. To avoid having to liftover all sites, we aligned the outgroup species to the *An. funestus* reference genome. The *An. funestus-like* and *An. longipalpis type C* VCFs were merged with the *An. funestus* Burkina Faso VCF using bcftools v1.12 (14) and the command 'merge' with default options. Only sites polymorphic in *An. funestus s.s.* were retained. An input file for *est-sfs* was constructed using the custom script estsfs_format.py to partition the VCF into files with 100,000 sites. The program *est-sfs* was run with default parameters and the results integrated back into the *An. funestus* Burkina Faso VCF by adding an Ancestral Allele (AA) state to the INFO column of the VCF using the custom script polarize_vcf.py. We successfully assigned 99.5% of callable sites to an ancestral state conditioning on an assignment probability of >90% from *est-sfs* results.

Read mapping and SNP calling for the 11 additional *An. funestus s.s.* specimens collected outside of Burkina Faso (Table S1) followed the same methods as detailed above. To make a single pan-African VCF inclusive of all *An. fuenstus s.s*. samples we used bcftools v1.12 to merge the two separate VCFs containing variant and invariant sites.

All sequences have been deposited into GenBank's Short Read Archive, under accession numbers provided in Table S1 (if used in this study) and in Table S2 (if they were not used due to poor data quality). Counts of SNPs by chromosome arm in the total sample of 168 *An. funestus* from Burkina Faso are listed in Table S3. SNP counts by village and ecotype are provided in Table S4.

**Population Structure**

We began by using principal component analysis (PCA) in scikit-allel v1.1.3 (15). We removed any SNP positions with missing data and minor allele frequency <5%. To mitigate dependence among nearby polymorphic sites, we thinned for linkage disequilibrium (LD). Using the $R^2$ statistic as a measure of LD, we excluded sites above a threshold of 0.01. PCA was computed from 100,000 sites randomly selected from those remaining. Variance was scaled according to (16). Following the same methods, PCA was subsequently performed for each autosome arm (Fig 1A; Figs S3-S5).

To quantify genomic ancestry of K and F in terms of admixture proportions, we used the likelihood-free method implemented by the program ALStructure (17). VCF files for each chromosome arm, previously thinned for LD (see above), were converted to bed format using PLINK 1.9 (18). ALStructure was run using a two-cluster admixture model, performing five separate analyses (for the acrocentric X chromosome and each autosome arm) (Fig 1B).

**Estimating recombination maps**

To account for non-uniform recombination rates along the genome, we estimated recombination maps for K and F ecotypes with *ReLERNN* v1.0.0 (19), a machine learning method that uses recurrent neural networks. *ReLERNN* was run on the unphased and filtered *An. funestus* Burkina Faso VCF using a TitanX gpu with set options, --assumedMu 2.8E-9 --upperRhoThetaRatio 35 --unphased. The training of ReLERNN did not include a demographic model of population change for K and F. The classifier was trained using default options and recombination was predicted using default options except for setting the option, --batchSizeOverride to 10000, to avoid memory overflow. The resulting recombination maps were translated to cMMb and HapMap3 format using a custom script rho2cMMb.py (Data Availability).

Median recombination rates were similar among K and F (4.6E$^{-8}$ and 5.E$^{-8}$) and generally agreed with estimates for *An. funestus* from a previous study (7).

**Statistical phasing**

VCF files for each chromosome and ecotype were phased using SHAPEIT v.4.1.3 (20). SHAPEIT was run using the estimated recombination map (see above) and options,

--use-PS 0.0001 --pbwt-mac 1 --pbwt-mdr 0.10. The option --use-PS uses phase set (PS) information (*i.e*., PS fields from GATK HaplotypeCaller) during the phasing.

**Demographic inference**

We reconstructed the population history of *An. funestus* in Burkina Faso using approximate Bayesian computation (ABC) (21). Briefly, in the ABC framework data are generated by computer simulations under alternative demographic scenarios (models) of unknown likelihood, using parameter values drawn from a prior distribution. The resulting simulated data sets are reduced to posterior distributions of parameter values (summary statistics) that are compared to those in the observed data. If a model closely approximates the true demographic history, the general expectation is that the observed and simulated data should have comparable summary statistics. However, due to the loss of information inherent in reducing genotypes to summary statistics, and uncertainties regarding weighting their relative importance, we complemented the ABC approach with an independent methodology, Generative Adversarial Networks (GAN). As implemented in *pg-gan* software (22), this alternative approach uses real data to adaptively learn parameters capable of generating simulated data indistinguishable from real data by machine learning. We have additional confidence if parameters and summary statistics generated by ABC and GAN methods are comparable.

In overview, our demographic analyses proceeded in three steps (explained in more detail in the following sections). First, we used the ABC approach for model selection, to identify the most likely of five demographic scenarios representing alternative population histories in Burkina Faso. Second, in the ABC framework, we used the model with the highest probability to simulate millions of datasets. Simulated datasets were used to infer the marginal distributions of population parameters. Third, we derived population parameter estimates using an independent GAN method and compared the summary statistics generated by ABC and GAN.

<u>Model selection among demographic scenarios</u>

Our analyses used genetic data from three population samples of *An. funestus* (Table S1): K and F from Burkina Faso, and a previously sequenced population from Mozambique in the southeastern part of the species range where *An. funestus* likely emerged and expanded across Africa in the past few thousand years (7).

Before initiating the ABC analyses, we calculated prior distributions of population parameter values with Stairway Plot 2 (23), using the unfolded site-frequency spectrum (uSFS). To calculate the uSFS, we inferred the ancestral state for *An. funestus* SNPs using *est-sfs* (13) and a pair of closely related species (*Anopheles longipalpis* and *Anopheles parensis*) as outgroups (7). To mitigate against the effects of selection, we inferred the uSFS from noncoding genomic regions at least 5 kb away from annotated genes on chromosome arms 2L and 3L (chromosome arms lacking in high frequency inversion polymorphisms in the Burkina Faso samples). Each noncoding region was subdivided into 1 kb loci. Any locus lacking SNPs or containing >25% missing and/or masked data was removed. At each of the remaining loci, any SNP whose ancestral state could not be assigned with >90% probability was pruned. Stairway Plot 2 was then run using the resulting uSFS, with pct_training and nrand values set as recommended (23).

Results were expressed in generations and effective population size was scaled using the average mutation rate of $2.8^{-9}$ (24) (Table S5).

Next, we simulated data under five demographic models representing different degrees and timing of gene flow between K and F following their split (Fig. S5). These consisted of (i) population divergence without gene flow (Isolation Model); (ii) population divergence without gene flow for an initial period, followed by secondary contact (Secondary Contact model); (iii) population divergence with continuous gene flow initially, followed by isolation (Isolation-Initial Migration model); (iv) population divergence with continuous gene flow (Isolation-Migration model); and (v) random mating (Panmixia model).

To compare the five demographic models, we simulated 20,000 sets of genotypic data under each model, using the median values of demographic parameters drawn from prior distributions defined from Stairway Plot 2 (23) (Table S5). Simulations were generated using the custom script abc_sims.py (see Data Availability), which calls the coalescent simulation program *msprime* v1.0.2 (25). We estimated gene flow over four different epochs, whose timing were free parameters in the model. We also allowed for asymmetrical migration rate changes between K and F through time, and variation in effective population sizes of the ancestral (KF) and descendant (K and F) lineages.

To summarize the distribution of genetic diversity within and between ecotypes, we calculated 529 summary statistics for each simulation (Table S6). Following (26), we also modeled sequencing error via an additional parameter that was estimated together with demographic parameters, to improve demographic inference when retaining singleton sites. These statistics were computed in 50,000 bp non-overlapping windows, using the custom python script abc_stats.py (see Data Availability).

Corresponding summary statistics for observed data were calculated from the VCF file containing *An. funestus* from Burkina Faso (K and F ecotypes) and Mozambique, using the custom script abc_stats.py with the option '--observed'. To mitigate the influence of linked selection and prevent bias when comparing real to simulated data, summary statistics from observed data were calculated from chromosome arms 2L and 3L in windows of 50 kb, located at least 5 kb from defined protein coding regions on autosome arms.

Model choice was based on comparing simulation-derived summary statistics from the five models to summary statistics derived from the observed data, using the random forest approach implemented in the R package *abcrf* v.1.8.1 (27). We first built a classification random forest model of 1,000 decision trees and a training dataset of simulation-based summary statistics from all models (the 'reference table'). We then used the summary statistics computed from the observed data to predict the most likely demographic model, using a regression random forest model with 1000 trees. The random forest computation applied to the observed dataset provides a classification vote for each model (*i.e.*, the number of times a model is selected from the forest of decision trees). The model with the highest classification vote corresponds to the model best suited to the observed data. We assessed the global performance of the models by constructing a confusion matrix detailing the rate of model classification, and calculated the approximate posterior probabilities of competing models (Table S7). We selected the scenario with the highest classification vote and posterior probability as the most likely scenario.

Demographic parameter posterior estimation under the most likely demographic model

The most likely model (Isolation-Initial Migration, IIM) was used as a basis for generating 35,000,000 additional simulated genetic data sets to estimate demographic model parameters within ABC. For each simulation the population parameter values were drawn from prior distributions (Table S5). We also varied the recombination and mutation rates for each simulation by randomly selecting a recombination rate from the *ReLERRN* results and randomly drawing a mutation rate from the range $1.0^{-9}$ – $6.1^{-9}$ (24).

The simulated data were used to compute posterior estimates of parameter values using the R package *abc* (28), with the neural net option and settings tol = 0.01, sizenet = 10, and numnet = 15. Cross-validation was used to examine the effect of tolerance choice on parameter estimates by running 100 cross-validation simulations, using the rejection method and a vector of tolerances, tols = 0.05, 0.01, 0.005. The fit of the priors to the posteriors were examined using the plot function in *abc* for fit and bias. The distribution of summary statistics computed from the simulated data were compared to those obtained from the observed data.

Estimation of demographic model parameters using an independent approach

Our third step in demographic inference was an independent estimation of demographic parameters derived from a GAN approach, *pg-gan* (22), that differs from ABC. It uses real data to implement an adaptive parameter learning approach with the goal of generating simulated data that cannot be distinguished from real data by machine learning. When this goal is reached, the values of the population parameters that were used to generate the simulated data set can be considered as plausible estimates of the parameters of the real data.

A forked version of *pg-gan* (April 5, 2021) was used to simulate genetic data under the accepted (IIM) model (Data Availability; www.github.com/stsmall/pg-gan). We excluded the Mozambique population to reduce the number of parameters to be estimated, and modeled population sizes as exponential functions (growing or shrinking population sizes) to reduce the number of free parameters in the model after initial trials demonstrated poor discriminator accuracy.

The program *pg-gan* was run 30 independent times starting from different random seeds. Runs were evaluated based on final discriminator accuracy where a discriminator accuracy of ~50% demonstrated that the real and simulated data were indistinguishable. Parameter estimates for runs with ~50% discriminator accuracy over the last five steps were retained and parameters values were averaged to produce a single point estimate.

We then performed a posterior model checking analysis under the IIM model, to determine whether the model and population parameter estimates matched the observed genetic data well. We simulated 10,000 new datasets with population parameter values drawn from the posterior distributions of the population parameters estimated from the retained *pg-gan* runs. The median of each observed summary statistic was compared with the distribution of the 10,000 simulated test statistics. We calculated the percentile for each median value within the distribution of the simulated data statistic. All observed medians fell within the 17% – 84% percentiles for K and the 20% – 59% percentiles for F. There were no median values that fell into either tails of the simulated statistics.

Finally, the parameter values inferred from ABC and GAN were compared (Table S8). All the pg-gan estimates were within the ABC confidence intervals for the same parameters. Thus, we considered the pg-gan estimates a point-value of the parameter and retained the ABC confidence intervals to represent uncertainty.

**Estimates of genomic diversity and differentiation**

Diversity and differentiation statistics ($F_{ST}$, $d_{XY}$, and π) were calculated in 10 kb non-overlapping windows using the program *pixy* v1.1.0.beta1 (29), which generates unbiased estimates of nucleotide diversity (π) and divergence ($d_{XY}$) regardless of the amount of missing data. The $F_{ST}$ statistic was calculated using Hudson's estimator of $F_{ST}$ (30) (option --hudson) and the default frequency cut-off of MAF > 5%. Additionally, Tajima's *D* values were calculated separately using the custom script abc_stats.py in 10 kb non-overlapping windows for each population.

**Identifying $F_{ST}$ outliers**

To identify outliers, per-window $F_{ST}$ values were standardized to a Z score by subtracting the median and dividing by the standard deviation, following (31). This was done separately for the co-linear and inverted regions of the genome, in light of the generally elevated values of $F_{ST}$ in inverted regions of the genome owing to reduced recombination between inverted and standard orientations of chromosomal rearrangements. Genome windows with a Z-score ≥ 5 (~p-value < 0.0001) in collinear or inverted genomic partitions were considered $F_{ST}$ outliers and retained for downstream analyses.

**Reconstructing genome-wide genealogies to identify loci under positive selection**

We used *Relate* v1.1.7 (32), following the protocol at https://myersgroup.github.io/relate/, to reconstruct genealogies along the genome for the *An. funestus* from Burkina Faso. Input files were prepared from the phased Burkina Faso VCF with an additional file of masked site coordinates. Input files were then filtered to retain bi-allelic sites and verify the polarity of alleles. *Relate* was run on each chromosome arm separately in --mode All, with options: -m 2.8E-9 -N 400000 --haps $hap --sample $sample --map $map. The branch lengths of Relate-inferred genealogies were then adjusted using the results of the ABC demographic inference as a custom file input. This was done, as recommended, to account for population size variation on coalescent times. Relate-inferred genealogies were also exported to tree sequence format using --mode ConvertToTreeSequence, the format used by *tskit* v0.3.7 (33).

**Detecting selection within $F_{ST}$ outlier windows**

To detect selection within $F_{ST}$ outlier windows, we used the genome-wide genealogies inferred with *Relate* (see above) and the add-on module *DetectSelection.sh* included with the *Relate* package. This is a non-parametric test for selection on a single SNP based on coalescence rates of derived- and ancestral-allele-carrying lineages, and a measure of whether the derived allele's spread can be explained by the standard coalescent model (represented as a $\log_{10}$ p-value). First, we subset each tree so they contained only K or F individuals. Next, *DetectSelection.sh* module was run on each mutation with > 3 copies in the population using options: --mu $2.8^{-9}$ --first_bp $start_IoD

--last_bpf $end_IoD --years_per_gen 1 --coal $pop.coal (see Data Availability). This returns a vector of $\log_{10}$ p-values with a single p-value estimated for each discrete time epoch. Time epochs were inferred by Relate in generations before present and are as follows: 1000000, 719686, 517948, 372759, 268270, 193070, 138950, 100000, 71969, 51795, 37276, 26827, 19307, 13895, 10000, 7197, 5179, 3728, 2683, 1931, 1389, 1000, 0. The R package *qvalue* (https://github.com/StoreyLab/qvalue) was then used to adjust the p-value distribution to account for multiple testing by setting the False Discovery Rate to 0.014%. This resulted in a $\log_{10}$ p-value cutoff of -4, where values ≤ -4 would be interpreted as the SNP being under selection during that time. We retained SNPs that had a least one significant epoch of selection (see Data Availability).

**Estimating inversion and mutation ages**

Inversion age was estimated in two ways. First, the time to the most recent common ancestor (MRCA) of the inversion was calculated from genealogies spanning the inversion using *tskit* (33). For each focal inversion (2Ra, 3Ra, and 3Rb), the MRCA of all mosquitoes homozygous for the inversion was calculated using the custom script tree_stats.py. Second, *GEVA* (34) was used to calculate the age of mutations that are private to individuals homozygous for the inversion. For GEVA, the VCF and map files for each chromosome were converted using *GEVA* --vcf $VCF --map $map. Next, queried positions were divided into separate files and *GEVA* was run in parallel with options: --positions $FILE --Ne 200000 –mut 2.8E-9. Output files were filtered to retain only the summary rows with the filtered flag = 1, indicating the algorithms used heuristic filtering of pairs for quality control, and the joint clock, the average of the mutation and recombination clocks rate estimate.

*GEVA* also was used to estimate the age for every candidate SNP (see Data Availability). *GEVA* was run following the same methods as described for estimating the age of variants in inversions.

**Classifying 'no-migration' between K and F in windows along the genome**

*FILET* (Finding Introgressed Loci using Extra Trees Classifiers) (35) uses simulated training data in a random forest classifier to assign windows along the genome as introgressed or non-introgressed between two populations or species. We used *FILET* to detect regions of the genome that the classifier assigns as "no migration," suggesting that they have remained isolated since divergence between K and F. The collinear and rearranged regions were estimated separately. For collinear regions we utilized all K and F samples. For the rearranged regions, we used only F mosquitoes that were homozygous standard for the three focal inversions (2Ra, 3Ra, 3Rb), referred to here as $F_S$ karyotypes, to compare with K, under the assumption that only the $F_S$ subset of the F sample can recombine with the corresponding chromosomal regions of K, while any F mosquitoes carrying an inverted arrangement are effectively isolated in the rearranged genomic regions due to recombination suppression between opposite orientations of inversions in heterozygotes (36).

The *FILET* classifier was trained on population genetic summary statistics generated using the custom script abc_sims.py, a wrapper for the coalescent simulator *msmove* (github.com/geneva/msmove). The program *msmove* was used to generate 180,000 simulations (60,000 for each migration direction and 'no migration') with parameter combinations drawn from uniform distributions defined by the best fit demographic model (IIM). We masked our simulated data to match masking in our genome data following suggestions at www.github.com/kern-lab/FILET, using the custom python script makeFILETmask.py. We withheld 10,000 simulations for construction of a confusion matrix and to verify posterior cutoffs for reducing false positives.

Population genetic summary statistics for classifying windows were calculated in 10 kb windows with a sliding step of 2 kb, omitting any window for which > 50% of sites were masked or missing. The classifiers were trained using a feature vector of all population genetic summary statistics available to *FILET*. Following classification, we clustered adjacent windows showing evidence of no migration by joining consecutive windows with > 90% probability of classification.

**Functional annotation of SNPs**

The program *SnpEff (37)* was used to annotate selected SNPs from $F_{ST}$ outlier regions with respect to gene structure and potential effect. *SnpEff* was run separately for K and F ecotypes, using default options on each chromosome arm. SNP impact was determined using a custom database built from the *An. funestus* genome version 56 on vectorbase.org (accessed 2022-02-10).

**Assembly of ribosomal DNA (rDNA)**

Ribosomal DNA (rDNA) is a useful tool for molecular taxonomy of very closely related and morphologically indistinguishable sibling species of *Anopheles* (38). Despite the creation of multiple reference assemblies of the *An. funestus* nuclear genome (11, 39, 40), automated assembly of the complete *An. funestus* rDNA locus (or any anopheline rDNA locus) from short read sequence data is prohibited by its highly repetitive nature (> 500 tandemly arrayed genes). Furthermore, assembly of a complete rDNA gene unit is complicated by the highly repetitive nature of the component sequences that are most taxonomically informative for close relatives, *e.g.*, the intergenic spacer and external transcribed spacer (41). Motivated by the goal of identifying sequence variation diagnostic for *An. funestus* ecotypes, we began by searching for a complete rDNA unit contained on a long single molecule (Pacific Biosciences; PacBio) read in the PacBio database generated for construction of the *An. funestus* FUMOZ genome assembly [see (11) for details].

We used BLAST+ 2.7.1 command line tool and publicly available *An. funestus, An. gambiae,* and *An. albimanus* rDNA sequence fragments (41-43) as baits to identify *An. funestus* PacBio reads. PacBio reads containing rDNA were aligned using MAFFT v.7.394 (44) and a consensus was constructed using a majority rule decision for each base pair. The resulting consensus was aligned to the partial rDNA of *An. funestus* (41) to identify the genes and build annotations. The IGS region was defined as the region between the 3'-end of the 28S of one rDNA gene and the 5'-end of the 18S of another rDNA gene. Through this process, we successfully reconstructed the complete 18S, ITS1, 5.8S, ITS2, and 28S sequences, and identified partial IGS upstream of the 18S (4.7 kb) and downstream of the 28S (1.1 kb). This rDNA sequence for *An. funestus* has been submitted to GenBank under accession XXXXXXX. The lengths of each rDNA subunit in the submitted sequence are provided in Table S12.

All sequencing reads from the K and F *An. funestus* ecotypes from Burkina Faso were then aligned to the rDNA consensus and SNPs were called following the methods described above. The resulting VCF was used, along with the rDNA assembly, to create a FASTA-formatted sequence for each individual mosquito. These FASTA sequences were then aligned in Geneious Prime 2022.1.1 (https://www.geneious.com) using MAFFT to explore possible diagnostic sequence differences. A neighbor-joining tree was constructed using the Geneious tree builder under the HKY model (Fig. S14).

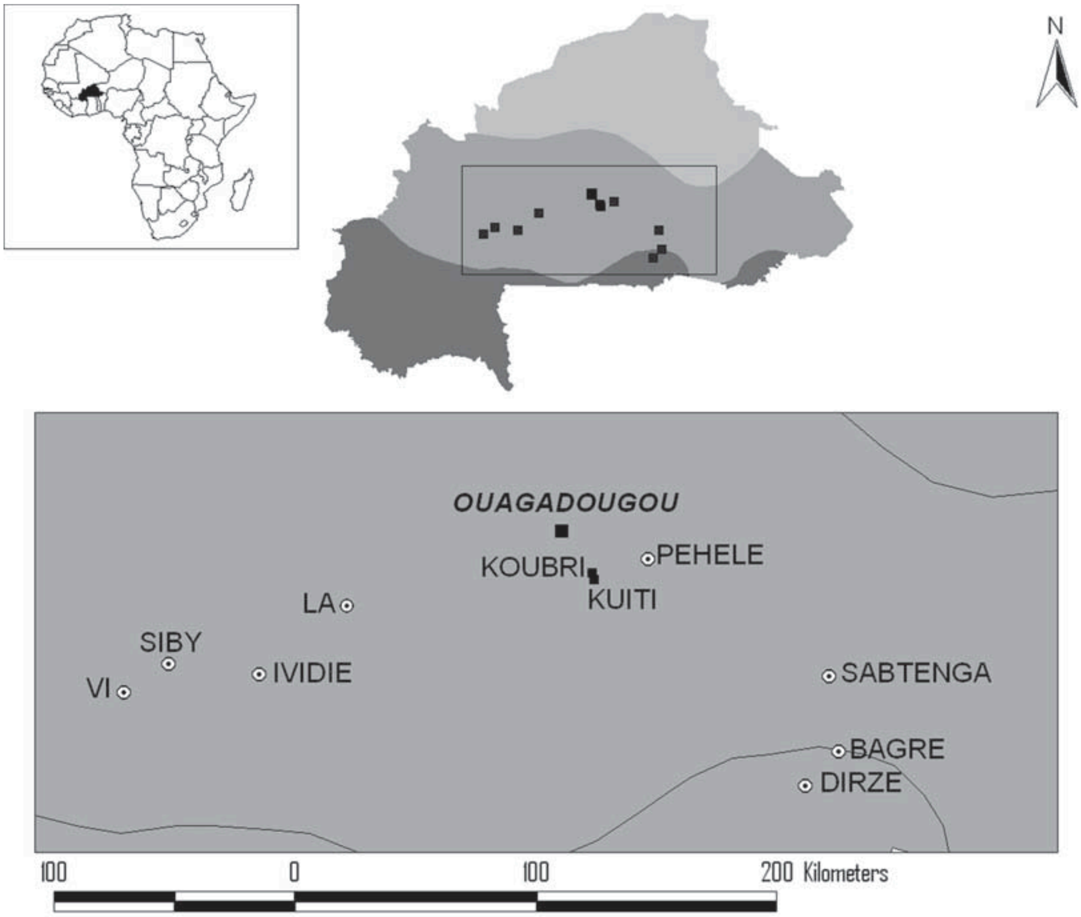

**Figure S1. Map of Burkina Faso showing village sampling locations [modified from (2)].** Detailed sample information can be found in Table S1. For GPS coordinates and further details about the study area, please see (1, 2, 45). Different shading represents three climatic regions (from north to south: Sahel, Sudan–Savanna, and Sudan–Guinea).

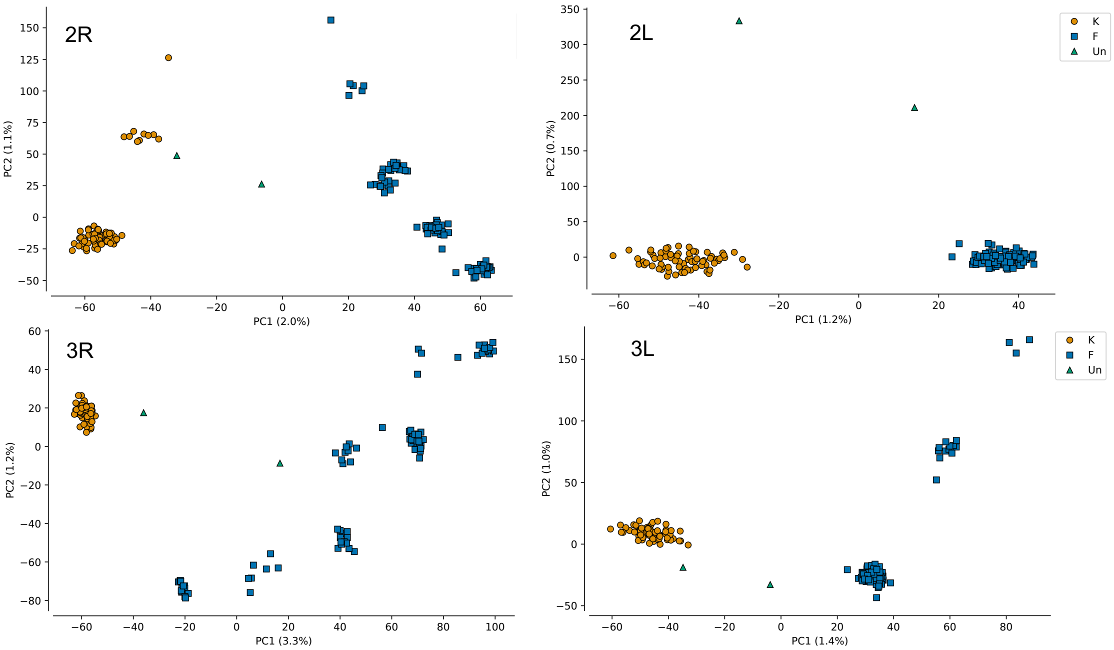

**Figure S2. Genetic structure of *An. funestus* mosquitoes sampled from Burkina Faso**. Plots of the first two principal components from each autosome arm PCA (the corresponding PCA for the X chromosome is shown in Fig 1A). Orange circles and blue squares represent Burkina Faso genotypes assigned to ecotypes K and F based on the X chromosome PCA. Green triangles represent two Burkina Faso genotypes that were unassigned ('Un') based on the X chromosome PCA

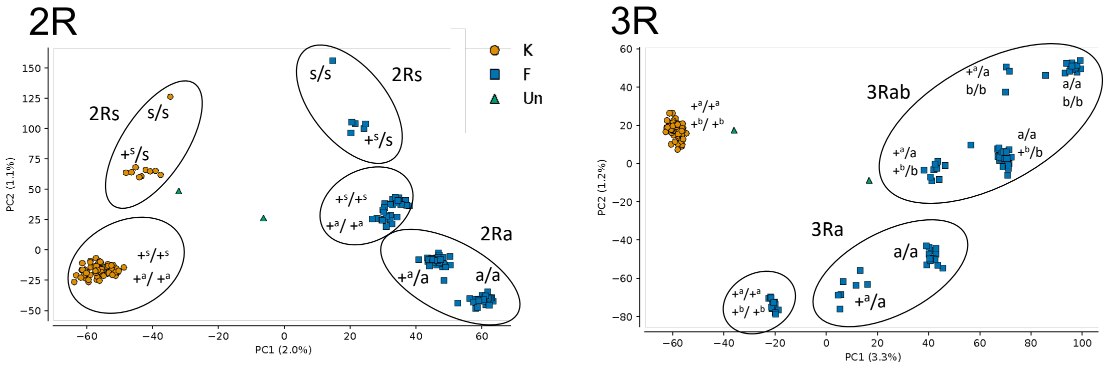

**Figure S3. PCA plot of chromosome arms 2R and 3R, identical to Fig S2, labelled by the computationally determined mosquito inversion genotypes.** Ecotype assignments of individual mosquitoes are represented by orange circles (K), blue squares (F), or green triangles ('Un', unassigned). Inverted orientations of 2Rs, 2Ra, 3Ra, and 3Rb are designated with a lowercase letter (e.g., 's'); standard orientations are represented by '+' with the corresponding letter for the rearrangement given in superscript (e.g., $+^s$). As illustrated by 2Rs, homozygous inverted, standard, and heterozygous genotypes are indicated as s/s, $+^s/+^s$, and $+^s$/s, respectively. Circles are drawn to highlight the specific inversion combinations found in the population. Note that different ecotypes with identical inversion genotypes (e.g., standard orientation of 2Ra, 2Rs, 3Ra, 3Rb) cluster in the PCA by ecotype and not by shared inversion genotype.

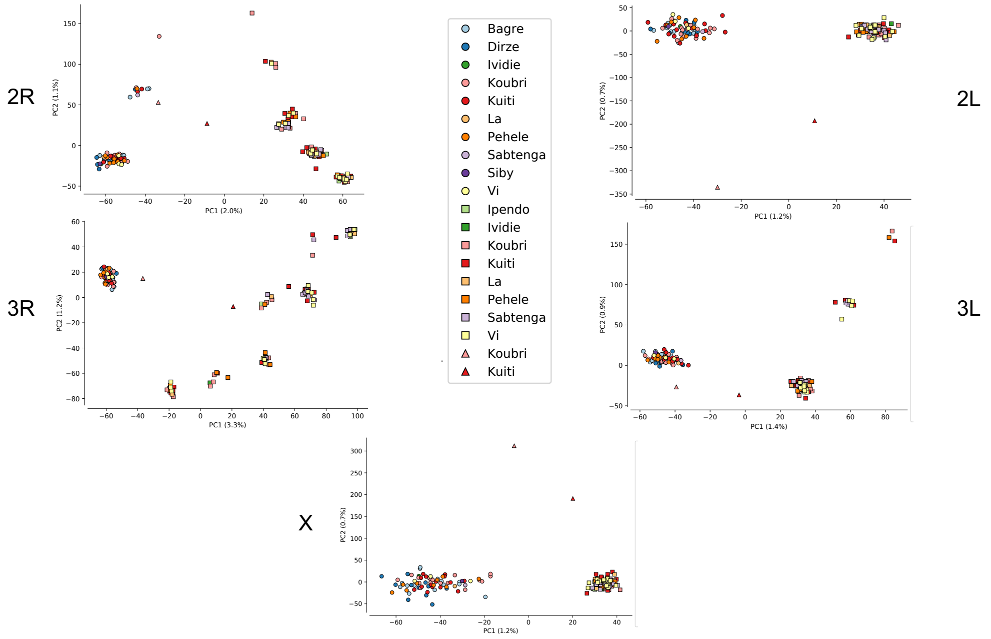

**Figure S4. PCA plots for the X and autosome arms in which mosquito genotypes are colored by the village where they were collected.** Mosquito ecotype assignments, derived from the X chromosome, are represented as different shapes, where circles refer to ecotype K, squares to ecotype F, and triangles to the two unassigned mosquitoes. Colors are used to indicate different villages (see key).

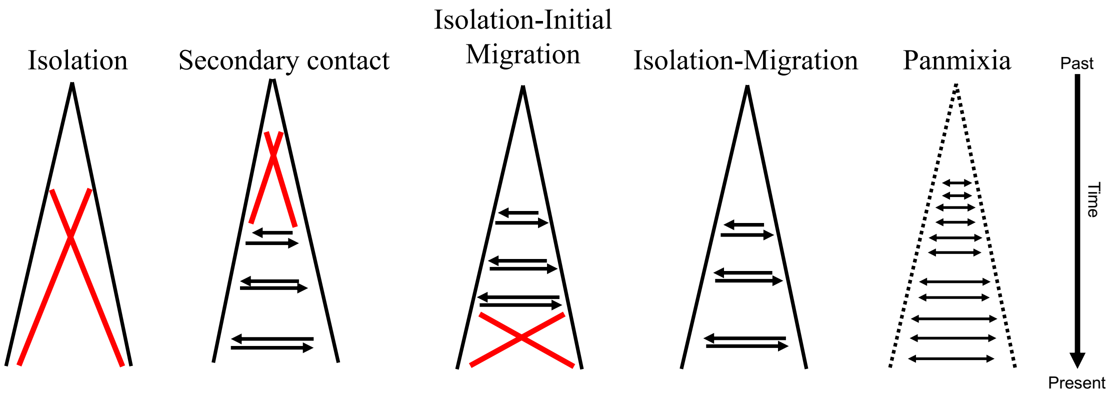

**Figure S5. Five alternative demographic models tested in the ABC framework.** We tested among five different models representing different degrees of isolation between K and F. The Isolation model at left represents complete isolation after divergence (no gene flow). The Panmixia model at right represents random mating. The intervening models were built around temporal isolation and migration. The Secondary contact model represents two populations that were initially isolated and are now connected by gene flow. The Isolation-Initial Migration model resents two populations, between which there is an initial period of gene flow and a subsequent period of isolation. The Isolation-Migration model represents two diverging populations with ongoing gene flow.

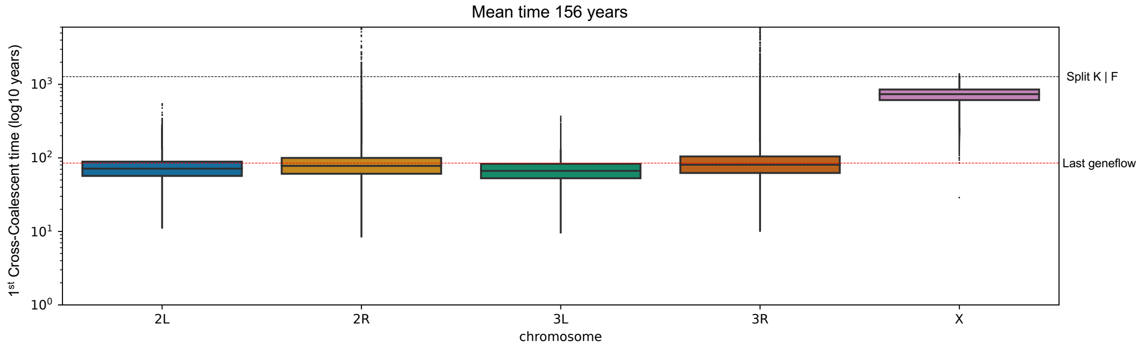

**Figure S6. Mean time to first K-F cross-coalescence by chromosome arm, as inferred from *Relate*** (32). Horizontal dashed lines represent the timing of the K-F split (black) and the timing of last gene flow (red).

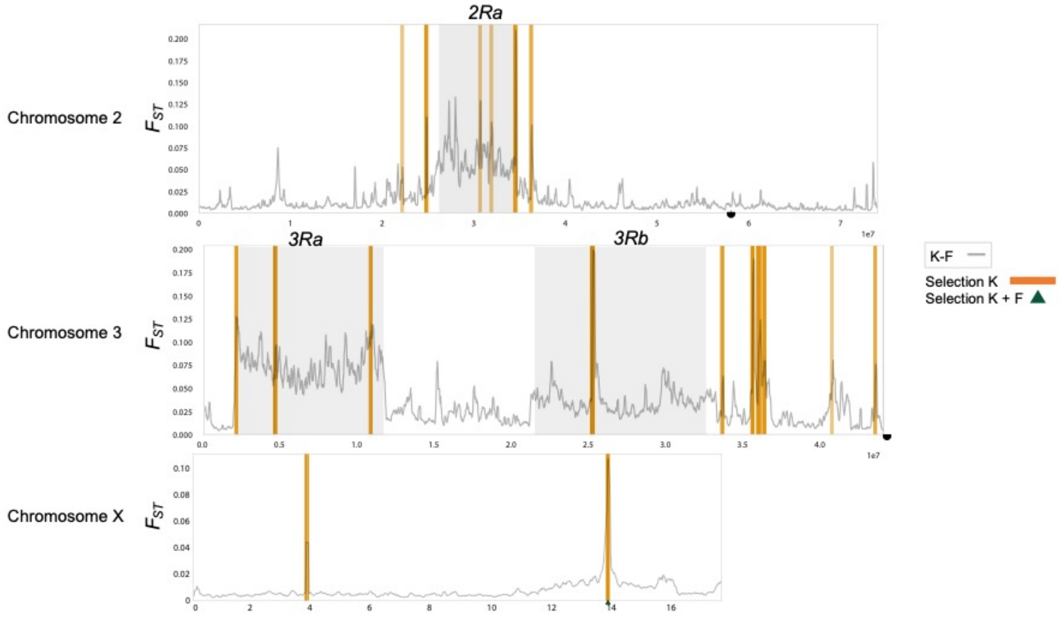

**Figure S7. Genomic locations in the K ecotype where FST outlier windows intersect loci under selection.** Plotted for each chromosome is relative divergence ($F_{ST}$) between K and F, calculated in 10-kb non-overlapping windows and smoothed with a moving average over 10 windows. Orange bars represent loci inferred to be under selection. Triangle indicates positions under selection in both K and F ecotypes. The span of inversions 2Ra, 3Ra, and 3Rb are indicated with gray shading. Centromere location is indicated by half-circles. Note that chromosomes have been truncated to emphasize the relevant regions of overlap between outlier windows and loci under selection.

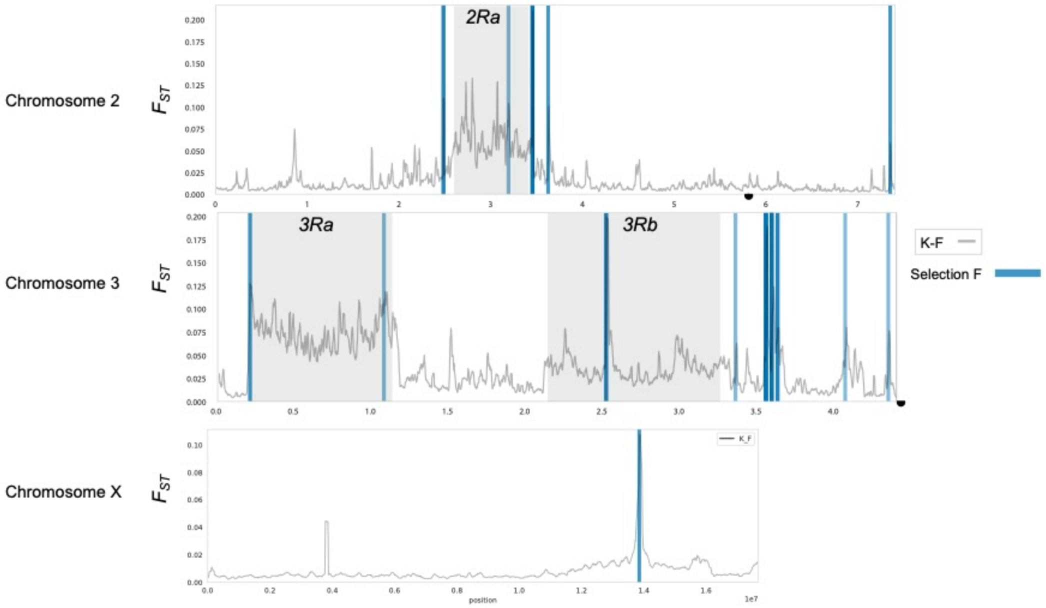

**Figure S8. Genomic locations in the F ecotype where $F_{ST}$ outlier windows intersect loci under selection.** Plotted for each chromosome is relative divergence ($F_{ST}$) between K and F, calculated in 10-kb non-overlapping windows and smoothed with a moving average over 10 windows. Blue bars represent loci inferred to be under selection. The span of inversions 2Ra, 3Ra, and 3Rb are indicated with gray shading. Centromere location is indicated by half-circles. Note that chromosomes have been truncated to emphasize the relevant regions of overlap between outlier windows and loci under selection.

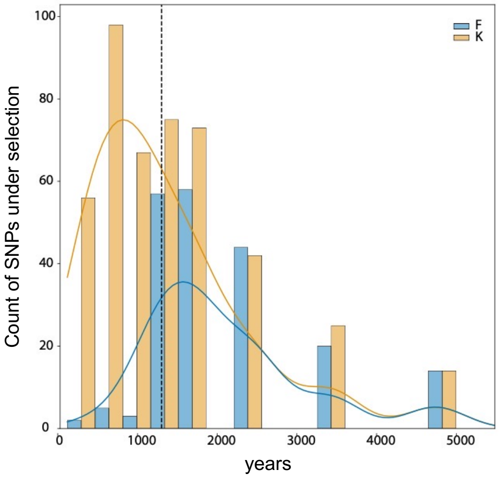

**Figure S9. Histogram of the time of first selection for SNPs under selection in K and F**. Bars are color-coded: blue, F; orange, K. The curves are kernel density estimations (KDE) for the count of selected SNPs in each time interval. The dashed vertical line represents the split time between K and F (~1,300 years ago).

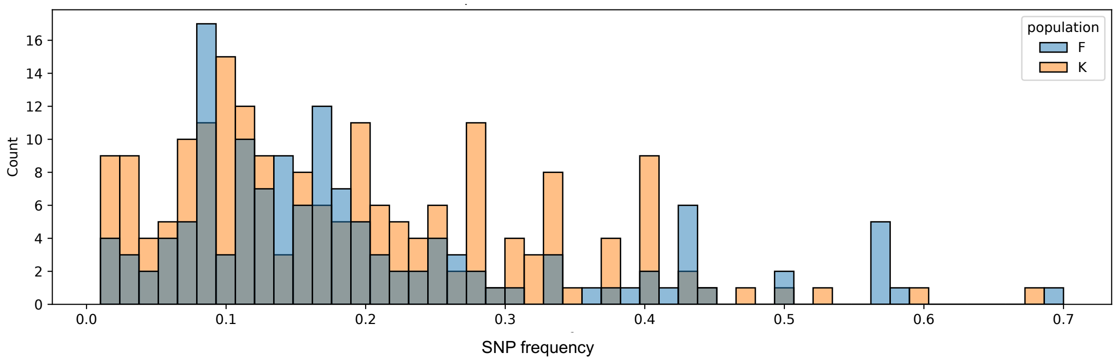

**Figure S10. Frequency of candidate SNPs at the time of first selection.** Bars are blue for F, orange for K, and gray in regions of overlap.

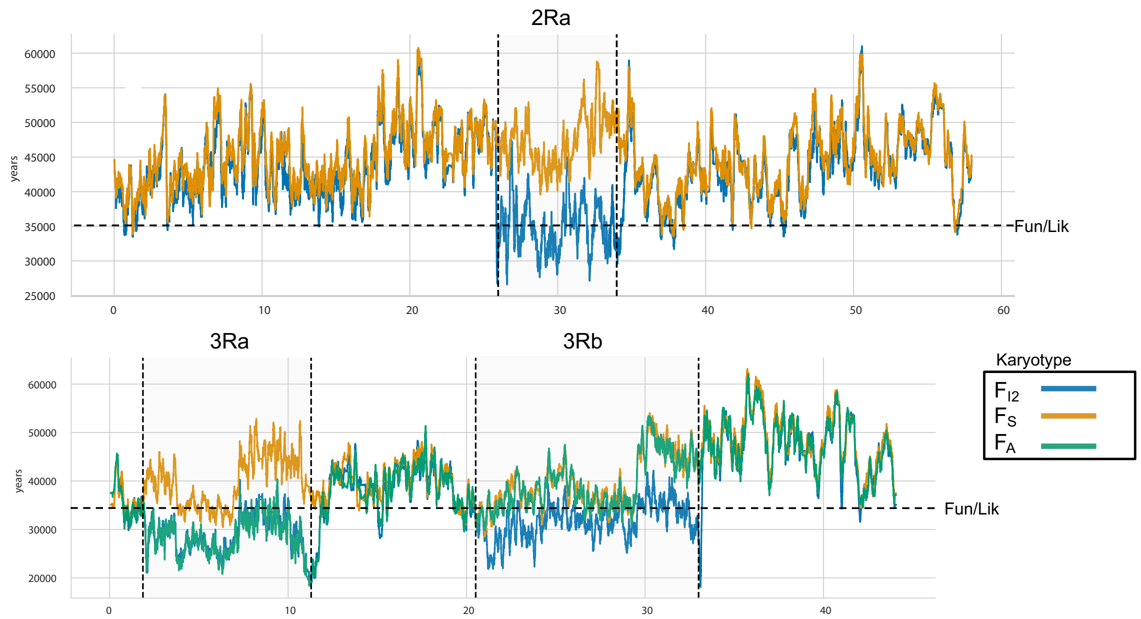

**Figure S11. Estimated ages of inversions 2Ra, 3Ra, and 3Rb**. See SI Appendix text for methodological details. Dashed line represents the inferred split between *An. funestus* (Fun) and its sister species, *An. funestus-like* (Lik). Colored lines represent subsamples of the F ecotype, where $F_S$ refers to mosquitoes that are homozygous standard for the rearrangement, $F_{I2}$ represents F mosquitoes that are homozygous for the inverted orientation, and $F_A$ refers to mosquitoes that are homozygous for the 3Ra inversion but standard for 3Rb.

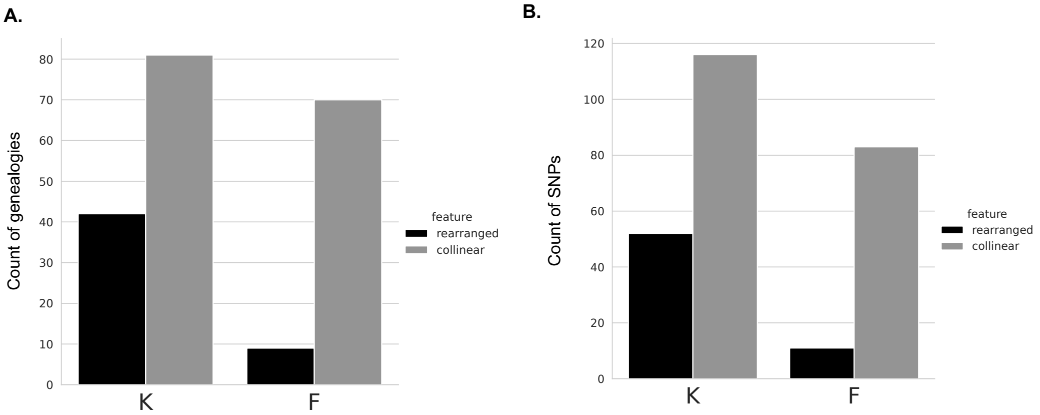

**Figure S12. Numbers of candidate loci and SNPs in inverted and collinear regions in ecotypes K and F**. A) candidate loci (genealogies). B) candidate SNPs.

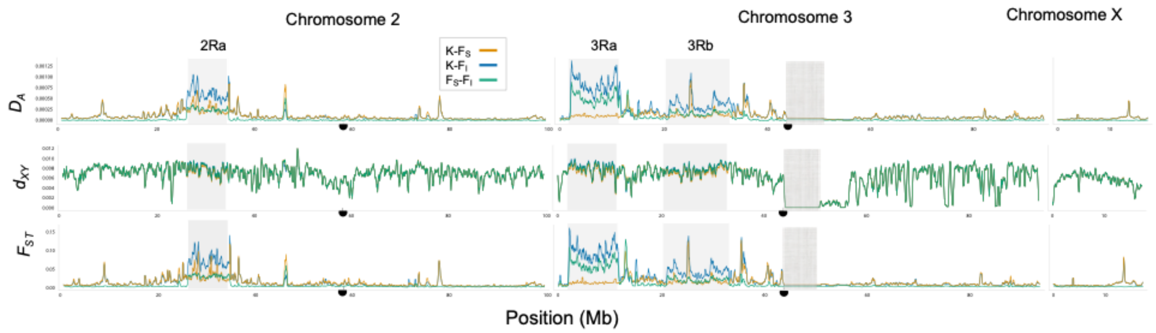

**Figure S13. Relative ($F_{ST}$), absolute ($d_{XY}$) and net divergence ($D_A$) between ecotypes**. F is subdivided by karyotype. $F_S$ refers to mosquitoes that are homozygous for the standard arrangement of inversion 2Ra on chromosome 2R, or 3Ra and 3Rb on chromosome 3R. The remainder, the set of F mosquitoes classified as $F_I$, carry at least one inverted chromosome 2R or 3R, respectively. Inversions are represented as shaded boxes with labels. Centromeres are represented as half circles. Heterochromatic regions, where there was no data, are shown as a textured box.

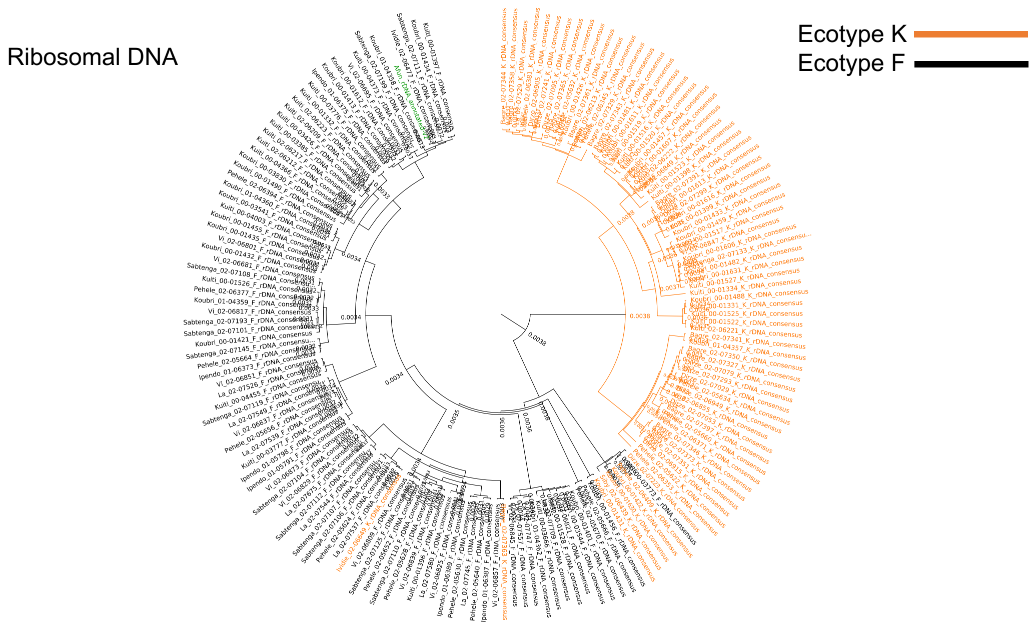

**Figure S14. Neighbor-joining tree of consensus rDNA sequences from individual K and F ecotypes.** Ecotypes K and F are distinguished by orange and black branches and text, respectively. Green text indicates the consensus rDNA assembly constructed from long single molecule PacBio reads from *An. funestus* FUMOZ.

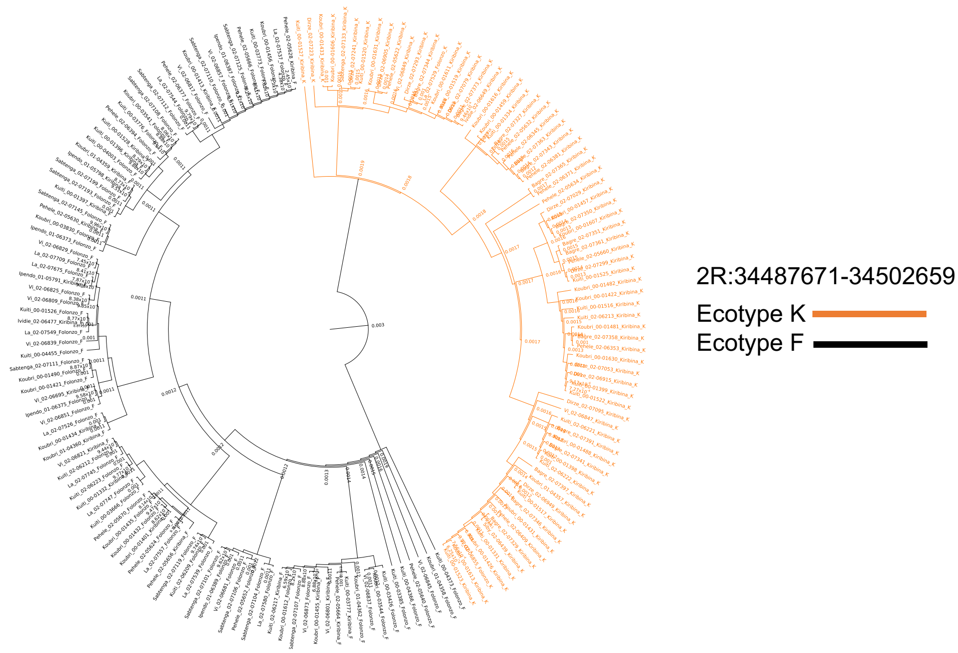

**Figure S15. Neighbor-joining tree based on sequences from individual K and F ecotypes corresponding to the AFUN019981 GPCR gene, spanning positions 34,487,671-34,502,659 on chromosome 2R.**

**Table S1. Information for mosquito samples used in this study.**

| | | | | | | Inversion Genotype[3] | | | | | | |
|---|---|---|---|---|---|---|---|---|---|---|---|---|
| sampleID[1] | **%Cov>10x[2]** | **Biosample ID** | **Country** | **Village** | **Date (M/D/Y)** | **2Ra** | **2Rs** | **3Ra** | **3Rb** | **3La** | **Ecotype[4]** | **Usage in Study** |
| **Ipendo_01-06373_Folonzo** | 0.868 | SAMN27766034 | Burkina Faso | Ipendo | 12/15/01 | 2 | 0 | 2 | 2 | 0 | F | main analyses |
| **Ipendo_01-06375_Folonzo** | 0.880 | SAMN27766035 | Burkina Faso | Ipendo | 12/15/01 | 1 | 0 | 2 | 2 | 0 | F | main analyses |
| **Ipendo_01-06387_Folonzo** | 0.884 | SAMN27766036 | Burkina Faso | Ipendo | 12/15/01 | 1 | 0 | 1 | 1 | 0 | F | main analyses |
| **Ipendo_01-06389_Folonzo** | 0.873 | SAMN27766037 | Burkina Faso | Ipendo | 12/15/01 | 2 | 0 | 2 | 1 | 0 | F | main analyses |
| **Koubri_00-01421_Folonzo** | 0.900 | SAMN27766038 | Burkina Faso | Koubri | 09/04/00 | 1 | 0 | 2 | 0 | 0 | F | main analyses |
| **Koubri_00-01432_Folonzo** | 0.907 | SAMN27766039 | Burkina Faso | Koubri | 09/05/00 | 2 | 0 | 2 | 0 | 0 | F | main analyses |
| **Koubri_00-01435_Folonzo** | 0.904 | SAMN27766040 | Burkina Faso | Koubri | 09/05/00 | 1 | 0 | 2 | 0 | 0 | F | main analyses |
| **Koubri_00-01456_Folonzo** | 0.911 | SAMN27766041 | Burkina Faso | Koubri | 09/06/00 | 0 | 0 | 1 | 1 | 0 | F | main analyses |
| **Koubri_00-01490_Folonzo** | 0.914 | SAMN27766042 | Burkina Faso | Koubri | 09/06/00 | 0 | 0 | 1 | 0 | N | F | main analyses |
| **Koubri_00-01612_Folonzo** | 0.905 | SAMN27766043 | Burkina Faso | Koubri | 09/13/00 | 0 | 1 | 1 | 1 | 0 | F | main analyses |
| **Koubri_00-03541_Folonzo** | 0.908 | SAMN27766044 | Burkina Faso | Koubri | 11/22/00 | 1 | 0 | 1 | 1 | 0 | F | main analyses |
| **Koubri_00-03544_Folonzo** | 0.911 | SAMN27766045 | Burkina Faso | Koubri | 11/22/00 | 2 | 0 | 2 | 0 | 0 | F | main analyses |
| **Koubri_00-03830_Folonzo** | 0.912 | SAMN27766046 | Burkina Faso | Koubri | 11/29/00 | 1 | 0 | 1 | 2 | 0 | F | main analyses |
| **Koubri_01-04358_Folonzo** | 0.905 | SAMN27766047 | Burkina Faso | Koubri | 11/13/01 | 0 | 0 | 1 | 1 | 1 | F | main analyses |
| **Koubri_01-04362_Folonzo** | 0.938 | SAMN27766048 | Burkina Faso | Koubri | 11/13/01 | 1 | 0 | 2 | 1 | 0 | F | main analyses |
| **Kuiti_00-01526_Folonzo** | 0.897 | SAMN27766049 | Burkina Faso | Kuiti | 09/12/00 | 2 | 0 | 2 | 1 | 0 | F | main analyses |
| **Kuiti_00-03385_Folonzo** | 0.904 | SAMN27766050 | Burkina Faso | Kuiti | 11/15/00 | 1 | 0 | 2 | 1 | 0 | F | main analyses |
| **Kuiti_00-03426_Folonzo** | 0.906 | SAMN27766051 | Burkina Faso | Kuiti | 11/20/00 | 1 | 0 | 2 | 1 | 0 | F | main analyses |
| **Kuiti_00-03666_Folonzo** | 0.903 | SAMN27766052 | Burkina Faso | Kuiti | 11/22/00 | 1 | 0 | 1 | 2 | 0 | F | main analyses |
| **Kuiti_00-03773_Folonzo** | 0.904 | SAMN27766053 | Burkina Faso | Kuiti | 11/29/00 | 1 | 0 | 2 | 1 | 0 | F | main analyses |
| **Kuiti_00-03776_Folonzo** | 0.920 | SAMN27766054 | Burkina Faso | Kuiti | 11/29/00 | 2 | 0 | 2 | 1 | 0 | F | main analyses |
| **Kuiti_00-04003_Folonzo** | 0.899 | SAMN27766055 | Burkina Faso | Kuiti | 12/06/00 | 2 | 0 | 2 | 1 | 0 | F | main analyses |

| | | | | | | | | | | | | |
|---|---|---|---|---|---|---|---|---|---|---|---|---|
| **Kuiti_00-04366_Folonzo** | 0.902 | SAMN27766056 | Burkina Faso | Kuiti | 12/13/00 | 0 | 0 | 2 | 2 | 0 | F | main analyses |
| **Kuiti_00-04373_Folonzo** | 0.902 | SAMN27766057 | Burkina Faso | Kuiti | 12/13/00 | 2 | 0 | 2 | 1 | 0 | F | main analyses |
| **Kuiti_00-04455_Folonzo** | 0.868 | SAMN27766058 | Burkina Faso | Kuiti | 12/13/00 | 1 | 0 | 2 | 0 | 0 | F | main analyses |
| **Kuiti_02-06209_Folonzo** | 0.926 | SAMN27766059 | Burkina Faso | Kuiti | 12/02/02 | 1 | 0 | 2 | 0 | 0 | F | main analyses |
| **Kuiti_02-06212_Folonzo** | 0.930 | SAMN27766060 | Burkina Faso | Kuiti | 12/02/02 | 1 | 0 | 2 | 1 | 0 | F | main analyses |
| **Kuiti_02-06223_Folonzo** | 0.939 | SAMN27766061 | Burkina Faso | Kuiti | 12/02/02 | 2 | 0 | 2 | 2 | 1 | F | main analyses |
| **La_02-07526_Folonzo** | 0.869 | SAMN27766062 | Burkina Faso | La | 12/18/02 | 0 | 1 | 0 | 0 | 0 | F | main analyses |
| **La_02-07537_Folonzo** | 0.882 | SAMN27766063 | Burkina Faso | La | 12/18/02 | 2 | 0 | 2 | 2 | 0 | F | main analyses |
| **La_02-07539_Folonzo** | 0.870 | SAMN27766064 | Burkina Faso | La | 12/18/02 | 1 | 0 | 2 | 2 | 1 | F | main analyses |
| **La_02-07544_Folonzo** | 0.886 | SAMN27766065 | Burkina Faso | La | 12/18/02 | 2 | 0 | 2 | 1 | 0 | F | main analyses |
| **La_02-07549_Folonzo** | 0.869 | SAMN27766066 | Burkina Faso | La | 12/18/02 | 2 | 0 | 2 | 2 | N | F | main analyses |
| **La_02-07557_Folonzo** | 0.872 | SAMN27766067 | Burkina Faso | La | 12/18/02 | 1 | 0 | 2 | 1 | N | F | main analyses |
| **La_02-07580_Folonzo** | 0.868 | SAMN27766068 | Burkina Faso | La | 12/18/02 | 1 | 0 | 2 | 1 | 0 | F | main analyses |
| **La_02-07675_Folonzo** | 0.883 | SAMN27766069 | Burkina Faso | La | 12/18/02 | 1 | 0 | 2 | 0 | 0 | F | main analyses |
| **La_02-07709_Folonzo** | 0.877 | SAMN27766070 | Burkina Faso | La | 12/18/02 | 2 | 0 | 2 | 2 | 0 | F | main analyses |
| **La_02-07745_Folonzo** | 0.879 | SAMN27766071 | Burkina Faso | La | 12/18/02 | 1 | 0 | 2 | 1 | N | F | main analyses |
| **La_02-07747_Folonzo** | 0.897 | SAMN27766072 | Burkina Faso | La | 12/18/02 | 1 | 0 | 1 | 1 | 0 | F | main analyses |
| **Pehele_02-05624_Folonzo** | 0.879 | SAMN27766073 | Burkina Faso | Pehele | 11/19/02 | 2 | 0 | 2 | 1 | 0 | F | main analyses |
| **Pehele_02-05640_Folonzo** | 0.867 | SAMN27766074 | Burkina Faso | Pehele | 11/19/02 | 1 | 0 | 2 | 0 | N | F | main analyses |
| **Pehele_02-05652_Folonzo** | 0.882 | SAMN27766075 | Burkina Faso | Pehele | 11/19/02 | 1 | 0 | 2 | 0 | 0 | F | main analyses |
| **Pehele_02-05666_Folonzo** | 0.887 | SAMN27766076 | Burkina Faso | Pehele | 11/19/02 | 0 | 0 | 2 | 1 | 0 | F | main analyses |
| **Pehele_02-05670_Folonzo** | 0.892 | SAMN27766077 | Burkina Faso | Pehele | 11/19/02 | 1 | 0 | 1 | 0 | 0 | F | main analyses |
| **Pehele_02-06377_Folonzo** | 0.867 | SAMN27766078 | Burkina Faso | Pehele | 12/07/02 | 0 | 0 | 2 | 1 | 0 | F | main analyses |
| **Pehele_02-06394_Folonzo** | 0.890 | SAMN27766079 | Burkina Faso | Pehele | 12/07/02 | 2 | 0 | 2 | 0 | 0 | F | main analyses |
| **Sabtenga_02-07101_Folonzo** | 0.874 | SAMN27766080 | Burkina Faso | Sabtenga | 12/15/02 | 1 | 0 | 2 | 1 | 0 | F | main analyses |

| | | | | | | | | | | | | |
|---|---|---|---|---|---|---|---|---|---|---|---|---|
| **Sabtenga_02-07104_Folonzo** | 0.880 | SAMN27766081 | Burkina Faso | Sabtenga | 12/15/02 | 0 | 0 | 2 | 1 | 1 | F | main analyses |
| **Sabtenga_02-07106_Folonzo** | 0.875 | SAMN27766082 | Burkina Faso | Sabtenga | 12/15/02 | 1 | 0 | 1 | 2 | 0 | F | main analyses |
| **Sabtenga_02-07107_Folonzo** | 0.855 | SAMN27766083 | Burkina Faso | Sabtenga | 12/15/02 | 1 | 0 | 1 | 1 | N | F | main analyses |
| **Sabtenga_02-07108_Folonzo** | 0.880 | SAMN27766084 | Burkina Faso | Sabtenga | 12/15/02 | 1 | 0 | 2 | 2 | 0 | F | main analyses |
| **Sabtenga_02-07110_Folonzo** | 0.885 | SAMN27766085 | Burkina Faso | Sabtenga | 12/15/02 | 2 | 0 | 2 | 1 | 0 | F | main analyses |
| **Sabtenga_02-07111_Folonzo** | 0.871 | SAMN27766086 | Burkina Faso | Sabtenga | 12/15/02 | 0 | 0 | 2 | 1 | N | F | main analyses |
| **Sabtenga_02-07112_Folonzo** | 0.867 | SAMN27766087 | Burkina Faso | Sabtenga | 12/15/02 | 1 | 0 | 2 | 1 | 0 | F | main analyses |
| **Sabtenga_02-07119_Folonzo** | 0.853 | SAMN27766088 | Burkina Faso | Sabtenga | 12/15/02 | 1 | 0 | 2 | 0 | 0 | F | main analyses |
| **Sabtenga_02-07125_Folonzo** | 0.863 | SAMN27766089 | Burkina Faso | Sabtenga | 12/15/02 | 2 | 0 | 2 | 2 | 1 | F | main analyses |
| **Sabtenga_02-07145_Folonzo** | 0.893 | SAMN27766090 | Burkina Faso | Sabtenga | 12/15/02 | 0 | 0 | 2 | 2 | 0 | F | main analyses |
| **Sabtenga_02-07193_Folonzo** | 0.883 | SAMN27766091 | Burkina Faso | Sabtenga | 12/15/02 | 2 | 0 | 1 | 1 | 0 | F | main analyses |
| **Sabtenga_02-07199_Folonzo** | 0.910 | SAMN27766092 | Burkina Faso | Sabtenga | 12/15/02 | 0 | 0 | 2 | 1 | N | F | main analyses |
| **Vi_02-06681_Folonzo** | 0.878 | SAMN27766093 | Burkina Faso | Vi | 12/11/02 | 1 | 0 | 2 | 1 | N | F | main analyses |
| **Vi_02-06809_Folonzo** | 0.877 | SAMN27766094 | Burkina Faso | Vi | 12/11/02 | 2 | 0 | 2 | 0 | 0 | F | main analyses |
| **Vi_02-06817_Folonzo** | 0.879 | SAMN27766095 | Burkina Faso | Vi | 12/11/02 | 2 | 0 | 2 | 2 | 0 | F | main analyses |
| **Vi_02-06825_Folonzo** | 0.860 | SAMN27766096 | Burkina Faso | Vi | 12/11/02 | 0 | 0 | 2 | 1 | N | F | main analyses |
| **Vi_02-06829_Folonzo** | 0.876 | SAMN27766097 | Burkina Faso | Vi | 12/11/02 | 2 | 0 | 2 | 2 | 1 | F | main analyses |
| **Vi_02-06837_Folonzo** | 0.894 | SAMN27766098 | Burkina Faso | Vi | 12/11/02 | 0 | 0 | 2 | 0 | 0 | F | main analyses |
| **Vi_02-06839_Folonzo** | 0.873 | SAMN27766099 | Burkina Faso | Vi | 12/11/02 | 2 | 0 | 2 | 1 | 1 | F | main analyses |
| **Vi_02-06845_Folonzo** | 0.885 | SAMN27766100 | Burkina Faso | Vi | 12/11/02 | 2 | 0 | 2 | 2 | N | F | main analyses |
| **Vi_02-06851_Folonzo** | 0.869 | SAMN27766101 | Burkina Faso | Vi | 12/11/02 | 2 | 0 | 2 | 1 | 0 | F | main analyses |
| **Vi_02-06857_Folonzo** | 0.850 | SAMN27766102 | Burkina Faso | Vi | 12/11/02 | 1 | 0 | 2 | 1 | 0 | F | main analyses |
| **Vi_02-06873_Folonzo** | 0.877 | SAMN27766103 | Burkina Faso | Vi | 12/11/02 | 1 | 0 | 2 | 0 | N | F | main analyses |
| **La_02-07529_Folonzo** | 0.880 | SAMN27766104 | Burkina Faso | La | 12/18/02 | 0 | 0 | 0 | 0 | 0 | K | main analyses |

| | | | | | | | | | | | | |
|---|---|---|---|---|---|---|---|---|---|---|---|---|
| **Vi_02-06855_Folonzo** | 0.888 | SAMN27766105 | Burkina Faso | Vi | 12/11/02 | 0 | 0 | 0 | 0 | 0 | K | main analyses |
| **Koubri_00-03530_Folonzo** | 0.926 | SAMN27766116 | Burkina Faso | Koubri | 11/22/00 | 0 | 0 | 2 | 2 | 0 | Un | PCA |
| **Kuiti_02-06207_Folonzo** | 0.939 | SAMN27766117 | Burkina Faso | Kuiti | 12/02/02 | 0 | 0 | 2 | 1 | 0 | Un | PCA |
| **Vi_02-06859_Folonzo** | 0.868 | SAMN27766118 | Burkina Faso | Vi | 12/11/02 | 2 | 0 | 2 | 0 | 0 | Un | PCA |
| **Ipendo_01-05791_Kiribina** | 0.874 | SAMN27766119 | Burkina Faso | Ipendo | 12/15/01 | 2 | 0 | 2 | 2 | 0 | F | main analyses |
| **Ipendo_01-05798_Kiribina** | 0.892 | SAMN27766120 | Burkina Faso | Ipendo | 12/15/01 | 1 | 0 | 2 | 1 | 0 | F | main analyses |
| **Ividie_02-06477_Kiribina** | 0.869 | SAMN27766121 | Burkina Faso | Ividie | 12/10/02 | 0 | 0 | 1 | 0 | 1 | F | main analyses |
| **Koubri_00-01401_Kiribina** | 0.904 | SAMN27766122 | Burkina Faso | Koubri | 09/04/00 | 0 | 0 | 0 | 0 | 0 | F | main analyses |
| **Koubri_00-01413_Kiribina** | 0.900 | SAMN27766123 | Burkina Faso | Koubri | 09/04/00 | 0 | 0 | 0 | 0 | 0 | F | main analyses |
| **Koubri_00-01434_Kiribina** | 0.890 | SAMN27766124 | Burkina Faso | Koubri | 09/05/00 | 0 | 1 | 1 | 0 | 0 | F | main analyses |
| **Koubri_00-01455_Kiribina** | 0.902 | SAMN27766125 | Burkina Faso | Koubri | 09/06/00 | 0 | 2 | 1 | 0 | 0 | F | main analyses |
| **Koubri_01-04359_Kiribina** | 0.910 | SAMN27766126 | Burkina Faso | Koubri | 11/13/01 | 0 | 0 | 0 | 0 | 0 | F | main analyses |
| **Koubri_01-04360_Kiribina** | 0.908 | SAMN27766127 | Burkina Faso | Koubri | 11/13/01 | 0 | 0 | 0 | 0 | 0 | F | main analyses |
| **Kuiti_00-01332_Kiribina** | 0.895 | SAMN27766128 | Burkina Faso | Kuiti | 09/05/00 | 0 | 0 | 0 | 0 | 0 | F | main analyses |
| **Kuiti_00-01396_Kiribina** | 0.899 | SAMN27766129 | Burkina Faso | Kuiti | 09/06/00 | 0 | 0 | 0 | 0 | 0 | F | main analyses |
| **Kuiti_00-01397_Kiribina** | 0.907 | SAMN27766130 | Burkina Faso | Kuiti | 09/06/00 | 0 | 1 | 0 | 0 | 0 | F | main analyses |
| **Kuiti_00-01528_Kiribina** | 0.897 | SAMN27766131 | Burkina Faso | Kuiti | 09/12/00 | 0 | 0 | 0 | 0 | 0 | F | main analyses |
| **Kuiti_00-03777_Kiribina** | 0.904 | SAMN27766132 | Burkina Faso | Kuiti | 11/29/00 | 0 | 0 | 0 | 0 | 0 | F | main analyses |
| **Kuiti_02-06217_Kiribina** | 0.903 | SAMN27766133 | Burkina Faso | Kuiti | 12/02/02 | 0 | 0 | 1 | 0 | 0 | F | main analyses |
| **Pehele_02-05628_Kiribina** | 0.858 | SAMN27766134 | Burkina Faso | Pehele | 11/19/02 | 0 | 0 | 1 | 0 | 0 | F | main analyses |
| **Pehele_02-05630_Kiribina** | 0.865 | SAMN27766135 | Burkina Faso | Pehele | 11/19/02 | 0 | 0 | 0 | 0 | 0 | F | main analyses |
| **Pehele_02-05656_Kiribina** | 0.871 | SAMN27766136 | Burkina Faso | Pehele | 11/19/02 | 0 | 0 | 0 | 0 | 0 | F | main analyses |
| **Pehele_02-05664_Kiribina** | 0.887 | SAMN27766137 | Burkina Faso | Pehele | 11/19/02 | 0 | 0 | 1 | 1 | 0 | F | main analyses |
| **Vi_02-06695_Kiribina** | 0.911 | SAMN27766138 | Burkina Faso | Vi | 12/11/02 | 0 | 1 | 0 | 0 | N | F | main analyses |
| **Vi_02-06801_Kiribina** | 0.898 | SAMN27766139 | Burkina Faso | Vi | 12/11/02 | 0 | 0 | 0 | 0 | 0 | F | main analyses |
| **Vi_02-06821_Kiribina** | 0.936 | SAMN27766140 | Burkina Faso | Vi | 12/11/02 | 0 | 0 | 0 | 0 | N | F | main analyses |

| | | | | | | | | | | | | |
|---|---|---|---|---|---|---|---|---|---|---|---|---|
| **Bagre_02-07327_Kiribina** | 0.868 | SAMN27766141 | Burkina Faso | Bagre | 12/17/02 | 0 | 1 | 0 | 0 | 0 | K | main analyses |
| **Bagre_02-07329_Kiribina** | 0.891 | SAMN27766142 | Burkina Faso | Bagre | 12/17/02 | 0 | 0 | 0 | 0 | 0 | K | main analyses |
| **Bagre_02-07341_Kiribina** | 0.895 | SAMN27766143 | Burkina Faso | Bagre | 12/17/02 | 0 | 0 | 0 | 0 | N | K | main analyses |
| **Bagre_02-07343_Kiribina** | 0.874 | SAMN27766144 | Burkina Faso | Bagre | 12/17/02 | 0 | 0 | 0 | 0 | 0 | K | main analyses |
| **Bagre_02-07344_Kiribina** | 0.882 | SAMN27766145 | Burkina Faso | Bagre | 12/17/02 | 0 | 0 | 0 | 0 | 0 | K | main analyses |
| **Bagre_02-07346_Kiribina** | 0.894 | SAMN27766146 | Burkina Faso | Bagre | 12/17/02 | 0 | 0 | 0 | 0 | 0 | K | main analyses |
| **Bagre_02-07350_Kiribina** | 0.857 | SAMN27766147 | Burkina Faso | Bagre | 12/17/02 | 0 | 0 | 0 | 0 | 0 | K | main analyses |
| **Bagre_02-07351_Kiribina** | 0.881 | SAMN27766148 | Burkina Faso | Bagre | 12/17/02 | 0 | 0 | 0 | 0 | 0 | K | main analyses |
| **Bagre_02-07358_Kiribina** | 0.856 | SAMN27766149 | Burkina Faso | Bagre | 12/17/02 | 0 | 0 | 0 | 0 | 0 | K | main analyses |
| **Bagre_02-07361_Kiribina** | 0.865 | SAMN27766150 | Burkina Faso | Bagre | 12/17/02 | 0 | 1 | 0 | 0 | 0 | K | main analyses |
| **Bagre_02-07363_Kiribina** | 0.862 | SAMN27766151 | Burkina Faso | Bagre | 12/17/02 | 0 | 0 | 0 | 0 | 0 | K | main analyses |
| **Bagre_02-07365_Kiribina** | 0.887 | SAMN27766152 | Burkina Faso | Bagre | 12/17/02 | 0 | 1 | 0 | 0 | 0 | K | main analyses |
| **Bagre_02-07373_Kiribina** | 0.886 | SAMN27766153 | Burkina Faso | Bagre | 12/17/02 | 0 | 0 | 0 | 0 | 0 | K | main analyses |
| **Bagre_02-07391_Kiribina** | 0.892 | SAMN27766154 | Burkina Faso | Bagre | 12/17/02 | 0 | 0 | 0 | 0 | 0 | K | main analyses |
| **Bagre_02-07397_Kiribina** | 0.892 | SAMN27766155 | Burkina Faso | Bagre | 12/17/02 | 0 | 0 | 0 | 0 | 0 | K | main analyses |
| **Dirze_02-06905_Kiribina** | 0.857 | SAMN27766156 | Burkina Faso | Dirze | 12/14/02 | 0 | 0 | 0 | 0 | 0 | K | main analyses |
| **Dirze_02-06915_Kiribina** | 0.872 | SAMN27766157 | Burkina Faso | Dirze | 12/14/02 | 0 | 0 | 0 | 0 | 0 | K | main analyses |
| **Dirze_02-06949_Kiribina** | 0.861 | SAMN27766158 | Burkina Faso | Dirze | 12/14/02 | 0 | 0 | 0 | 0 | 0 | K | main analyses |
| **Dirze_02-07029_Kiribina** | 0.867 | SAMN27766159 | Burkina Faso | Dirze | 12/14/02 | 0 | 0 | 0 | 0 | 0 | K | main analyses |
| **Dirze_02-07053_Kiribina** | 0.871 | SAMN27766160 | Burkina Faso | Dirze | 12/14/02 | 0 | 0 | 0 | 0 | 0 | K | main analyses |
| **Dirze_02-07079_Kiribina** | 0.864 | SAMN27766161 | Burkina Faso | Dirze | 12/14/02 | 0 | 0 | 0 | 0 | 0 | K | main analyses |
| **Dirze_02-07095_Kiribina** | 0.869 | SAMN27766162 | Burkina Faso | Dirze | 12/14/02 | 0 | 0 | 0 | 0 | 0 | K | main analyses |
| **Dirze_02-07223_Kiribina** | 0.856 | SAMN27766163 | Burkina Faso | Dirze | 12/14/02 | 0 | 0 | 0 | 0 | 0 | K | main analyses |
| **Dirze_02-07241_Kiribina** | 0.895 | SAMN27766164 | Burkina Faso | Dirze | 12/14/02 | 0 | 1 | 0 | 0 | 0 | K | main analyses |
| **Dirze_02-07293_Kiribina** | 0.864 | SAMN27766165 | Burkina Faso | Dirze | 12/14/02 | 0 | 0 | 0 | 0 | 0 | K | main analyses |
| **Dirze_02-07299_Kiribina** | 0.884 | SAMN27766166 | Burkina Faso | Dirze | 12/14/02 | 0 | 0 | 0 | 0 | 0 | K | main analyses |

| | | | | | | | | | | | | |
|---|---|---|---|---|---|---|---|---|---|---|---|---|
| **Ividie_02-06649_Kiribina** | 0.890 | SAMN27766167 | Burkina Faso | Ividie | 12/10/02 | 0 | 0 | 0 | 0 | 0 | K | main analyses |
| **Koubri_00-01422_Kiribina** | 0.917 | SAMN27766168 | Burkina Faso | Koubri | 09/04/00 | 0 | 0 | 0 | 0 | 0 | K | main analyses |
| **Koubri_00-01426_Kiribina** | 0.904 | SAMN27766169 | Burkina Faso | Koubri | 09/05/00 | 0 | 0 | 0 | 0 | N | K | main analyses |
| **Koubri_00-01431_Kiribina** | 0.895 | SAMN27766170 | Burkina Faso | Koubri | 09/05/00 | 0 | 0 | 0 | 0 | 0 | K | main analyses |
| **Koubri_00-01433_Kiribina** | 0.903 | SAMN27766171 | Burkina Faso | Koubri | 09/05/00 | 0 | 2 | 0 | 0 | 0 | K | main analyses |
| **Koubri_00-01457_Kiribina** | 0.902 | SAMN27766172 | Burkina Faso | Koubri | 09/06/00 | 0 | 0 | 0 | 0 | 0 | K | main analyses |
| **Koubri_00-01459_Kiribina** | 0.911 | SAMN27766173 | Burkina Faso | Koubri | 09/06/00 | 0 | 0 | 0 | 0 | 0 | K | main analyses |
| **Koubri_00-01481_Kiribina** | 0.905 | SAMN27766174 | Burkina Faso | Koubri | 09/06/00 | 0 | 0 | 0 | 0 | 0 | K | main analyses |
| **Koubri_00-01482_Kiribina** | 0.909 | SAMN27766175 | Burkina Faso | Koubri | 09/06/00 | 0 | 0 | 0 | 0 | 0 | K | main analyses |
| **Koubri_00-01488_Kiribina** | 0.908 | SAMN27766176 | Burkina Faso | Koubri | 09/06/00 | 0 | 0 | 0 | 0 | 0 | K | main analyses |
| **Koubri_00-01606_Kiribina** | 0.909 | SAMN27766177 | Burkina Faso | Koubri | 09/12/00 | 0 | 1 | 0 | 0 | 0 | K | main analyses |
| **Koubri_00-01607_Kiribina** | 0.890 | SAMN27766178 | Burkina Faso | Koubri | 09/12/00 | 0 | 0 | 0 | 0 | 0 | K | main analyses |
| **Koubri_00-01611_Kiribina** | 0.904 | SAMN27766179 | Burkina Faso | Koubri | 09/13/00 | 0 | 0 | 0 | 0 | 0 | K | main analyses |
| **Koubri_00-01613_Kiribina** | 0.898 | SAMN27766180 | Burkina Faso | Koubri | 09/13/00 | 0 | 0 | 0 | 0 | 0 | K | main analyses |
| **Koubri_00-01616_Kiribina** | 0.903 | SAMN27766181 | Burkina Faso | Koubri | 09/13/00 | 0 | 0 | 0 | 0 | 0 | K | main analyses |
| **Koubri_00-01630_Kiribina** | 0.905 | SAMN27766182 | Burkina Faso | Koubri | 09/13/00 | 0 | 0 | 0 | 0 | 0 | K | main analyses |
| **Koubri_00-01631_Kiribina** | 0.922 | SAMN27766183 | Burkina Faso | Koubri | 09/13/00 | 0 | 0 | 0 | 0 | 0 | K | main analyses |
| **Koubri_01-04357_Kiribina** | 0.903 | SAMN27766184 | Burkina Faso | Koubri | 11/13/01 | 0 | 0 | 0 | 0 | 0 | K | main analyses |
| **Kuiti_00-01331_Kiribina** | 0.899 | SAMN27766185 | Burkina Faso | Kuiti | 09/05/00 | 0 | 0 | 0 | 0 | 0 | K | main analyses |
| **Kuiti_00-01334_Kiribina** | 0.905 | SAMN27766186 | Burkina Faso | Kuiti | 09/05/00 | 0 | 0 | 0 | 0 | 0 | K | main analyses |
| **Kuiti_00-01398_Kiribina** | 0.902 | SAMN27766187 | Burkina Faso | Kuiti | 09/06/00 | 0 | 0 | 0 | 0 | 0 | K | main analyses |
| **Kuiti_00-01399_Kiribina** | 0.909 | SAMN27766188 | Burkina Faso | Kuiti | 09/06/00 | 0 | 0 | 0 | 0 | 0 | K | main analyses |
| **Kuiti_00-01516_Kiribina** | 0.907 | SAMN27766189 | Burkina Faso | Kuiti | 09/11/00 | 0 | 0 | 0 | 0 | 0 | K | main analyses |
| **Kuiti_00-01517_Kiribina** | 0.901 | SAMN27766190 | Burkina Faso | Kuiti | 09/11/00 | 0 | 0 | 0 | 0 | 0 | K | main analyses |
| **Kuiti_00-01519_Kiribina** | 0.909 | SAMN27766191 | Burkina Faso | Kuiti | 09/11/00 | 0 | 0 | 0 | 0 | 0 | K | main analyses |
| **Kuiti_00-01520_Kiribina** | 0.906 | SAMN27766192 | Burkina Faso | Kuiti | 09/11/00 | 0 | 1 | 0 | 0 | 0 | K | main analyses |

| | | | | | | | | | | | | |
|---|---|---|---|---|---|---|---|---|---|---|---|---|
| **Kuiti_00-01522_Kiribina** | 0.899 | SAMN27766193 | Burkina Faso | Kuiti | 09/11/00 | 0 | 1 | 0 | 0 | 0 | K | main analyses |
| **Kuiti_00-01523_Kiribina** | 0.911 | SAMN27766194 | Burkina Faso | Kuiti | 09/11/00 | 0 | 0 | 0 | 0 | 0 | K | main analyses |
| **Kuiti_00-01525_Kiribina** | 0.895 | SAMN27766195 | Burkina Faso | Kuiti | 09/12/00 | 0 | 0 | 0 | 0 | 0 | K | main analyses |
| **Kuiti_00-01527_Kiribina** | 0.902 | SAMN27766196 | Burkina Faso | Kuiti | 09/12/00 | 0 | 0 | 0 | 0 | 0 | K | main analyses |
| **Kuiti_02-06213_Kiribina** | 0.903 | SAMN27766197 | Burkina Faso | Kuiti | 12/02/02 | 0 | 0 | 0 | 0 | 0 | K | main analyses |
| **Kuiti_02-06221_Kiribina** | 0.876 | SAMN27766198 | Burkina Faso | Kuiti | 12/02/02 | 0 | 0 | 0 | 0 | 0 | K | main analyses |
| **Kuiti_02-06222_Kiribina** | 0.897 | SAMN27766199 | Burkina Faso | Kuiti | 12/02/02 | 0 | 0 | 0 | 0 | 0 | K | main analyses |
| **Pehele_02-05622_Kiribina** | 0.887 | SAMN27766200 | Burkina Faso | Pehele | 11/19/02 | 0 | 0 | 0 | 0 | 0 | K | main analyses |
| **Pehele_02-05632_Kiribina** | 0.884 | SAMN27766201 | Burkina Faso | Pehele | 11/19/02 | 0 | 0 | 0 | 0 | 0 | K | main analyses |
| **Pehele_02-05634_Kiribina** | 0.859 | SAMN27766202 | Burkina Faso | Pehele | 11/19/02 | 0 | 1 | 0 | 0 | 0 | K | main analyses |
| **Pehele_02-05660_Kiribina** | 0.874 | SAMN27766203 | Burkina Faso | Pehele | 11/19/02 | 0 | 0 | 0 | 0 | 0 | K | main analyses |
| **Pehele_02-06345_Kiribina** | 0.872 | SAMN27766204 | Burkina Faso | Pehele | 12/07/02 | 0 | 0 | 0 | 0 | 0 | K | main analyses |
| **Pehele_02-06353_Kiribina** | 0.858 | SAMN27766205 | Burkina Faso | Pehele | 12/07/02 | 0 | 0 | 0 | 0 | N | K | main analyses |
| **Pehele_02-06371_Kiribina** | 0.845 | SAMN27766206 | Burkina Faso | Pehele | 12/07/02 | 0 | 0 | 0 | 0 | 0 | K | main analyses |
| **Pehele_02-06381_Kiribina** | 0.857 | SAMN27766207 | Burkina Faso | Pehele | 12/07/02 | 0 | 0 | 0 | 0 | 0 | K | main analyses |
| **Pehele_02-06409_Kiribina** | 0.901 | SAMN27766208 | Burkina Faso | Pehele | 12/07/02 | 0 | 0 | 0 | 0 | 0 | K | main analyses |
| **Sabtenga_02-07133_Kiribina** | 0.891 | SAMN27766209 | Burkina Faso | Sabtenga | 12/15/02 | 0 | 1 | 0 | 0 | 0 | K | main analyses |
| **Siby_02-06439_Kiribina** | 0.876 | SAMN27766210 | Burkina Faso | Siby | 12/09/02 | 0 | 0 | 0 | 0 | 0 | K | main analyses |
| **Vi_02-06847_Kiribina** | 0.922 | SAMN27766211 | Burkina Faso | Vi | 12/11/02 | 0 | 0 | 0 | 0 | N | K | main analyses |
| **Vi_02-06849_Kiribina** | 0.917 | SAMN27766212 | Burkina Faso | Vi | 12/11/02 | 0 | 0 | 0 | 0 | 0 | K | main analyses |
| **Vi_02-06861_Kiribina** | 0.926 | SAMN27766220 | Burkina Faso | Vi | 12/11/02 | 0 | 0 | 0 | 0 | 0 | Un | PCA |
| **GhaF264 (7)** | 0.882 | SAMN15546077 | Ghana | Jimiso Kakraba | 2004 | 0 | N | 2 | 2 | N | NA | PCA |
| **GhaF265 (7)** | 0.831 | SAMN15546078 | Ghana | Jimiso Kakraba | 2004 | 0 | N | 2 | 2 | N | NA | PCA |
| **Moz220 (7)** | 0.952 | SAMN15546079 | Mozambique | Chibuto | 2007 | 0 | N | 0 | 1 | N | NA | demographic models |
| **MozF123 (7)** | 0.874 | SAMN15546080 | Mozambique | Chibuto | 2007 | 0 | N | 0 | 1 | N | NA | demographic models |

| | | | | | | | | | | | | |
|---|---|---|---|---|---|---|---|---|---|---|---|---|
| **MozF260 (7)** | 0.916 | SAMN15546081 | Mozambique | Chibuto | 2007 | 0 | N | 0 | 1 | N | NA | demographic models |
| **MozF29 (7)** | 0.851 | SAMN15546082 | Mozambique | Chibuto | 2007 | 0 | N | 0 | 1 | N | NA | demographic models |
| **MozF35 (7)** | 0.801 | SAMN15546083 | Mozambique | Chibuto | 2007 | 0 | N | 0 | 1 | N | NA | demographic models |
| **MozF804 (7)** | 0.844 | SAMN15546084 | Mozambique | Chibuto | 2007 | 0 | N | 0 | 0 | N | NA | demographic models |
| **Ugf399 (7)** | 0.894 | SAMN15546076 | Uganda | Tororo Amoni | 2001 | 0 | N | 2 | 2 | N | NA | PCA |
| **Ugf401 (7)** | 0.896 | SAMN15546075 | Uganda | Tororo Amoni | 2001 | 0 | N | 2 | 2 | N | NA | PCA |
| **Ugf403 (7)** | 0.831 | SAMN15546074 | Uganda | Tororo Amoni | 2001 | 1 | N | 1 | 2 | N | NA | PCA |
| ***An. funestus-like* (7)** | 0.945 | SAMN15546033 | Malawi | Karonga | 2007 | | | | | | | Outgroup |
| ***An. longipalpis C* (7)** | 0.948 | SAMN15546038 | Zambia | Nyimba | 2011 | | | | | | | Outgroup |

[1] The character string comprising the SampleID includes the chromosomal form designations 'Folonzo' or 'Kiribina'. These designations are based on the original cytogenetic karyotyping data that was recorded, following the deterministic karyotype-based algorithm (46). In this study, we identified disagreements between cytogenetic definitions of chromosomal forms and PCA-based (genomic) definitions of ecotypes (e.g., La_02-07529_Folonzo and Vi_02-06855_Folonzo, which are assigned to the K ecotype). In addition, we have identified possible cytogenetic mis-readings or recording errors that were contradicted by computational karyotyping [sensu (6)], for example Ipendo_01-05791_Kiribina. We have nevertheless preserved the original designations in the SampleID for consistency with the original collection records.

[2] %Cov >10x: percentage of the genome with coverage exceeding 10x.

[3] Inversion genotype provided for 2Ra, 2Rs, 3Ra, 3Rb, and 3La was initially determined by cytogenetic karyotyping, but the genotypes in Table S1 have been validated, and if necessary corrected, based on computational karyotyping [sensu (6)]. Genotypes '0' and '2' denote standard or inverted homozygotes, respectively, while '1' denotes an inversion heterozygote. 'N' indicates that the karyotype was unreadable.

[4] Ecotype assignment is based on PCA of the X chromosome (Fig. 1A). Un, unassigned. NA, not applicable.

**Table S2.** ***An. funestus*** **specimens sequenced but not used in this study due to poor data quality.**

| SampleID | %Cov >10x | Biosample ID |
|---|---|---|
| **Ipendo_01-05793_Folonzo** | 0.248601 | SAMN27766106 |
| **La_02-07553_Folonzo** | 0.102298 | SAMN27766107 |
| **La_02-07687_Folonzo** | 0.16428 | SAMN27766108 |
| **La_02-07703_Folonzo** | 0.608235 | SAMN27766109 |
| **Pehele_02-06341_Folonzo** | 0.173455 | SAMN27766110 |
| **Pehele_02-06363_Folonzo** | 0.19684 | SAMN27766111 |
| **Sabtenga_02-07146_Folonzo** | 0.207235 | SAMN27766112 |
| **Sabtenga_02-07195_Folonzo** | 0.183476 | SAMN27766113 |
| **Vi_02-06673_Folonzo** | 0.392111 | SAMN27766114 |
| **Vi_02-06871_Folonzo** | 0.177583 | SAMN27766115 |
| **Dirze_02-06753_Kiribina** | 0.245866 | SAMN27766213 |
| **Dirze_02-06897_Kiribina** | 0.162061 | SAMN27766214 |
| **Dirze_02-07217_Kiribina** | 0.184455 | SAMN27766215 |
| **Dirze_02-07231_Kiribina** | 0.145669 | SAMN27766216 |
| **Kuiti_00-01518_Kiribina** | 0.134712 | SAMN27766217 |
| **Pehele_02-06339_Kiribina** | 0.160681 | SAMN27766218 |
| **Pehele_02-06365_Kiribina** | 0.153771 | SAMN27766219 |
| **Dirze_02-06753_Kiribina** | 0.245866 | SAMN27766213 |
| **Dirze_02-06897_Kiribina** | 0.162061 | SAMN27766214 |
| **Dirze_02-07217_Kiribina** | 0.184455 | SAMN27766215 |
| **Dirze_02-07231_Kiribina** | 0.145669 | SAMN27766216 |
| **Kuiti_00-01518_Kiribina** | 0.134712 | SAMN27766217 |
| **Pehele_02-06339_Kiribina** | 0.160681 | SAMN27766218 |
| **Pehele_02-06365_Kiribina** | 0.153771 | SAMN27766219 |

**Table S3. SNP counts by chromosome from the 168 *An. funestus* sampled from Burkina Faso[1].**

| Chromosome | SNPs |
|---|---|
| **X** | 2,160,904 |
| **2** | 12,878,731 |
| **3** | 9,243,658 |
| **Total** | 24,283,293 |

[1]Sequencing coverage averaged 38x reads per nucleotide. Of 160,265,210 bp of accessible sites in the *An. funestus* genome, a total of 114,270,430 bp of usable sites across all individuals remained after filtering (X, 10,148,005; chr 2, 58,111,028; chr 3, 46,011,397).

**Table S4. SNP counts by village and *An. funestus* ecotype.**

| | Ecotype K | | Ecotype F | |
|---|---|---|---|---|
| **Village** | **Sample Size** | **SNPs** | **Sample Size** | **SNPs** |
| **Bagre** | 15 | 5664774 | 0 | NA |
| **Dirze** | 11 | 4680874 | 0 | NA |
| **Ipendo** | 0 | NA | 6 | 3516083 |
| **Ividie** | 1 | 794699 | 1 | 825109 |
| **Koubri** | 17 | 6152454 | 17 | 7124062 |
| **Kuiti** | 15 | 5747290 | 19 | 7679127 |
| **La** | 1 | 796249 | 11 | 5298706 |
| **Pehele** | 9 | 4193753 | 11 | 5321981 |
| **Sabtenga** | 1 | 800873 | 13 | 5932778 |
| **Siby** | 1 | 79060 | 0 | NA |
| **Vi** | 3 | 2059769 | 14 | 6281467 |

**Table S5. Detailed ranges of priors from Stairway Plot 2 used for simulations in the ABC analyses. Time is in generations before sampling (2002).**

**A. Priors for effective population size (Ne)**

| Ecotype | Time (gens) | Ne ~U(L, H) |
|---|---|---|
| K | 0 | 50000, 300000 |
| K | 200 | 10000, 200000 |
| K | 400 | 50000, 200000 |
| K | 600 | 50000, 200000 |
| K | 1400 | 50000, 200000 |
| K | 3000 | 50000, 200000 |
| K | 6200 | 50000, 200000 |
| K | 10000 | 50000, 100000 |
| K | 13000 | 50000, 100000 |
| F | 0 | 300000, 5000000 |
| F | 400 | 300000, 2000000 |
| F | 600 | 50000, 300000 |
| F | 1000 | 300000, 2000000 |
| F | 1400 | 100000, 1000000 |
| F | 3000 | 50000, 500000 |
| F | 6200 | 50000, 200000 |
| F | 13000 | 50000, 100000 |
| F | 20000 | 10000, 100000 |
| F | 30000 | 10000, 100000 |
| F | 40000 | 10000, 100000 |
| F | 50000 | 10000, 100000 |
| F | 60000 | 10000, 100000 |
| F | 70000 | 10000, 100000 |

**B. Priors on migration rates and split times**

| Parameter | time (gens) | rate (proportion) |
|---|---|---|
| migration_1 | 0, 2999 | 0, 0.1 |
| migration_2 | 3000, 5000 | 0, 0.1 |
| migration_3 | 5001, 7000 | 0, 0.1 |
| migration_4 | 7001, 9500 | 0, 0 |
| split K/F | 7000, 14000 | NA |
| Split KF/Moz | 40000, 90000 | NA |

**C. Priors for variable recombination and mutation rates**

| Parameter | Prior Low | Prior High | Estimate |
|---|---|---|---|
| Recombination rate/gen | 1.64E-08 | 6.35E-08 | 4.1E-08 |
| Mutation rate/gen | 1.00E-09 | 6.1E-09 | 6.02E-09 |

**Table S6. Summary statistics used for demographic parameter inference**

| Name | Type | No. statistics (total) | Ref |
|---|---|---|---|
| Tajima's D (mean) | Per population | 1 (2) | (47) |
| Tajima's D (standard deviation) | Per population | 1 (2) | (47) |
| Nucleotide diversity (pi) (mean) | Per population | 1 (2) | (48) |
| Nucleotide diversity (pi) (standard deviation) | Per population | 1 (2) | (48) |
| Haplotype heterozygosity (mean) | Per population | 1 (2) | |
| Haplotype heterozygosity (standard deviation) | Per population | 1 (2) | |
| Truncated site frequency spectrum | Per population | 6 (12) | |
| Spatial site frequency spectrum (folded) | Per population | 34 (68) | (26) |
| Identity by state length (IBS) [11 quantiles, 4 subsample sizes] | Per population | 44 (88) | (49) |
| Allele frequency spectrum of IBS | Per population | 66 (132) | (50) |
| Allele frequency spectrum of IBS (standard deviation) | Per population | 66 (132) | (50) |
| $r^2$ (8 distance intervals; www.github.com/stsmall/abc_scripts2/ldintervals.txt) | Per population | 8 (16) | (49) |
| $F_{ST}$ (quantiles) | Pairwise | 5 | (30) |
| $d_{min}$ (quantiles) | Pairwise | 5 | (51) |
| $G_{min}$ (quantiles) | Pairwise | 5 | (52) |
| $d_{XY}$ (quantiles) | Pairwise | 5 | (48) |
| $IBS_{MaxXY}$ | Pairwise | 1 | (35) |
| $d_{d1}$ (quantiles) | Pairwise | 5 | (35) |
| $d_{d2}$ (quantiles) | Pairwise | 5 | (35) |
| $d_{d\text{-}Rank1}$ (quantiles) | Pairwise | 5 | (35) |
| $d_{d\text{-}Rank2}$ (quantiles) | Pairwise | 5 | (35) |
| $Z_X$ (quantiles) | Pairwise | 5 | (35) |
| joint site frequency spectrum (binned) | Pairwise | 23 | (53) |
| **Total** | | **529** | |

**Table S7. Confusion matrix for the five demographic models tested. Also shown are the misclassification proportions (Classification Error) and approximate posterior probability of the models.**

| | **Predicted classification** | | | | | | | | |
|---|---|---|---|---|---|---|---|---|---|
| | | **I** | **SC** | **IIM** | **IM** | **PM** | **Sum** | **Class Error** | **Posterior Probability** |
| **Actual classification** | **I** | 0.88 | 0.02 | 0.07 | 0.025 | 0.005 | **1.00** | **0.12** | **0.03** |
| | **SC** | 0.07 | 0.33 | 0.26 | 0.31 | 0.03 | **1.00** | **0.67** | **0.12** |
| | **IIM** | 0.04 | 0.15 | 0.45 | 0.35 | 0.01 | **1.00** | **0.55** | **0.44** |
| | **IM** | 0.01 | 0.31 | 0.17 | 0.42 | 0.09 | **1.00** | **0.58** | **0.40** |
| | **PM** | 0 | 0.02 | 0.01 | 0.05 | 0.92 | **1.00** | **0.08** | **0.01** |

I, Isolation model; SC, Secondary contact; IIM, Initial Isolation with Migration; IM, Isolation with Migration; PM, Panmictic; Class Error, classification error.

**Table S8. Comparison of parameters inferred for the IIM model by pg-GAN and ABC.** Parameter values correspond to K and F ecotype populations at the time of sampling (2002).

| | **Ne-K** | **Ne-F** | **Ne-Anc** | **Tsplit (generations)** | **Mig** | **Mutation rate per base, per generation** | **Recombination rate per base, per generation** |
|---|---|---|---|---|---|---|---|
| Pg-GAN | 69,831 | 3,256,961 | 44,670 | 14,013 | 0 | $6.01^{-09}$ | $4.10^{-08}$ |
| ABC median (95% CI) | 86,309 (45,763-128,348) | 2,376,519 (474,017-4,820,988) | 34,087 (11,951-87,295) | 12,365 (8,817-20,220) | 0 | $3.55^{-09}$ ($1.00^{-09}$-$6.10^{-09}$) | $3.99^{-08}$ ($1.64^{-08}$-$6.35^{-08}$) |

Anc, ancestral KF population; Tsplit, time of divergence between K and F ecotypes in generations; Mig, current migration rate between K and F.

**Table S9. $F_{ST}$ outlier windows**

| Block ID | chromosome | win_start | win_end | partition | Selected SNPs | FILET NoMig |
|---|---|---|---|---|---|---|
| **1** | 2 | 22150001 | 22160000 | colinear | Y | Y |
| **2** | 2 | 24760001 | 24770000 | colinear | Y | Y |
| | | 24770001 | 24780000 | colinear | Y | Y |
| **3** | 2 | 27840001 | 27850000 | inverted | N | Y |
| **4** | 2 | 30580001 | 30590000 | inverted | N | Y |
| **5** | 2 | 30600001 | 30610000 | inverted | Y | Y |
| **6** | 2 | 31850001 | 31860000 | inverted | Y | Y |
| **7** | 2 | 34470001 | 34480000 | colinear | Y | Y |
| | | 34480001 | 34490000 | colinear | Y | Y |
| | | 34490001 | 34500000 | colinear | Y | Y |
| | | 34500001 | 34510000 | colinear | Y | Y |
| | | 34510001 | 34520000 | colinear | N | Y |
| **8** | 2 | 36190001 | 36200000 | colinear | Y | Y |
| | | 36200001 | 36210000 | colinear | N | Y |
| **9** | 2 | 73470001 | 73480000 | colinear | Y | Y |
| **10** | 3 | 2090001 | 2100000 | inverted | Y | Y |
| **11** | 3 | 4620001 | 4630000 | inverted | Y | Y |
| **12** | 3 | 10790001 | 1080000 | inverted | Y | Y |
| **13** | 3 | 11500001 | 11510001 | colinear | N | N |
| **14** | 3 | 25140001 | 25150000 | inverted | Y | Y |
| | | 25150001 | 25160000 | inverted | Y | Y |
| | | 25160001 | 25170000 | inverted | Y | Y |
| | | 25180001 | 25190000 | inverted | Y | Y |
| | | 25190001 | 25200000 | inverted | Y | Y |
| **15** | 3 | 33560001 | 33570000 | colinear | Y | Y |
| **16** | 3 | 35520001 | 35530000 | colinear | Y | Y |
| | | 35530001 | 35540000 | colinear | Y | Y |
| | | 35540001 | 35550000 | colinear | Y | Y |
| | | 35550001 | 35560000 | colinear | N | Y |
| **17** | 3 | 35880001 | 35890000 | colinear | Y | Y |
| | | 35890001 | 35900000 | colinear | Y | Y |
| | | 35900001 | 35910000 | colinear | Y | Y |
| | | 35910001 | 35920000 | colinear | Y | Y |
| **18** | 3 | 36020001 | 36030000 | colinear | Y | Y |
| **19** | 3 | 36280001 | 36290000 | colinear | Y | Y |
| **20** | 3 | 36290001 | 36300000 | colinear | Y | Y |

| | | | | | | |
|---|---|---|---|---|---|---|
| **21** | 3 | 40660001 | 40670000 | colinear | Y | Y |
| **22** | 3 | 40720001 | 40730000 | colinear | N | Y |
| **23** | 3 | 43460001 | 43470000 | colinear | Y | Y |
| **24** | 3 | 43490001 | 43500000 | colinear | Y | Y |
| **25** | X | 3760001 | 3770000 | colinear | Y | Y |
| | | 3770001 | 3780000 | colinear | Y | Y |
| **26** | X | 13840001 | 13850000 | colinear | Y | Y |
| | | 13850001 | 13860000 | colinear | Y | Y |

**Table S10. Candidate SNPs and loci (differentially selected SNPs and genealogies)**

| chr | snp_pos | partition | tree_start | tree_end | pop | Mut age | impact | geneID | function |
|---|---|---|---|---|---|---|---|---|---|
| 3 | 40663176 | COL | 40663165 | 40663219 | F | 406042 | intron | AFUN000692 | dynein light chain Tctex1-like |
| 3 | 10790198 | INV | 10790190 | 10790301 | F | 561387 | 3_prime_UTR | AFUN002773 | PHD finger protein rhinoceros |
| 3 | 10792850 | INV | 10792847 | 10793005 | K | 356576 | missense | AFUN002773 | PHD finger protein rhinoceros |
| 3 | 10793208 | INV | 10793202 | 10793458 | K | 186452 | missense | AFUN002773 | PHD finger protein rhinoceros |
| 3 | 10797119 | INV | 10797094 | 10797174 | K | 586185 | missense | AFUN002773 | PHD finger protein rhinoceros |
| 3 | 10797120 | INV | 10797094 | 10797174 | K | 388774 | missense | AFUN002773 | PHD finger protein rhinoceros |
| 3 | 10798570 | INV | 10798526 | 10798640 | K | 13641 | synonymous | AFUN002773 | PHD finger protein rhinoceros |
| 3 | 10799062 | INV | 10799055 | 10799097 | K | 356576 | synonymous | AFUN002773 | PHD finger protein rhinoceros |
| 3 | 10799274 | INV | 10799254 | 10799280 | K | 388774 | missense | AFUN002773 | PHD finger protein rhinoceros |
| 2 | 30609917 | INV | 30609894.5 | 30610028.5 | K | 147001 | missense | AFUN003706 | unspecified |
| 2 | 30609918 | INV | 30609894.5 | 30610028.5 | K | 147001 | missense | AFUN003706 | unspecified |
| 2 | 30609950 | INV | 30609894.5 | 30610028.5 | K | 143858 | missense | AFUN003706 | unspecified |
| 3 | 35885210 | COL | 35885185 | 35885264 | F | 414816 | 5_prime_UTR | AFUN004278 | beta-hydroxy-delta-5-steroid dehydrogenase |
| 3 | 35885219 | COL | 35885185 | 35885264 | F | 327045 | 5_prime_UTR | AFUN004278 | beta-hydroxy-delta-5-steroid dehydrogenase |
| 3 | 35885690 | COL | 35885556 | 35885773 | F | 299960 | intron | AFUN004278 | beta-hydroxy-delta-5-steroid dehydrogenase |
| 3 | 35885891 | COL | 35885889 | 35885937 | F | 776346 | intron | AFUN004278 | beta-hydroxy-delta-5-steroid dehydrogenase |
| 3 | 35886365 | COL | 35886353 | 35886650 | F | 93370 | intron | AFUN004278 | beta-hydroxy-delta-5-steroid dehydrogenase |
| 3 | 35888125 | COL | 35888112 | 35888165 | F | 639116 | intron | AFUN004278 | beta-hydroxy-delta-5-steroid dehydrogenase |
| 3 | 35889799 | COL | 35889759 | 35889850 | F | 405947 | intron | AFUN004278 | beta-hydroxy-delta-5-steroid dehydrogenase |
| 3 | 35892624 | COL | 35892591 | 35892699 | F | 275118 | intron | AFUN004278 | beta-hydroxy-delta-5-steroid dehydrogenase |
| 3 | 35893594 | COL | 35893583 | 35893612 | F | 380462 | intron | AFUN004278 | beta-hydroxy-delta-5-steroid dehydrogenase |
| 3 | 35896837 | COL | 35896835 | 35897116 | F | 134826 | intron | AFUN004278 | beta-hydroxy-delta-5-steroid dehydrogenase |
| 3 | 35898711 | COL | 35898674 | 35898856 | F | 129123 | intron | AFUN004278 | beta-hydroxy-delta-5-steroid dehydrogenase |

| | | | | | | | | | |
|---|---|---|---|---|---|---|---|---|---|
| 3 | 35898859 | COL | 35898856 | 35899160 | F | 388774 | intron | AFUN004278 | beta-hydroxy-delta-5-steroid dehydrogenase |
| 3 | 35900083 | COL | 35900038 | 35900181 | F | 388774 | 3_prime_UTR | AFUN004278 | beta-hydroxy-delta-5-steroid dehydrogenase |
| 3 | 35900269 | COL | 35900181 | 35900360 | F | 246938 | 3_prime_UTR | AFUN004278 | beta-hydroxy-delta-5-steroid dehydrogenase |
| 3 | 35900605 | COL | 35900580 | 35900651 | F | 514894 | 3_prime_UTR | AFUN004278 | beta-hydroxy-delta-5-steroid dehydrogenase |
| 3 | 35900837 | COL | 35900651 | 35900864 | F | 433140 | 3_prime_UTR | AFUN004278 | beta-hydroxy-delta-5-steroid dehydrogenase |
| 3 | 35884882 | COL | 35884777 | 35884960 | K | 573652 | 5_prime_UTR | AFUN004278 | beta-hydroxy-delta-5-steroid dehydrogenase |
| 3 | 35884952 | COL | 35884777 | 35884960 | K | 598992 | 5_prime_UTR | AFUN004278 | beta-hydroxy-delta-5-steroid dehydrogenase |
| 3 | 35886033 | COL | 35886027 | 35886067 | K | 776346 | intron | AFUN004278 | beta-hydroxy-delta-5-steroid dehydrogenase |
| 3 | 35899447 | COL | 35899442 | 35899769 | K | 194688 | synonymous | AFUN004278 | beta-hydroxy-delta-5-steroid dehydrogenase |
| 3 | 35899531 | COL | 35899442 | 35899769 | K | 143858 | synonymous | AFUN004278 | beta-hydroxy-delta-5-steroid dehydrogenase |
| 3 | 25189229 | INV | 25189209 | 25189229 | K | 533382 | intron | AFUN004303 | putative tyramine receptor |
| 3 | 25192804 | INV | 25192794 | 25192936 | K | 667347 | intron | AFUN004303 | putative tyramine receptor |
| 3 | 25192874 | INV | 25192794 | 25192936 | K | 123689 | intron | AFUN004303 | putative tyramine receptor |
| 3 | 25193010 | INV | 25192936 | 25193099 | K | 180515 | intron | AFUN004303 | putative tyramine receptor |
| 3 | 25193077 | INV | 25192936 | 25193099 | K | 241658 | intron | AFUN004303 | putative tyramine receptor |
| 3 | 25193169 | INV | 25193122 | 25193290 | K | 330618 | intron | AFUN004303 | putative tyramine receptor |
| 3 | 25193366 | INV | 25193366 | 25193495 | K | 612078 | intron | AFUN004303 | putative tyramine receptor |
| 3 | 25193851 | INV | 25193777 | 25193948 | K | 306513 | missense | AFUN004303 | putative tyramine receptor |
| 2 | 34470981 | COL | 34470981 | 34470989 | F | 543511 | synonymous | AFUN005531 | diacylglycerol kinase (ATP dependent) |
| 2 | 34471136 | COL | 34471124 | 34471148 | F | 452273 | synonymous | AFUN005531 | diacylglycerol kinase (ATP dependent) |
| 2 | 34472199 | COL | 34472179.5 | 34472243 | F | 376570 | synonymous | AFUN005531 | diacylglycerol kinase (ATP dependent) |
| 2 | 34472729 | COL | 34472729 | 34472735 | K | 526143 | synonymous | AFUN005531 | diacylglycerol kinase (ATP dependent) |
| 2 | 34472530 | COL | 34472478.5 | 34472586.5 | K | 682086 | intron | AFUN005531 | diacylglycerol kinase (ATP dependent) |
| 2 | 34480166 | COL | 34480021 | 34480181 | F | 462586 | synonymous | AFUN005532 | pyridoxamine 5'-phosphate oxidase |
| 2 | 34481872 | COL | 34481872 | 34481879 | F | 653079 | 5_prime_UTR_premature_start_codon_gain | AFUN005533 | Arrestin_C domain-containing protein |

| | | | | | | | | | |
|---|---|---|---|---|---|---|---|---|---|
| 2 | 34482425 | COL | 34482251 | 34482442 | K | 653079 | upstream_gene | AFUN005533 | Arrestin_C domain-containing protein |
| X | 3775983 | COL | 3775982 | 3776078 | K | 632283 | upstream_gene | AFUN006243 | ZP domain-containing protein |
| X | 3779495 | COL | 3779488.5 | 3779507 | K | 612078 | downstream_gene | AFUN006243 | ZP domain-containing protein |
| 2 | 31858904 | INV | 31858738 | 31858907 | K | 573652 | upstream_gene | AFUN008518 | accessory gland protein Acp62F-like |
| 2 | 31858973 | INV | 31858907 | 31859043 | K | 503885 | missense | AFUN008518 | accessory gland protein Acp62F-like |
| 2 | 31859144 | INV | 31859061 | 31859196 | K | 632283 | missense | AFUN008518 | accessory gland protein Acp62F-like |
| 3 | 2091909 | INV | 2091908 | 2091941 | F | 423879 | missense | AFUN008640 | ANK_REP_REGION domain-containing protein |
| 3 | 2092775 | INV | 2092748 | 2092896 | F | 433140 | synonymous | AFUN008640 | ANK_REP_REGION domain-containing protein |
| 3 | 2098969 | INV | 2098965 | 2099021 | F | 252333 | missense | AFUN008640 | ANK_REP_REGION domain-containing protein |
| 3 | 2090591 | INV | 2090577 | 2090629 | K | 334191 | synonymous | AFUN008640 | ANK_REP_REGION domain-containing protein |
| 3 | 2090774 | INV | 2090756 | 2090883 | K | 299960 | synonymous | AFUN008640 | ANK_REP_REGION domain-containing protein |
| 3 | 2096120 | INV | 2096111 | 2096147 | K | 625451 | synonymous | AFUN008640 | ANK_REP_REGION domain-containing protein |
| 3 | 2096810 | INV | 2096737 | 2096819 | K | 681927 | synonymous | AFUN008640 | ANK_REP_REGION domain-containing protein |
| 3 | 2099759 | INV | 2099756 | 2099825 | K | 625451 | synonymous | AFUN008640 | ANK_REP_REGION domain-containing protein |
| 3 | 2099760 | INV | 2099756 | 2099825 | K | 612078 | missense | AFUN008640 | ANK_REP_REGION domain-containing protein |
| 3 | 33564172 | COL | 33564170 | 33564183 | K | 364367 | synonymous | AFUN008750 | unspecified |
| 3 | 33569558 | COL | 33569554 | 33570034 | K | 320053 | synonymous | AFUN014940 | organic cation transporter |
| 2 | 34492992 | COL | 34492992 | 34493184 | F | 503885 | intron | AFUN019981 | putative G-protein coupled receptor GPCR |
| 2 | 34500804 | COL | 34500647 | 34500837 | F | 487840 | intron | AFUN019981 | putative G-protein coupled receptor GPCR |
| 2 | 34500825 | COL | 34500647 | 34500837 | F | 646097 | intron | AFUN019981 | putative G-protein coupled receptor GPCR |
| 2 | 34505033 | COL | 34505019 | 34505218 | F | 393021 | intron | AFUN019981 | putative G-protein coupled receptor GPCR |
| X | 13847740 | COL | 13847549 | 13847765 | F | 467203 | intron | AFUN020012 | diacylglycerol kinase (ATP dependent) |
| X | 13849825 | COL | 13849597 | 13849954 | F | 586185 | intron | AFUN020012 | diacylglycerol kinase (ATP dependent) |
| X | 13849852 | COL | 13849597 | 13849954 | F | 598992 | intron | AFUN020012 | diacylglycerol kinase (ATP dependent) |
| X | 13849861 | COL | 13849597 | 13849954 | F | 618765 | intron | AFUN020012 | diacylglycerol kinase (ATP dependent) |
| X | 13849871 | COL | 13849597 | 13849954 | F | 660213 | intron | AFUN020012 | diacylglycerol kinase (ATP dependent) |

| | | | | | | | | | |
|---|---|---|---|---|---|---|---|---|---|
| X | 13849896 | COL | 13849597 | 13849954 | F | 573652 | intron | AFUN020012 | diacylglycerol kinase (ATP dependent) |
| X | 13840552 | COL | 13840495 | 13840599 | K | 598992 | intron | AFUN020012 | diacylglycerol kinase (ATP dependent) |
| X | 13841030 | COL | 13840922 | 13841030 | K | 178564 | intron | AFUN020012 | diacylglycerol kinase (ATP dependent) |
| X | 13842991 | COL | 13842960 | 13843101 | K | 776346 | intron | AFUN020012 | diacylglycerol kinase (ATP dependent) |
| X | 13843180 | COL | 13843101 | 13843303 | K | 231435 | intron | AFUN020012 | diacylglycerol kinase (ATP dependent) |
| X | 13843473 | COL | 13843303 | 13843629 | K | 275118 | intron | AFUN020012 | diacylglycerol kinase (ATP dependent) |
| X | 13844222 | COL | 13844149 | 13844323 | K | 356576 | 5_prime_UTR | AFUN020012 | diacylglycerol kinase (ATP dependent) |
| X | 13847287 | COL | 13847024 | 13847316 | K | 410574 | intron | AFUN020012 | diacylglycerol kinase (ATP dependent) |
| X | 13847473 | COL | 13847316 | 13847549 | K | 356576 | 5_prime_UTR | AFUN020012 | diacylglycerol kinase (ATP dependent) |
| X | 13847524 | COL | 13847316 | 13847549 | K | 372327 | 5_prime_UTR | AFUN020012 | diacylglycerol kinase (ATP dependent) |
| X | 13847665 | COL | 13847549 | 13847765 | K | 343488 | intron | AFUN020012 | diacylglycerol kinase (ATP dependent) |
| X | 13848349 | COL | 13848346 | 13848366 | K | 313210 | intron | AFUN020012 | diacylglycerol kinase (ATP dependent) |
| X | 13848565 | COL | 13848421 | 13848568 | K | 364367 | intron | AFUN020012 | diacylglycerol kinase (ATP dependent) |
| X | 13849179 | COL | 13848974 | 13849232 | K | 33811 | 5_prime_UTR | AFUN020012 | diacylglycerol kinase (ATP dependent) |
| X | 13849602 | COL | 13849597 | 13849954 | K | 219392 | 5_prime_UTR | AFUN020012 | diacylglycerol kinase (ATP dependent) |
| X | 13849610 | COL | 13849597 | 13849954 | K | 256879 | 5_prime_UTR | AFUN020012 | diacylglycerol kinase (ATP dependent) |
| X | 13849620 | COL | 13849597 | 13849954 | K | 190959 | 5_prime_UTR | AFUN020012 | diacylglycerol kinase (ATP dependent) |
| X | 13849624 | COL | 13849597 | 13849954 | K | 143992 | 5_prime_UTR | AFUN020012 | diacylglycerol kinase (ATP dependent) |
| X | 13849627 | COL | 13849597 | 13849954 | K | 327122 | 5_prime_UTR_premature_start_codon_gain | AFUN020012 | diacylglycerol kinase (ATP dependent) |
| X | 13849660 | COL | 13849597 | 13849954 | K | 146858 | 5_prime_UTR | AFUN020012 | diacylglycerol kinase (ATP dependent) |
| X | 13849799 | COL | 13849597 | 13849954 | K | 302671 | intron | AFUN020012 | diacylglycerol kinase (ATP dependent) |
| X | 13849888 | COL | 13849597 | 13849954 | K | 231435 | intron | AFUN020012 | diacylglycerol kinase (ATP dependent) |
| X | 13849921 | COL | 13849597 | 13849954 | K | 219275 | intron | AFUN020012 | diacylglycerol kinase (ATP dependent) |
| X | 13849948 | COL | 13849597 | 13849954 | K | 241658 | intron | AFUN020012 | diacylglycerol kinase (ATP dependent) |
| X | 13851327 | COL | 13851295 | 13851344 | K | 146263 | upstream_gene | AFUN020012 | diacylglycerol kinase (ATP dependent) |

| | | | | | | | | | |
|---|---|---|---|---|---|---|---|---|---|
| X | 13851734 | COL | 13851584 | 13851764 | K | 426357 | upstream_gene | AFUN020012 | diacylglycerol kinase (ATP dependent) |
| X | 13851807 | COL | 13851764 | 13851844 | K | 208506 | upstream_gene | AFUN020012 | diacylglycerol kinase (ATP dependent) |
| X | 13851818 | COL | 13851764 | 13851844 | K | 469387 | upstream_gene | AFUN020012 | diacylglycerol kinase (ATP dependent) |
| X | 13851835 | COL | 13851764 | 13851844 | K | 284199 | upstream_gene | AFUN020012 | diacylglycerol kinase (ATP dependent) |
| X | 13856703 | COL | 13856450 | 13856724 | K | 80261 | intron | AFUN020012 | diacylglycerol kinase (ATP dependent) |
| X | 13857323 | COL | 13856982 | 13857332 | K | 55583 | intron | AFUN020012 | diacylglycerol kinase (ATP dependent) |
| X | 13859456 | COL | 13859365 | 13859496 | K | 231435 | upstream_gene | AFUN020012 | diacylglycerol kinase (ATP dependent) |
| X | 13859476 | COL | 13859365 | 13859496 | K | 56797 | upstream_gene | AFUN020012 | diacylglycerol kinase (ATP dependent) |
| X | 13840613 | COL | 13840609.5 | 13840679 | K | 537638 | intron | AFUN020012-RA | diacylglycerol kinase (ATP dependent) |
| X | 13848073 | COL | 13847972 | 13848155 | K | 293547 | intron | AFUN020012-RA | diacylglycerol kinase (ATP dependent) |
| X | 13848811 | COL | 13848807.5 | 13848830 | K | 385696 | 5_prime_UTR | AFUN020012-RA | diacylglycerol kinase (ATP dependent) |
| X | 13849213 | COL | 13848972.5 | 13849228 | K | 310006 | 5_prime_UTR | AFUN020012-RA | diacylglycerol kinase (ATP dependent) |
| X | 13850009 | COL | 13849953.5 | 13850079.5 | K | 190704 | intron | AFUN020012-RA | diacylglycerol kinase (ATP dependent) |
| X | 13850021 | COL | 13849953.5 | 13850079.5 | K | 184459 | intron | AFUN020012-RA | diacylglycerol kinase (ATP dependent) |
| X | 13850030 | COL | 13849953.5 | 13850079.5 | K | 269236 | intron | AFUN020012-RA | diacylglycerol kinase (ATP dependent) |
| X | 13850035 | COL | 13849953.5 | 13850079.5 | K | 300591 | intron | AFUN020012-RA | diacylglycerol kinase (ATP dependent) |
| X | 13850036 | COL | 13849953.5 | 13850079.5 | K | 255208 | intron | AFUN020012-RA | diacylglycerol kinase (ATP dependent) |
| X | 13850052 | COL | 13849953.5 | 13850079.5 | K | 303237 | intron | AFUN020012-RA | diacylglycerol kinase (ATP dependent) |
| X | 13850053 | COL | 13849953.5 | 13850079.5 | K | 275696 | intron | AFUN020012-RA | diacylglycerol kinase (ATP dependent) |
| X | 13857368 | COL | 13857331 | 13857481 | K | 963645 | intron | AFUN020012-RC | diacylglycerol kinase (ATP dependent) |
| X | 13857404 | COL | 13857331 | 13857481 | K | 423879 | intron | AFUN020012-RC | diacylglycerol kinase (ATP dependent) |
| 3 | 36288290 | COL | 36288226 | 36288291 | F | 660213 | intron | AFUN020132 | RNA-binding protein Musashi |
| 3 | 36289857 | COL | 36289831 | 36289881 | F | 296892 | intron | AFUN020132 | RNA-binding protein Musashi |
| 3 | 36296552 | COL | 36296543 | 36296831.5 | F | 537638 | intron | AFUN020132 | RNA-binding protein Musashi |
| 3 | 36283776 | COL | 36283775 | 36283850 | K | 514894 | intron | AFUN020132 | RNA-binding protein Musashi |
| 3 | 36281230 | COL | 36281229.5 | 36281416 | K | 368519 | intron | AFUN020132 | RNA-binding protein Musashi |

| | | | | | | | | | |
|---|---|---|---|---|---|---|---|---|---|
| 3 | 36284130 | COL | 36284030 | 36284371 | K | 155243 | intron | AFUN020132 | RNA-binding protein Musashi |
| 3 | 36291594 | COL | 36291593 | 36291800 | K | 162101 | intron | AFUN020132 | RNA-binding protein Musashi |
| 3 | 36298204 | COL | 36298198.5 | 36298218.5 | K | 345222 | intron | AFUN020132 | RNA-binding protein Musashi |
| 3 | 35907484 | COL | 35907373 | 35907606 | F | 327045 | 5_prime_UTR_premature_start_codon_gain | AFUN020333 | unspecified |
| 3 | 35907595 | COL | 35907373 | 35907606 | F | 452273 | 5_prime_UTR_premature_start_codon_gain | AFUN020333 | unspecified |
| 3 | 35907600 | COL | 35907373 | 35907606 | F | 372327 | 5_prime_UTR | AFUN020333 | unspecified |
| 3 | 35907758 | COL | 35907606 | 35907901 | F | 356576 | synonymous | AFUN020333 | unspecified |
| 3 | 35909870 | COL | 35909681 | 35910141 | F | 397268 | intron | AFUN020333 | unspecified |
| 3 | 35910135 | COL | 35909681 | 35910141 | F | 269236 | intron | AFUN020333 | unspecified |
| 3 | 35910146 | COL | 35910141 | 35910538 | F | 186452 | intron | AFUN020333 | unspecified |
| 3 | 35911107 | COL | 35910635 | 35911196 | F | 182466 | intron | AFUN020333 | unspecified |
| 3 | 35912163 | COL | 35912024 | 35912172 | F | 348953 | intron | AFUN020333 | unspecified |
| 3 | 35912574 | COL | 35912213 | 35912688 | F | 163776 | intron | AFUN020333 | unspecified |
| 3 | 35911763 | COL | 35911756 | 35911942 | K | 341492 | intron | AFUN020333 | unspecified |
| 3 | 35911951 | COL | 35911950 | 35912024 | K | 143858 | intron | AFUN020333 | unspecified |
| 3 | 35912214 | COL | 35912213 | 35912688 | K | 313209 | intron | AFUN020333 | unspecified |
| 3 | 36026476 | COL | 36026476 | 36026570 | K | 632283 | intron | AFUN020333 | unspecified |
| 3 | 36026697 | COL | 36026697 | 36026944 | K | 498498 | intron | AFUN020333 | unspecified |
| 3 | 36026729 | COL | 36026697 | 36026944 | K | 515976 | intron | AFUN020333 | unspecified |
| 3 | 36028039 | COL | 36027981 | 36028155 | K | 467203 | intron | AFUN020333 | unspecified |
| 3 | 36028447 | COL | 36028364 | 36028553 | K | 514894 | intron | AFUN020333 | unspecified |
| 3 | 36028457 | COL | 36028364 | 36028553 | K | 498498 | intron | AFUN020333 | unspecified |
| 3 | 36028534 | COL | 36028364 | 36028553 | K | 509389 | intron | AFUN020333 | unspecified |
| X | 3769836 | COL | 3769822 | 3769903.5 | K | 667347 | missense | AFUN020681 | Glyco_18 domain-containing protein |

| | | | | | | | | | |
|---|---|---|---|---|---|---|---|---|---|
| X | 3768895 | COL | 3768668 | 3768895 | K | 579919 | missense | AFUN020681,AFUN006243 | Glyco_18 domain-containing protein |
| X | 3768907 | COL | 3768895 | 3769153 | K | 537638 | missense | AFUN020681,AFUN006243 | Glyco_18 domain-containing protein |
| X | 3768928 | COL | 3768895 | 3769153 | K | 579919 | missense | AFUN020681,AFUN006243 | Glyco_18 domain-containing protein |
| X | 3769153 | COL | 3768895 | 3769153 | K | 414913 | missense | AFUN020681,AFUN006243 | Glyco_18 domain-containing protein |
| X | 3770068 | COL | 3770007 | 3770095 | K | 555386 | missense | AFUN020681,AFUN006243 | Glyco_18 domain-containing protein |
| X | 3776981 | COL | 3776970 | 3776996 | K | 639116 | missense | AFUN021645,AFUN006243 | chitinase |
| 2 | 24769967 | COL | 24769908 | 24769980 | K | 468513 | missense | AFUN021979 | unspecified |
| 2 | 24769987 | COL | 24769980 | 24770165 | K | 472361 | missense | AFUN021979 | unspecified |
| 2 | 24770624 | COL | 24770165 | 24770864 | K | 605535 | missense | AFUN021979 | unspecified |
| 2 | 24770625 | COL | 24770165 | 24770864 | K | 605818 | missense | AFUN021979 | unspecified |
| 2 | 24770692 | COL | 24770165 | 24770864 | K | 592588 | upstream_gene | AFUN021979 | unspecified |
| 2 | 24770761 | COL | 24770165 | 24770864 | K | 574188 | missense | AFUN021979 | unspecified |
| 2 | 24770776 | COL | 24770165 | 24770864 | K | 646097 | missense | AFUN021979 | unspecified |
| 2 | 24770001 | COL | 24769980 | 24770165 | K | 442603 | missense | AFUN021979 | unspecified |
| 2 | 24772645 | COL | 24772491 | 24772734 | F | 1062136 | upstream_gene | AFUN021980 | unspecified |
| 2 | 24775145 | COL | 24774329 | 24775276 | K | 93370 | 5_prime_UTR | AFUN021980 | unspecified |
| 2 | 24765743 | COL | 24765729 | 24765778 | K | 334191 | 3_prime_UTR | AFUN021982 | unspecified |
| 3 | 4625340 | INV | 4625232 | 4625547 | K | 156848 | upstream_gene | AFUN022083 | unspecified |
| 3 | 4625425 | INV | 4625232 | 4625547 | K | 203288 | upstream_gene | AFUN022083 | unspecified |
| 3 | 4623804 | INV | 4623720 | 4623940 | K | 293547 | missense | AFUN022084 | unspecified |
| 3 | 4620585 | INV | 4620533 | 4620624 | K | 293547 | missense | AFUN022086 | unspecified |
| 3 | 4621840 | INV | 4621829 | 4621947 | K | 293547 | synonymous | AFUN022086 | unspecified |
| 3 | 4621930 | INV | 4621829 | 4621947 | K | 293547 | synonymous | AFUN022086 | unspecified |
| 2 | 24771794 | COL | 24771791 | 24772377 | F | 698129 | upstream_gene | intergenic | |

| | | | | | | | | |
|---|---|---|---|---|---|---|---|---|
| 2 | 24772438 | COL | 24772377 | 24772490.5 | F | 903147 | upstream_gene | intergenic |
| 2 | 24772449 | COL | 24772377 | 24772490.5 | F | 743503 | upstream_gene | intergenic |
| 2 | 24776025 | COL | 24776024 | 24776150 | F | 810639 | upstream_gene | intergenic |
| 2 | 24776781 | COL | 24776606 | 24776781 | F | 452273 | upstream_gene | intergenic |
| 2 | 24778820 | COL | 24778765 | 24778882 | F | 727776 | upstream_gene | intergenic |
| 2 | 24779080 | COL | 24779062 | 24779080 | F | 423879 | upstream_gene | intergenic |
| 2 | 31852056 | INV | 31852053.5 | 31852190.5 | F | 252333 | intergenic_region | intergenic |
| 2 | 31852630 | INV | 31852398.5 | 31852712 | F | 236491 | intergenic_region | intergenic |
| 2 | 34475330 | COL | 34475330 | 34475507 | F | 214587 | downstream_gene | intergenic |
| 2 | 34475507 | COL | 34475330 | 34475507 | F | 205509 | downstream_gene | intergenic |
| 2 | 34478800 | COL | 34478800 | 34478828 | F | 526143 | upstream_gene | intergenic |
| 2 | 34479128 | COL | 34479064 | 34479174 | F | 625451 | upstream_gene | intergenic |
| 2 | 34508730 | COL | 34508677 | 34508891 | F | 598992 | downstream_gene | intergenic |
| 2 | 34508891 | COL | 34508677 | 34508891 | F | 855694 | downstream_gene | intergenic |
| 2 | 34509103 | COL | 34509103 | 34509173 | F | 639265 | downstream_gene | intergenic |
| 2 | 34509238 | COL | 34509236 | 34509256.5 | F | 493112 | downstream_gene | intergenic |
| 2 | 36190456 | COL | 36190456 | 36190465 | F | 696826 | intergenic_region | intergenic |
| 2 | 36193386 | COL | 36193337 | 36193654 | F | 207730 | downstream_gene | intergenic |
| 2 | 73470864 | COL | 73470864 | 73471122 | F(K) | 414816 | intergenic_region | intergenic |
| 2 | 73475351 | COL | 73475351 | 73475514 | F | 573652 | intergenic_region | intergenic |
| 3 | 25149575 | INV | 25149574.5 | 25149637 | F | 472692 | intergenic_region | intergenic |
| 3 | 25149609 | INV | 25149574.5 | 25149637 | F | 467203 | intergenic_region | intergenic |
| 3 | 25149627 | INV | 25149574.5 | 25149637 | F | 493227 | intergenic_region | intergenic |
| 3 | 25167167 | INV | 25167165.5 | 25167210 | F | 579919 | intergenic_region | intergenic |
| 3 | 25197512 | INV | 25197452.5 | 25197627 | F | 327351 | downstream_gene | intergenic |
| 3 | 33568260 | COL | 33568211 | 33568346 | F | 549384 | downstream_gene | intergenic |

| | | | | | | | | |
|---|---|---|---|---|---|---|---|---|
| 3 | 33568311 | COL | 33568211 | 33568346 | F | 493112 | downstream_gene | intergenic |
| 3 | 33568334 | COL | 33568211 | 33568346 | F | 625451 | downstream_gene | intergenic |
| 3 | 35520715 | COL | 35520593 | 35520716 | F | 384618 | upstream_gene | intergenic |
| 3 | 35524010 | COL | 35523920 | 35524011 | F | 618765 | upstream_gene | intergenic |
| 3 | 35525096 | COL | 35525094 | 35525302.5 | F | 447438 | intergenic_region | intergenic |
| 3 | 35527697 | COL | 35527552 | 35527856.5 | F | 498498 | intergenic_region | intergenic |
| 3 | 35529760 | COL | 35529596.5 | 35529811 | F | 317075 | intergenic_region | intergenic |
| 3 | 35530510 | COL | 35530493.5 | 35530535.5 | F | 327045 | intergenic_region | intergenic |
| 3 | 35534431 | COL | 35534322 | 35534431 | F | 561518 | downstream_gene | intergenic |
| 3 | 35534887 | COL | 35534852 | 35535040 | F | 246938 | downstream_gene | intergenic |
| 3 | 35543885 | COL | 35543883.5 | 35544032.5 | F | 410382 | intergenic_region | intergenic |
| 3 | 35901779 | COL | 35901761 | 35901816 | F | 226487 | upstream_gene | intergenic |
| 3 | 35905089 | COL | 35904971 | 35905113 | F | 182466 | upstream_gene | intergenic |
| 3 | 35905766 | COL | 35905762 | 35905780 | F | 334191 | upstream_gene | intergenic |
| 3 | 35906214 | COL | 35906109 | 35906303 | F | 275118 | upstream_gene | intergenic |
| 3 | 43460643 | COL | 43459919 | 43460901.5 | F | 543511 | downstream_gene | intergenic |
| 3 | 43460790 | COL | 43459919 | 43460901.5 | F | 388774 | downstream_gene | intergenic |
| 2 | 22154412 | COL | 22154404 | 22154642 | K | 573652 | intergenic_region | intergenic |
| 2 | 34476681 | COL | 34476362 | 34477190 | K | 681927 | downstream_gene | intergenic |
| 2 | 34476683 | COL | 34476362 | 34477190 | K | 681927 | downstream_gene | intergenic |
| 2 | 34480528 | COL | 34480517 | 34480531 | K | 110994 | upstream_gene | intergenic |
| 2 | 36190150 | COL | 36190150 | 36190345 | K | 306513 | intergenic_region | intergenic |
| 2 | 36190430 | COL | 36190359 | 36190433 | K | 653079 | intergenic_region | intergenic |
| 2 | 36191475 | COL | 36191384 | 36191607 | K | 727606 | intergenic_region | intergenic |
| 2 | 36193311 | COL | 36192902 | 36193339 | K | 182466 | downstream_gene | intergenic |
| 2 | 73470864 | COL | 73470864 | 73471122 | K(F) | 414816 | intergenic_region | intergenic |

| | | | | | | | | |
|---|---|---|---|---|---|---|---|---|
| 3 | 4622583 | INV | 4622419 | 4622627 | K | 163776 | upstream_gene | intergenic |
| 3 | 4628694 | INV | 4628688 | 4628804 | K | 115897 | downstream_gene | intergenic |
| 3 | 25143957 | INV | 25143802 | 25143992 | K | 504003 | intergenic_region | intergenic |
| 3 | 25146445 | INV | 25146442 | 25146736 | K | 625451 | intergenic_region | intergenic |
| 3 | 25158357 | INV | 25158191 | 25158357 | K | 397268 | intergenic_region | intergenic |
| 3 | 25159506 | INV | 25159476 | 25159506 | K | 203288 | intergenic_region | intergenic |
| 3 | 25167936 | INV | 25167891 | 25167976 | K | 586322 | intergenic_region | intergenic |
| 3 | 25197464 | INV | 25197464 | 25197637 | K | 503885 | downstream_gene | intergenic |
| 3 | 25198158 | INV | 25198158 | 25198254 | K | 543511 | downstream_gene | intergenic |
| 3 | 25198314 | INV | 25198314 | 25198407 | K | 639116 | downstream_gene | intergenic |
| 3 | 25198712 | INV | 25198648 | 25198732 | K | 388865 | downstream_gene | intergenic |
| 3 | 25198729 | INV | 25198648 | 25198732 | K | 360472 | downstream_gene | intergenic |
| 3 | 33565019 | COL | 33564970 | 33565090 | K | 263479 | downstream_gene | intergenic |
| 3 | 33565052 | COL | 33564970 | 33565090 | K | 113419 | downstream_gene | intergenic |
| 3 | 33566268 | COL | 33566251 | 33566365 | K | 287270 | downstream_gene | intergenic |
| 3 | 35521121 | COL | 35521094 | 35521225 | K | 313210 | upstream_gene | intergenic |
| 3 | 35529215 | COL | 35529148 | 35529311 | K | 632283 | intergenic_region | intergenic |
| 3 | 35532237 | COL | 35532237 | 35532358 | K | 592865 | downstream_gene | intergenic |
| 3 | 35537960 | COL | 35537854 | 35538043 | K | 953343 | upstream_gene | intergenic |
| 3 | 35538043 | COL | 35537854 | 35538043 | K | 704440 | upstream_gene | intergenic |
| 3 | 35538177 | COL | 35538172 | 35538230 | K | 768046 | upstream_gene | intergenic |
| 3 | 35538442 | COL | 35538353 | 35538670 | K | 531891 | upstream_gene | intergenic |
| 3 | 40669151 | COL | 40669091 | 40669164 | K | 203288 | upstream_gene | intergenic |
| 3 | 43469015 | COL | 43469010 | 43469341 | K | 167354 | intergenic_region | intergenic |
| X | 3763412 | COL | 3763264 | 3763412 | K | 433140 | downstream_gene | intergenic |
| 2 | 24768486 | COL | 24768404 | 24768527 | K | 59306 | upstream_gene | intergenic |

| | | | | | | | | |
|---|---|---|---|---|---|---|---|---|
| 2 | 36192792 | COL | 36192790 | 36192837 | K | 592588 | downstream_gene | intergenic |
| 3 | 25142719 | INV | 25142420.5 | 25142765.5 | K | 246996 | intergenic_region | intergenic |
| 3 | 25149403 | INV | 25149395.5 | 25149496 | K | 337841 | intergenic_region | intergenic |
| 3 | 25159963 | INV | 25159919.5 | 25160076 | K | 278123 | intergenic_region | intergenic |
| 3 | 25160403 | INV | 25160401.5 | 25160646.5 | K | 82015 | intergenic_region | intergenic |
| 3 | 25160644 | INV | 25160401.5 | 25160646.5 | K | 212318 | intergenic_region | intergenic |
| 3 | 25181881 | INV | 25181828.5 | 25181889.5 | K | 287270 | intergenic_region | intergenic |
| 3 | 25182135 | INV | 25182134 | 25182271.5 | K | 498498 | intergenic_region | intergenic |
| 3 | 35524272 | COL | 35524270.5 | 35524276.5 | K | 653231 | upstream_gene | intergenic |
| 3 | 35524449 | COL | 35524438 | 35524469.5 | K | 555386 | upstream_gene | intergenic |
| 3 | 43492854 | COL | 43492827 | 43493308 | K | 579919 | intergenic_region | intergenic |

**Table S11. Confusion matrix for the FILET no-migration classifier**

| | | **Predicted** | | |
|---|---|---|---|---|
| **True** | K ↔ F | noMig | mig12 | mig21 |
| | noMig | 1.000 | 0.000 | 0.000 |
| | mig12 | 0.003 | 0.992 | 0.005 |
| | mig21 | 0.000 | 0.001 | 0.999 |

**Table S12. Long single-molecule assembly of the *An. funestus* rDNA gene[1]**

| rDNA subunit | Length (bp) | Status |
|---|---:|---|
| IGS | 4,720 | Partial, 3'-end |
| 18S | 2,004 | Complete |
| ITS1 | 1,821 | Complete |
| 5.8S | 446 | Complete |
| ITS2 | 421 | Complete |
| 28S | 4,179 | Complete |
| IGS | 1,114 | Partial, 5'-end |
| Total | 14,705 | |

[1]GenBank accession TBD.
IGS, intergenic spacer; ITS1 and ITS2, internal transcribed spacer 1 and 2.
